# Leptin Receptor^+^ cells create a perisinusoidal niche for thrombopoiesis in the bone marrow by synthesizing CXCL14

**DOI:** 10.64898/2026.01.27.702029

**Authors:** Yuanyuan Xue, Ruixuan Zhang, Maowu Luo, Amanda R. Reyes, Salma Merchant, Trevor S. Tippetts, Gerik Grabowski, Tai Ngo, Yanan Sophia Zhang, Zhiguo Shang, Nisi Jiang, Nathaniel S. Tjahjono, Elise C. Jeffery, Yafeng Li, Tao Wei, Wen Gu, Liming Du, Ralph J. DeBerardinis, Kevin M. Dean, David W. Schmidtke, Thomas P. Mathews, Daniel Lucas, Zhiyu Zhao, Sean J. Morrison

## Abstract

Leptin Receptor-expressing (LepR^+^) stromal cells in the bone marrow are a critical source of growth factors for the maintenance of hematopoietic stem cells (HSCs) and most restricted hematopoietic progenitors. An important unresolved question is whether they also regulate terminal differentiation in some hematopoietic cells. We found that bone marrow LepR^+^ cells promote thrombopoiesis by synthesizing the chemokine CXCL14. *Cxcl14*-expressing LepR^+^ cells extend fine processes that wrap around perisinusoidal megakaryocytes. Deletion of *Cxcl14* from LepR^+^ cells did not significantly alter HSC function or most aspects of bone marrow hematopoiesis, including megakaryocyte generation; however, it significantly reduced the numbers of proplatelet-forming megakaryocytes in the bone marrow and platelets in the blood, as well as platelet quality. Megakaryocytes from *Lepr-cre; Cxcl14 ^fl/fl^* mice expressed lower levels of genes that encode granule proteins as compared to control mice, leading to the release of platelets with fewer α-granules and defective thrombus formation/growth. CXCL14 promoted platelet formation by remodeling lipid metabolism in megakaryocytes, increasing fatty acid transporter expression and enabling megakaryocytes to take up more polyunsaturated fatty acids from circulation. A high fat diet rescued the formation of proplatelet-forming megakaryocytes and platelets, as well as platelet function, in *Lepr-cre; Cxcl14 ^fl/fl^* mice. CXCL14 protein was sufficient to promote platelet formation by megakaryocytes in vitro and in vivo. LepR^+^ cells thus create a perisinusoidal niche for thrombopoiesis by producing CXCL14, which regulates lipid metabolism and terminal differentiation in megakaryocytes.

**Significance:** To our knowledge, this study presents the first evidence that LepR^+^ stromal cells in the bone marrow regulate terminal differentiation or metabolic function in hematopoietic cells. LepR^+^ cells regulate the maturation of megakaryocytes into proplatelet-forming megakaryocytes and platelet production in the bone marrow as well as platelet quality: platelets produced in the absence of CXCL14 from LepR^+^ cells are depleted of α-granules required for thrombus formation/growth. These findings thus identify a niche for thrombopoiesis in the bone marrow, a mechanism by which LepR^+^ cells regulate thrombopoiesis, and a new function for CXCL14 in the regulation of bone marrow thrombopoiesis.

## INTRODUCTION

Leptin Receptor-expressing (LepR^+^) stromal cells promote the maintenance of hematopoietic stem cells (HSCs) and most restricted hematopoietic progenitors in the bone marrow by producing Stem cell factor (SCF) (1–3), CXCL12 (4, 5), Pleiotrophin (6), Interleukin 7 (IL7) (7), and Colony stimulating factor 1 (CSF1) (8–10). LepR^+^ cells have also been described as CXCL12-abundant reticular (CAR) cells (11, 12). Some LepR^+^ cells are perisinusoidal, where they create niches for HSCs, erythroid progenitors, and other restricted hematopoietic progenitors (1, 2, 4, 13–15). Other LepR^+^ cells are peri-arteriolar, where they create niches for lymphoid progenitors (16). LepR^+^ cells also promote vascular regeneration by producing Angiopoietin-1 (17) and VEGF-C (18) as well as peripheral nerve maintenance in bone marrow by producing nerve growth factor (19). However, it is unknown whether LepR^+^ cells regulate terminal differentiation in hematopoietic cells.

Platelets regulate hemostasis, inflammation, and tissue repair by enabling blood clotting (20). Platelets are produced in the bone marrow by megakaryocytes (21, 22), which localize to sinusoidal blood vessels (23, 24) and release platelets into circulation (25). A number of cytokines promote the formation of megakaryocytes in the bone marrow including thrombopoietin (26–28), SCF (29–31), leukemia inhibitory factor (32), fibroblast growth factor 4 (33, 34), IL-3, IL-6 and IL-11 (29, 35–38). Thrombopoietin is produced by hepatocytes in the liver (39) and SCF is produced by LepR^+^ cells in the bone marrow (1) but the cellular sources of other factors are unknown.

Several other cytokines and chemokines promote the maturation of megakaryocytes into proplatelet-forming megakaryocytes or the generation of platelets including Insulin growth factor 1 (40), IL-1a (41), chemokine ligand 5 (42), and Interferon alpha (IFNα) (43). IFNα is produced by plasmacytoid dendritic cells in the bone marrow (43) but to our knowledge the cellular sources of other cytokines and chemokines are unknown.

CXCL14 is a chemokine that is expressed in multiple tissues (44), including in bone marrow stromal cells (45). The receptor for CXCL14 is uncertain: CXCR4, which is widely expressed by hematopoietic cells, can serve as a receptor for CXCL14 (46, 47) but other receptors have been suggested as well (48–52). CXCL14 promotes immune cell migration to sites of inflammation (53, 54) as well as platelet migration and thrombus consolidation (47). However, CXCL14 has not been shown to play any role in the regulation of hematopoiesis or thrombopoiesis in the bone marrow.

## RESULTS

### *Cxcl14* is mainly expressed by LepR^+^ cells in the bone marrow

To assess the *Cxcl14* expression pattern in the bone marrow, we reanalyzed published single cell RNA sequencing data (55). *Cxcl14* was expressed by LepR^+^ cells but rarely by osteoblasts, chondrocytes, fibroblasts, endothelial cells, pericytes, or Schwann cells (Fig. 1A). Quantitative reverse transcription PCR (qRT-PCR) confirmed *Cxcl14* was expressed by LepR^+^CD45^−^Ter119^−^CD31^−^ stromal cells but not by endothelial cells, HSCs or WBM cells (Fig. 1B). *Cxcl14* was expressed by both *Osteolectin* negative *Lepr*^+^ cells, which are perisinusoidal (1), and *Osteolectin*^+^*Lepr*^+^ cells, which are peri-arteriolar (16) (Fig. 1A).

**Figure 1:**
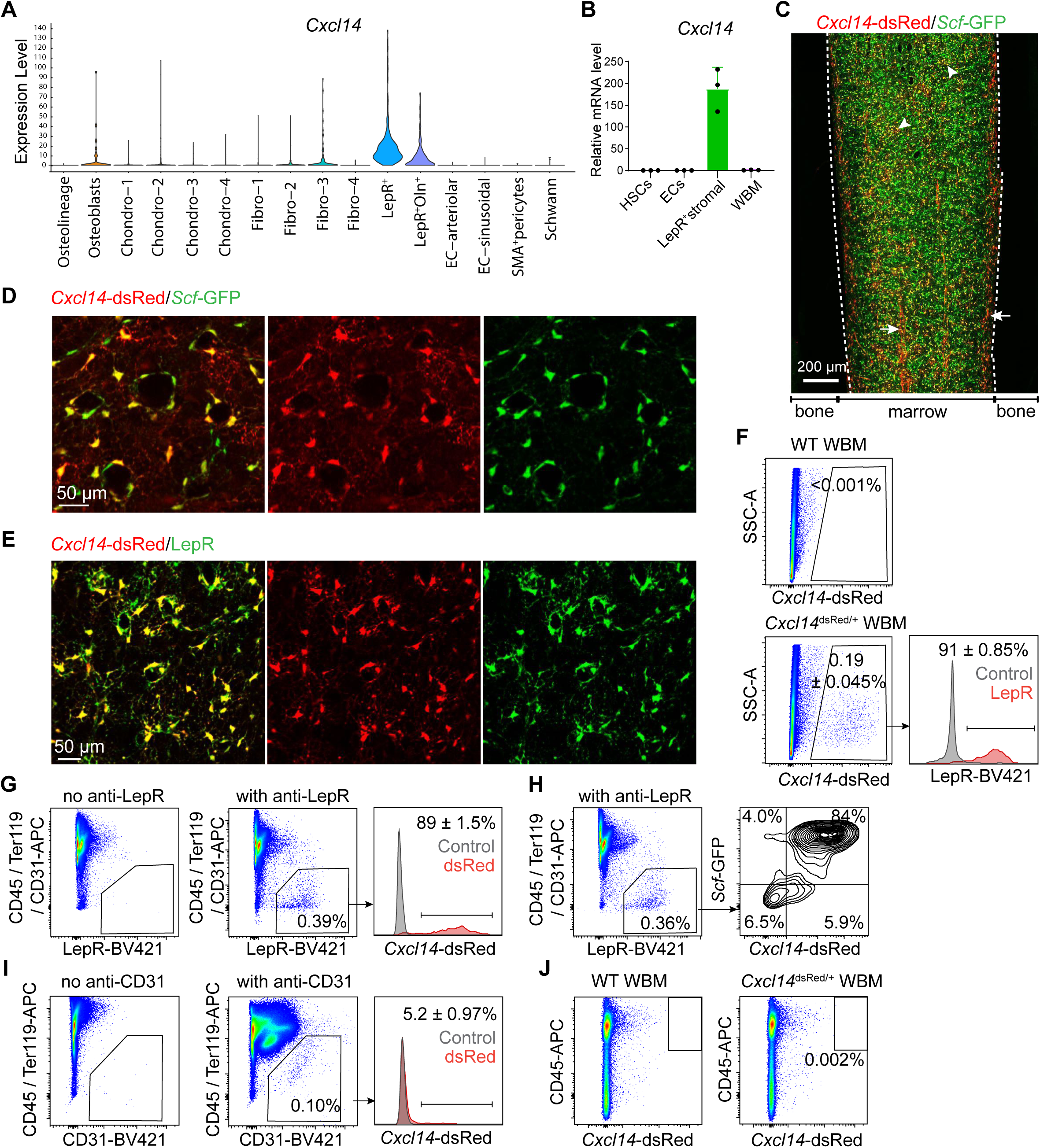
LepR^+^ stromal cells were the main source of *Cxcl14* in the bone marrow. (A) Single cell RNA sequencing (55) showing *Cxcl14* expression by LepR^+^ stromal cells from enzymatically dissociated bone and bone marrow (a total of 3-4 mice from 3 independent experiments). (B) *Cxcl14* expression by quantitative reverse-transcription PCR in CD150^+^CD48^-^ Lineage^-^Sca1^+^ckit^+^ HSCs, CD31^+^CD45^−^Ter119^−^ endothelial cells, LepR^+^CD45^−^Ter119^−^CD31^−^ stromal cells, and unfractionated cells from the bone marrow of two month-old mice (a total of 3 samples per cell population from 3 independent experiments; each dot represents a different sample). (C and D) Deep confocal imaging of femur bone marrow from adult *Cxcl14 ^dsRed^Scf ^GFP^* mice: *Cxcl14*-dsRed^+^ cells localized around sinusoids (arrowheads) and arterioles (arrow) throughout bone marrow (white dashed lines indicate the interface of bone marrow and bone; see also the low magnification image of a whole bone in *SI Appendix,* Fig. S1C). Most *Cxcl14*- dsRed^+^ cells were *Scf*-GFP^+^ (images are representative of 3 mice). (E) Femur bone marrow from adult *Cxcl14 ^dsRed^* mice stained with anti-LepR antibody showing that most *Cxcl14*-dsRed^+^ cells were LepR^+^ (images are representative of 3 mice). (F-J) Flow cytometric analysis of enzymatically dissociated bone marrow from *Cxcl14 ^dsRed/+^*or *Cxcl14 ^dsRed^Scf ^GFP^* mice. Only 0.19±0.04% of bone marrow cells from *Cxcl14 ^dsRed/+^* mice were *Cxcl14*-dsRed^+^ and, of these, 91% were LepR^+^ (F). Conversely, 89% of LepR^+^ cells were *Cxcl14*-dsRed^+^ (G). 84% of LepR^+^ cells from bone marrow of *Cxcl14 ^dsRed^Scf ^GFP^* mice were *Cxcl14*-dsRed^+^ and *Scf*-GFP^+^ (H). Few CD31^+^CD45^−^Ter119^−^ endothelial cells from bone marrow of *Cxcl14 ^dsRed/+^*mice were *Cxcl14*- dsRed^+^ (I; 3-6 mice from 3 independent experiments). (J) Flow cytometric analysis of non-enzymatically dissociated bone marrow shows few or no CD45^+^ cells expressed *Cxcl14*-dsRed (3-6 mice from 3 independent experiments; see also *SI Appendix,* Fig. S1D). Data represent mean ± standard deviation in panel B.

We generated a *Cxcl14*-dsRed (*Cxcl14* ^dsRed^) knock-in reporter allele (*SI Appendix,* Fig. S1A-C). Confocal imaging of femurs from adult *Cxcl14* ^dsRed/+^*; Scf* ^GFP/+^ mice showed *Cxcl14* ^dsRed^ expression by perisinusoidal and peri-arteriolar stromal cells (Fig. 1C-E; *SI Appendix,* Fig. S1C). The *Cxcl14* ^dsRed^-expressing perisinusoidal stromal cells were *Scf-*GFP^+^ (Fig. 1D and 1H) and LepR^+^ (Fig. 1E).

Flow cytometric analysis of enzymatically dissociated bone marrow cells from adult *Cxcl14* ^dsRed/+^*; Scf* ^GFP/+^ mice showed that *Cxcl14* ^dsRed^ was expressed by 0.19±0.045% of bone marrow cells, 91±0.85% of which were LepR^+^ (Fig. 1F). Furthermore, 89±1.5% of LepR^+^ cells were *Cxcl14* ^dsRed^ positive (Fig. 1G). We rarely detected *Cxcl14* ^dsRed^ expression by endothelial cells (Fig. 1I), hematopoietic cells in bone marrow, megakarocytes (Fig. 1J; *SI Appendix,* Fig. S1D), or platelets in the blood (*SI Appendix,* Fig. S1E). The flow cytometry gates used to sort LepR^+^ cells and endothelial cells are in Fig. 1G-I and the gates for hematopoietic cells are in *SI Appendix,* Fig. S1F-I. Reanalysis of published single cell RNA sequencing data (55, 56) showed LepR^+^ cells expressed the highest levels of *Cxcl14* in the bone marrow at all ages, including postnatal day 4 (P4), P14 (*SI Appendix,* Fig. S2A) and 2, 12, and 24 months of age (*SI Appendix,* Fig. S2B). LepR^+^ cells thus appear to be the major source of *Cxcl14* in the bone marrow throughout life.

### CXCL14 is required for thrombopoiesis but not megakaryopoiesis

To test if CXCL14 regulates hematopoiesis, we generated mice with a floxed *Cxcl14* allele by inserting loxp sites on either side of exon 2 (*SI Appendix,* Fig. S3A-B). Cre mediated recombination would be predicted to cause a frameshift, yielding a strong loss of CXCL14 function. *Lepr-cre; Cxcl14 ^fl/fl^*mice were born in expected numbers and did not significantly differ from littermate controls in terms of gross appearance (*SI Appendix,* Fig. S3C) or body mass (*SI Appendix,* Fig. S3D). LepR^+^ cells had substantially lower levels of *Cxcl14* transcripts in 4 month-old *Lepr-cre; Cxcl14 ^fl/fl^*mice as compared to littermate controls (approximately 30% of control levels; *SI Appendix,* Fig. S3E). ELISA analysis showed that bone marrow serum had 73% lower levels of CXCL14 in 4 month-old *Lepr-cre; Cxcl14^fl/fl^* mice as compared to littermate controls but these mice did not differ in terms of CXCL14 levels in blood plasma (Fig. 2A). LepR^+^ cells are thus the major source of CXCL14 in the bone marrow but are not a significant source of CXCL14 in the blood.

**Figure 2:**
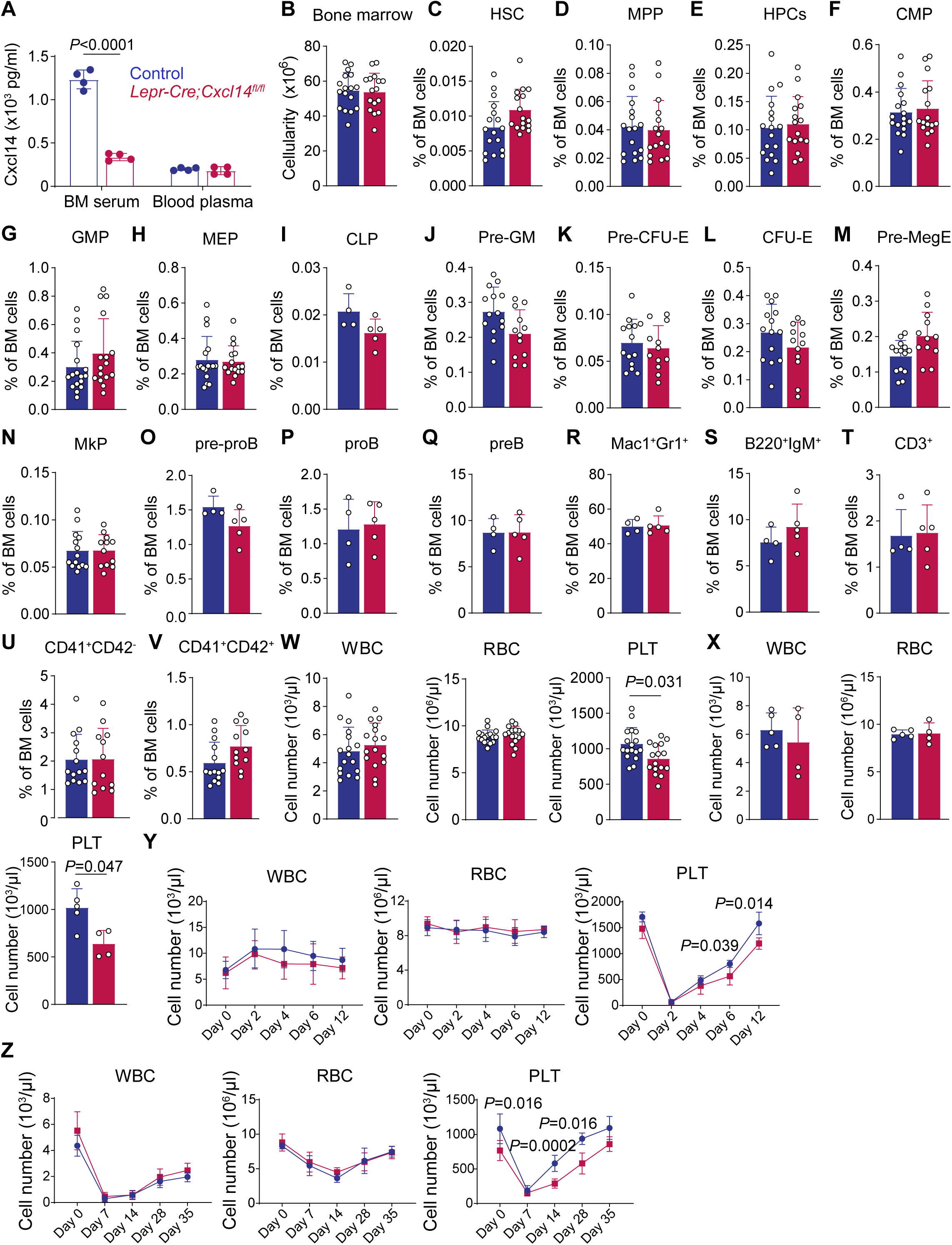
*Lepr-cre; Cxcl14^fl/fl^* mice exhibited grossly normal hematopoiesis but reduced platelet counts. (A) ELISA of CXCL14 levels in bone marrow serum and blood plasma from *Lepr-cre; Cxcl14^fl/fl^* and control mice (a total 4 mice per group from 2 independent experiments). (B-V) Bone marrow cellularity (B) and the frequencies of HSCs (C), MPPs (D), HPCs (E), CMPs (F), GMPs (G), MEPs (H), CLPs (I), Pre-GMs (J), Pre-CFUEs (K), CFU-Es (L), Pre-MegEs (M), megakaryocyte progenitors (N), Pre-proB cells (O), ProB cells (P), PreB cells (Q), Mac1^+^Gr1^+^ myeloid cells (R), B220^+^IgM^+^ B cells (S), CD3^+^ T cells (T), CD41^+^CD42d^-^ megakaryocytes (U) and CD41^+^CD42d^+^ megakaryocytes (V) in the bone marrow of *Lepr-cre; Cxcl14 ^fl/fl^* and littermate control mice (a total of 4-17 mice per genotype from 3-7 independent experiments). (W) White blood cell (WBC), red blood cell (RBC) and platelet (PLT) counts in the blood of *Lepr-cre; Cxcl14^fl/fl^* mice and littermate controls at 4 months of age (W; a total of 16-17 mice per genotype from 7 independent experiments) and 7 months of age (X; a total of 4-5 mice per genotype from 3 independent experiments). (Y) WBC, RBC and PLT counts from the blood of *Lepr-cre; Cxcl14^fl/fl^* and control mice before (day 0) and 2, 4, 6 and 12 days after treatment with anti-CD42b antibody to deplete platelets (a total of 6 mice per genotype from 2 independent experiments). (Z) WBC, RBC and PLT counts from *Lepr-cre; Cxcl14^fl/fl^*and control mice before (day 0) and 7, 14, 28 and 35 days after sublethal irradiation (a total of 4-8 mice per genotype from 2 independent experiments). All data represent mean ± standard deviation and each dot represents a different mouse in panels A-X. All statistical tests were two-sided. Statistical significance was assessed using a matched samples two-way ANOVA followed by Sidak’s multiple comparisons adjustment (A), a Student’s *t*-test (B), or Mann-Whitney tests (C-V) or Students’ *t*-tests (W-Z) followed by Holm-Sidak’s multiple comparisons adjustments.

By flow cytometric analysis, four month-old *Lepr-cre; Cxcl14^fl/fl^*and littermate control mice did not significantly differ in terms of bone marrow cellularity (Fig. 2B) or the frequencies of HSCs (Fig. 2C), MPPs (Fig. 2D), restricted hematopoietic progenitors (Fig. 2E-Q), or differentiated myeloid cells, B cells, T cells, or megakaryocytes (Fig. 2R-V) in the bone marrow. They also did not significantly differ in terms of spleen cellularity (*SI Appendix,* Fig. S3F) or the frequencies of HSCs, MPPs, restricted progenitors, or differentiated hematopoietic cells in the spleen (*SI Appendix,* Fig. S3G-Y). Finally, *Lepr-cre; Cxcl14^fl/fl^*and littermate control mice did not significantly differ in the reconstituting potential of bone marrow cells upon competitive transplantation into irradiated mice (*SI Appendix,* Fig. S3Z) or the frequencies of bone marrow or spleen cells that formed colonies in culture, including megakaryocyte colonies (*SI Appendix,* Fig. S3AA-AD). CXCL14 is thus not required for HSC maintenance, HSC function, or bone marrow hematopoiesis, including the generation of megakaryocytes.

Four month-old *Lepr-cre; Cxcl14^fl/fl^* and littermate control mice did not significantly differ in terms of white blood cell (WBC) or red blood cell (RBC) counts but *Lepr-cre; Cxcl14^fl/fl^* mice had significantly lower platelet counts (Fig. 2W; 20% reduction). By 7 months of age, this phenotype was more severe as *Lepr-cre; Cxcl14^fl/fl^* mice exhibited a 37% reduction in platelet counts, with normal WBC and RBC counts (Fig. 2X).

Treatment of *Cxcl14*-dsRed mice with anti-CD42b antibody depleted platelets in the blood (Fig. 2Y) and significantly increased the numbers of *Cxcl14-*DsRed^+^ stromal cells and LepR^+^*Cxcl14-*DsRed^+^ stromal cells in the bone marrow (*SI Appendix,* Fig. S4A-C). There was a trend toward increased *Cxcl14* expression by LepR^+^ cells after platelet depletion but the difference was not statistically significant (*SI Appendix,* Fig. S4D-E). *LepR-Cre;Cxcl14^fl/fl^* mice exhibited significantly delayed platelet regeneration, but not white or red blood cell regeneration, as compared to control mice (Fig. 2Y). Platelet depletion thus increased the number of *Cxcl14* expressing stromal cells in the bone marrow.

*LepR-Cre;Cxcl14^fl/fl^* mice also exhibited significantly delayed platelet regeneration, but not white or red blood cell regeneration, as compared to control mice after sublethal irradiation (Fig. 2Z). Consistent with the fact that irradiation depletes hematopoietic cells and LepR^+^ cells (57), sublethal irradiation reduced white blood cell, red blood cell, and platelet counts (Fig. 2Z) as well as the numbers of *Cxcl14-*DsRed^+^ cells and LepR^+^*Cxcl14-*DsRed^+^ cells in the bone marrow (*SI Appendix,* Fig. S4F-G). The level of *Cxcl14*-dsRed expression per LepR^+^ cell also declined after irradiation (*SI Appendix,* Fig. S4H-I) consistent with the observation that irradiation broadly reduces hematopoietic growth factor expression by LepR^+^ cells (58).

### *Cxcl14* promotes the formation of proplatelet-forming megakaryocytes

Megakaryocytes grow in size and mature into proplatelet-forming megakaryocytes that extend processes into sinusoidal blood vessels to shed platelets into circulation (22). Many mature megakaryocytes and proplatelet-forming megakaryocytes are too large and/or too fragile to be observed by flow cytometry (59, 60), so we also examined these cells by immunofluorescence analysis in bone marrow sections (Fig. 3A and 3B). *Lepr-cre; Cxcl14^fl/fl^* and control bone marrow did not significantly differ in terms of the numbers of all CD41^+^ megakaryocytes (Fig. 3A-C) but the megakaryocytes in control bone marrow were significantly larger (Fig. 3D) and included more proplatelet-forming megakaryocytes (Fig. 3E). We did not observe a significant difference in DNA content between CD42d^+^ megakaryocytes from *Lepr-cre; Cxcl14^fl/fl^* and control mice by flow cytometry (*SI Appendix,* Fig. S5A-B); however, it is hard to interpret this as the most mature megakaryocytes may have been excluded from this analysis as a result of their size (59, 60).

**Figure 3:**
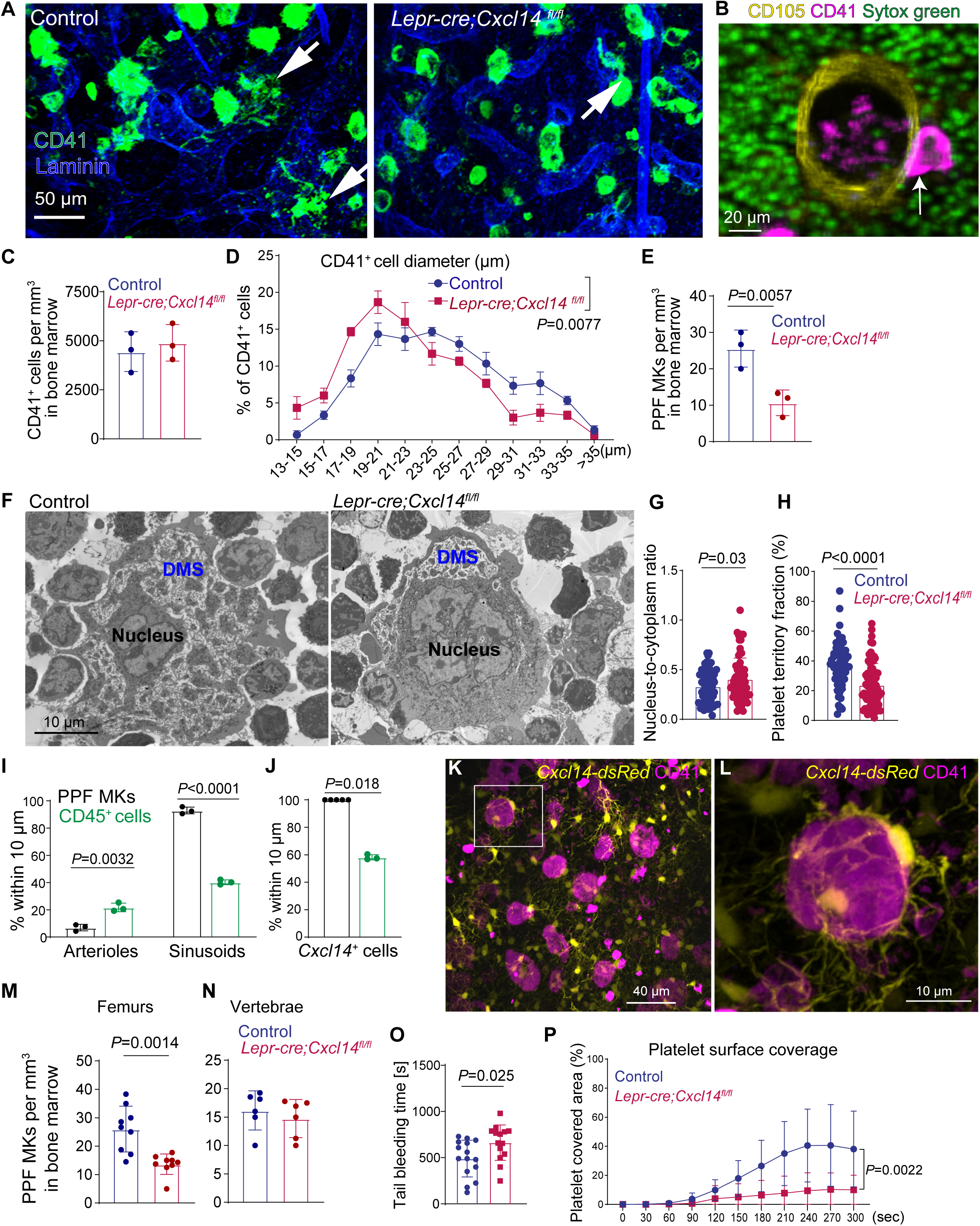
*Lepr-cre; Cxcl14^fl/fl^* mice produced fewer proplatelet-forming megakaryocytes in the bone marrow. (A) Representative images of femur bone marrow from adult *Lepr-cre; Cxcl14^fl/fl^* and control mice showing CD41^+^ proplatelet-forming megakaryocytes (arrows; images are representative of 3 independent experiments). (B) High magnification image of a CD41^+^ proplatelet-forming megakaryocyte adjacent to a sinusoid. (C) The number of CD41^+^ megakaryocytes per mm^3^ in bone marrow sections did not significantly differ between *Lepr-cre; Cxcl14^fl/fl^* and control mice (3 mice per genotype from 3 independent experiments). (D) Distribution of CD41^+^ megakaryocyte cell diameters (3 mice per genotype from 3 independent experiments). (E) The number of CD41^+^ proplatelet-forming megakaryocytes (PPF MKs) per mm^3^ in bone marrow sections (3 mice per genotype from 3 independent experiments). (F) Transmission electron microscopy images of megakaryocytes in the bone marrow of *Lepr-cre; Cxcl14^fl/fl^*and control mice showed reduced demarcation membrane system (DMS) in megakaryocytes from mutant mice. (G–H) Megakaryocyte ultrastructural organization in the bone marrow of adult *Lepr-cre; Cxcl14^fl/fl^* and control mice: Nucleus-to-cytoplasm area ratio (G) and platelet territory fraction (H; the percentage of total megakaryocyte area occupied by platelet territories (3 mice per genotype from 3 independent experiments; each dot represents a megakaryocyte; 57 and 74 megakaryocytes were analyzed for each group). (I) CD41^+^ proplatelet-forming megakaryocytes were significantly more likely than all CD45^+^ cells to localize adjacent to sinusoids and less likely to localize adjacent to arterioles (a total of 3 mice per group from 3 independent experiments). (J) CD41^+^ proplatelet-forming megakaryocytes were significantly more likely than all CD45^+^ cells to localize adjacent to *Cxcl14*-dsRed^+^ stromal cells (3 or 5 mice per group from 3 independent experiments). (K, L) Representative images of femur bone marrow from *Cxcl14^dsRed^* mice showing fine processes from *Cxcl14*^+^ stromal cells wrapping around CD41^+^ megakaryocytes (images are representative of 4 independent experiments). (M, N) The number of CD41^+^ proplatelet-forming megakaryocytes (PPF MKs) per mm^3^ of bone marrow sections from femurs (M) or vertebrae (N) of *Prx1-cre; Cxcl14^fl/fl^* and control mice (6 or 9 mice per genotype from 3 or 4 independent experiments). (O) Bleeding time after severing the tail 1 mm from the end in *Lepr-cre; Cxcl14^fl/fl^* and control mice (a total of 14-15 mice per genotype from 3 independent experiments). (P) Platelet surface coverage over time under flow on collagen-coated surfaces using blood from *Lepr-cre; Cxcl14^fl/fl^* and control mice (a total of 8 mice pre genotype from 3 independent experiments). All data represent mean ± standard deviation and each dot represents a different mouse except in panels D, G, H, and P. All statistical tests were two-sided. Statistical significance was assessed using matched samples two-way ANOVAs followed by Sidak’s multiple comparisons adjustments (C and E), a nested *t*-test (D), Student’s *t*-tests (G, H, and O), or Student’s *t*-tests (I, J, M, and N) or Mann-Whitney tests (J, N and P) followed by Holm-Sidak’s multiple comparisons adjustments, or non-parametric analyses for longitudinal data (nparLD) (P).

During their maturation, megakaryocytes form a demarcation membrane system that envelops developing platelets in the cytoplasm. By transmission electron microscopy, we observed significantly reduced nucleus to cytoplasm ratio and platelet territory area within megakaryocytes from *Lepr-cre; Cxcl14^fl/fl^* as compared to control mice (Fig. 3F-H), indicating impaired megakaryocyte maturation (61, 62).

Arterioles and sinusoids were similar in appearance, density, and diameter in the bone marrow of *Lepr-cre; Cxcl14^fl/fl^* and control mice (*SI Appendix,* Fig. S5H-L). CD41^+^ megakaryocytes and proplatelet-forming megakaryocytes were both significantly more likely than all CD45^+^ hematopoietic cells to localize within 10 µm of sinusoids and significantly less likely than all CD45^+^ cells to localize within 10 µm of arterioles (Fig. 3I and *SI Appendix,* Fig. S5C-D): 72±7.5% of megakaryocytes and 93±7.4% of proplatelet-forming megakaryocytes were within 10 µm of sinusoids. Moreover, 84±7.8% of megakaryocytes and all proplatelet-forming megakaryocytes were within 10 µm of *Cxcl14-*DsRed^+^ cells (Fig. 3J and *SI Appendix,* Fig. S5E). We observed no significant difference in the percentages of megakaryocytes or proplatelet-forming megakaryocytes that localized to sinusoids in *Lepr-cre; Cxcl14^fl/fl^* versus control mice (*SI Appendix,* Fig. S5F and S5G). Lattice light-sheet microscopy revealed that most perisinusoidal megakaryocytes were enveloped by fine processes from *Cxcl14-*DsRed^+^ stromal cells (Fig. 3K-L; see also Movie S1). Larger megakaryocytes tended to have more interactions (within 1 μm) per unit surface area with processes from *Cxcl14*-DsRed^+^ cells than smaller megakaryocytes (*SI Appendix,* Fig. S5M). Megakaryocytes and proplatelet-forming megakaryocytes thus localized mainly to sinusoids but this did not depend upon CXCL14.

To test if CXCL14 functions locally within individual bone marrow compartments or systematically, we assessed hematopoiesis in the long bones (femurs and tibia) and vertebrae of *Prx1-cre; Cxcl14^fl/fl^* mice and littermate controls. Prx1-cre recombines in limb mesenchyme, including in LepR^+^ cells, but not in vertebrae. We observed no significant differences between *Prx1-cre; Cxcl14^fl/fl^* and littermate control mice in terms of red or white blood cell counts but *Prx1-cre; Cxcl14^fl/fl^* mice had significantly lower platelet counts (*SI Appendix,* Fig. S6A). *Prx1-cre; Cxcl14^fl/fl^* and littermate control mice also did not significantly differ in body mass, bone marrow cellularity, or the frequencies of hematopoietic stem or progenitor cells by flow cytometry in long bones or vertebrae (*SI Appendix,* Fig. S6B-AM). However, the number of proplatelet-forming megakaryocytes was significantly lower in femur (Fig. 3M), but not vertebra (Fig. 3N), bone marrow from *Prx1-cre; Cxcl14^fl/fl^* mice. This suggests that *Cxcl14* acts locally within individual bone marrow compartments to promote platelet formation.

### CXCL14 from LepR^+^ cells is necessary to produce normally functioning platelets

*Lepr-cre; Cxcl14^fl/fl^* mice exhibited prolonged bleeding as compared to control mice when their tails were severed 1 mm from the end (Fig. 3O). This could not be explained entirely by the reduction in platelet counts as a more severe platelet depletion would normally be required to prolong bleeding (63). We would not expect CXCL14 produced by LepR^+^ in the bone marrow to alter the function of platelets in the blood because *Lepr-cre; Cxcl14^fl/fl^* mice had normal levels of CXCL14 in the blood (Fig. 2A) and megakaryocytes and platelets do not express *Cxcl14* (*SI Appendix,* Fig. S1D-E). Therefore, we hypothesized that *Cxcl14* deletion from LepR^+^ cells in the bone marrow not only impaired megakaryocyte maturation (Fig. 3F-H) but also undermined the quality of the platelets produced in the bone marrow.

To test this, we compared platelet function between *Lepr-cre; Cxcl14^fl/fl^* and control mice. Platelets from the blood of *Lepr-cre; Cxcl14^fl/fl^* mice showed markedly reduced platelet surface coverage and thrombus growth under flow on collagen as compared to platelets from control mice (Fig. 3P and *SI Appendix,* Fig. S7A). Platelets from *Lepr-cre; Cxcl14^fl/fl^*and control mice exhibited similar surface expression of CD41, CD61 and GPVI (*SI Appendix,* Fig. S7B and S7C), and showed no significant differences in integrin αIIbβ3 activation in response to adenosine diphosphate (ADP) or thrombin receptor-activating peptide 4 (TRAP4) (*SI Appendix,* Fig. S7D), or platelet-leukocyte aggregation in the blood (*SI Appendix,* Fig. S7E). This is consistent with a prior study that found germline deletion of *Cxcl14* is not necessary for platelet activation (47). In contrast, platelets from the blood of *Lepr-cre; Cxcl14^fl/fl^* mice exhibited reduced surface P-selectin (CD62P) levels as compared to control platelets after ADP or TRAP4 treatment, suggesting impaired α-granule secretion (*SI Appendix,* Fig. S7F and S7G). Transmission electron microscopy revealed that platelets from the blood of *Lepr-cre; Cxcl14^fl/fl^* mice had fewer α-granules and more vacuoles than platelets from control mice (*SI Appendix,* Fig. S7I–P). Moreover, α-granule-associated proteins, including VWF, PF4, and THBS1, were depleted in platelets from *Lepr-cre; Cxcl14^fl/fl^* as compared to control mice (*SI Appendix,* Fig. S7H). Thus, loss of CXCL14 from LepR^+^ bone marrow stromal cells impaired the formation of granules in platelets, compromising their ability to form thrombi under flow.

### CXCL14 reprograms lipid metabolism in megakaryocytes

To better understand the cause of the defects in megakaryocyte maturation and platelet function in *Lepr-cre; Cxcl14^fl/fl^* as compared to control mice, we performed RNA sequencing on megakaryocytes from the bone marrow of *Lepr-cre; Cxcl14^fl/fl^* and littermate control mice using paramagnetic beads. Four of the ten most significantly enriched Gene Ontology (GO) terms among the differentially expressed genes related to coagulation and megakaryocyte or platelet differentiation (red highlighted gene sets in Fig. 4A). Nearly all of the megakaryocyte and platelet differentiation genes (64) detected by RNA sequencing were significantly downregulated in megakaryocytes from *Lepr-cre; Cxcl14^fl/fl^* as compared to control mice (red highlighted genes in Fig. 4B). These included genes that encode proteins that regulate granule secretion (Syntaxin-11 (Stx11) and P-selectin (Selp)) as well as granule cargo proteins (Platelet factor 4 (Pf4), Tissue inhibitor of metalloproteinase 3 (Timp3), Thrombospondin-1 (Thbs1), Serpine2, Von Willebrand factor (Vwf), Extracellular matrix protein 1 (Ecm1), and Protein S (Pros1); Fig. 4B). The depletion of gene products that encode proteins necessary for granule secretion and thrombus formation/growth in megakaryocytes within the bone marrow of *Lepr-cre; Cxcl14^fl/fl^* as compared to control mice suggests that the defects in platelet function within these mice arose within the bone marrow, concomitant with the defects in megakaryocyte maturation.

**Figure 4:**
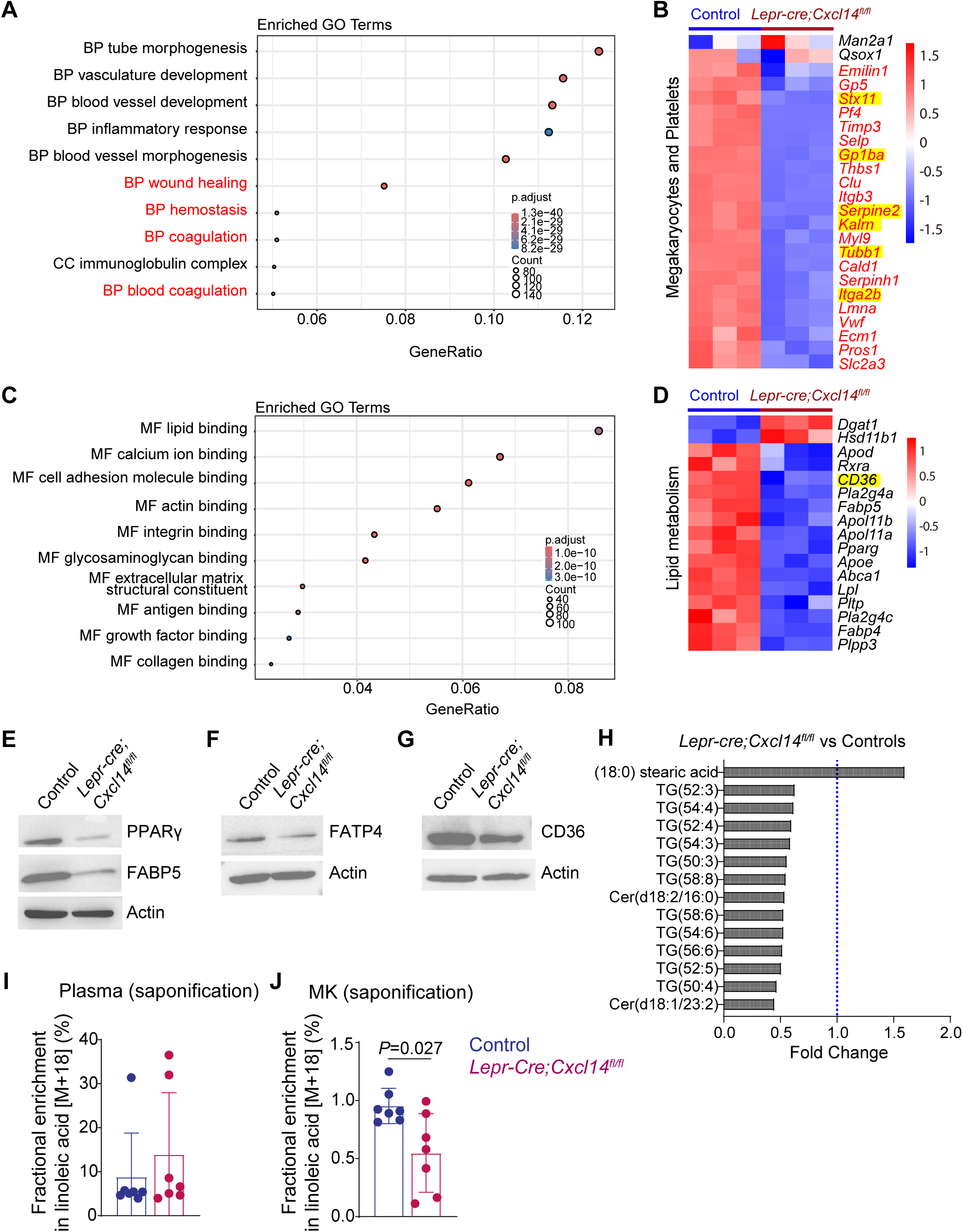
Megakaryocytes from *Lepr-cre; Cxcl14^fl/fl^* mice exhibited changes in lipid metabolism as compared to control megakaryocytes. (A-D) We performed RNA sequencing on megakaryocytes from the bone marrow of *Lepr-cre; Cxcl14^fl/fl^* and control mice (3 mice per genotype from 3 independent experiments). (A) The 10 most significant Gene Ontology (GO) terms that were enriched for genes differentially expressed among *Lepr-cre; Cxcl14^fl/fl^* and control megakaryocytes (log2 fold change > 0.5 in either direction and FDR < 0.05), including GO terms related to biological processes (BP), cellular components (CC) and molecular function (MF). GO terms highlighted in red contain genes involved in megakaryocyte and platelet differentiation or function. (B) Heatmap of genes involved in megakaryocyte or platelet differentiation (z scores of log2-transformed, normalized read counts; list of genes is from ref (64)). (C) The 10 most significantly enriched GO terms related to molecular function (MF) from genes that were differentially expressed among *Lepr-cre; Cxcl14^fl/fl^*and control megakaryocytes (log2 fold change > 0.5 in either direction and FDR < 0.05). (D) Heatmap of genes from the lipid binding GO term that were directly related to lipid metabolism (z scores of log2-transformed, normalized read counts). (E-G) Western blot of proteins that regulate lipid metabolism or fatty acid transport in extracts from CD42d^+^ megakaryocytes isolated using paramagnetic beads from *Lepr-cre; Cxcl14^fl/fl^* or littermate control bone marrow (results are representative of 2 samples per genotype from 2 independent experiments). (H) Lipidomic analysis of CD42d^+^ megakaryocytes from *Lepr-cre; Cxcl14^fl/fl^* and control bone marrow. Significantly changed lipid species (FDR < 0.05, log2 fold change > 0.5 in either direction) in megakaryocytes from *Lepr-cre; Cxcl14^fl/fl^*as compared to control bone marrow (a total of 7-9 mice per genotype from 2 independent experiments). (I and J) Fractional enrichment of ¹³C_18_-linoleic acid in blood plasma or megakaryocytes from *Lepr-cre; Cxcl14^fl/fl^*and control mice after intravenous infusion of ¹³C_18_- linoleic acid for 24 hours (a total of 7 mice per genotype from 2 independent experiments). Data represent mean ± standard deviation and each dot represents a different mouse in I and J. All statistical tests were two-sided. Statistical significance was assessed using the Omics Data Analyzer (see lipidomic methods; H), or a Mann-Whitney test (I) or a Student’s *t*-test (J) followed by Holm-Sidak’s multiple comparisons adjustment.

Some loci that encode genes that were differentially expressed among megakaryocytes from *Lepr-cre; Cxcl14^fl/fl^* as compared to control mice (Figure 4B) contain genetic variants associated with differences in platelet counts or platelet function. Variants at the *GP1BA*, *FAM171A2–ITGA2B*, and *CD36* loci are associated with differences in platelet counts while variants at the *KALRN* and *CTSZ–TUBB1* loci are associated with differences in mean platelet volume (65, 66). Variants at the *KALRN*, *SERPINE2*, and *STX11* loci are associated with differences in platelet reactivity or aggregation (67, 68).

The molecular function GO term most enriched among genes differentially expressed among megakaryocytes from the bone marrow of *Lepr-cre; Cxcl14^fl/fl^* and control mice was related to lipid metabolism (Fig. 4C). Genes in this GO term were expressed at significantly lower levels in megakaryocytes from *Lepr-cre; Cxcl14^fl/fl^*as compared to control mice, including 15 of 17 lipid metabolism genes (Fig. 4D). These included *Pparg*, which encodes the PPARγ transcription factor that regulates fatty acid metabolism (69). PPARγ acts partly by promoting the transcription of fatty acid transporters including *Cd36*, *Fabp5,* and *Fatp4* (70, 71). The levels of PPARγ, FABP5, FATP4 and CD36 protein were all reduced in CD42d^+^ megakaryocytes from the bone marrow of *Lepr-cre; Cxcl14^fl/fl^* mice (Fig. 4E-G). This raised the possibility that megakaryocytes from *Lepr-cre; Cxcl14^fl/fl^* mice have defects in lipid metabolism, including reduced fatty acid uptake. Changes in lipid metabolism, including increased uptake of polyunsaturated fatty acids, are necessary in megakaryocytes for the formation of proplatelet-forming megakaryocytes and platelets, partly to form demarcation membrane system (72–75). People with reduced CD36 function exhibit thrombocytopenia and increased bleeding (73).

To test if PPARγ is necessary for megakaryocyte maturation and platelet formation, we treated wild-type mice with T0070907, a potent and selective inhibitor of PPARγ (76) that can be used to inhibit PPARγ in mice in vivo (77, 78). We confirmed that T0070907 inhibited PPARγ function by treating CD42^+^ megakaryocytes in culture: T0070907 significantly reduced *CD36* and *Fabp4* transcript levels (*SI Appendix,* Fig. S8A-B). Daily intraperitoneal injection of T0070907 for a week did not significantly affect body mass, red or white blood cell counts, bone marrow cellularity, or the frequencies of HSCs, CD41^+^CD150^+^LK megakaryocyte progenitors (MkPs), or immature CD41^+^CD42^-^ megakaryocytes in the bone marrow (*SI Appendix,* Fig. S8C-H). However, it did significantly reduce platelet counts in the blood (*SI Appendix,* Fig. S8D) and the frequencies of more mature CD41^+^CD42^+^ megakaryocytes in the bone marrow by flow cytometry (*SI Appendix,* Fig. S8I) as well as the numbers of CD41^+^ megakaryocytes and proplatelet-forming megakaryocytes in bone marrow sections by immunofluorescence analysis (*SI Appendix,* Fig. S8J-K). T0070907 treatment thus phenocopied the depletion of proplatelet-forming megakaryocytes and platelets in *Lepr-cre; Cxcl14^fl/fl^* mice.

We also treated CD42^+^ megakaryocytes with recombinant CXCL14 and/or T0070907 in culture. CXCL14 significantly increased the formation of proplatelet-forming megakaryocytes and T0070907 blocked this effect (*SI Appendix,* Fig. S8L). CXCL14 thus acts directly on megakaryocytes to promote their maturation into proplatelet-forming megakaryocytes in a manner mediated by PPARγ.

We performed lipidomic analysis on CD42d^+^ megakaryocytes isolated using paramagnetic beads from *Lepr-cre; Cxcl14^fl/fl^*and control bone marrow. We detected 260 lipid species (*SI Appendix,* Table S2). All of the lipids that significantly differed between megakaryocytes from *Lepr-cre; Cxcl14^fl/fl^* and control mice are shown in Fig. 4H. Eleven species of triglycerides and 2 species of ceramides were significantly depleted in megakaryocytes from *Lepr-cre; Cxcl14^fl/fl^* mice. Most of the depleted triglycerides contained polyunsaturated fatty acids, suggesting depletion of polyunsaturated fatty acids in megakaryocytes from *Lepr-cre; Cxcl14^fl/fl^* mice. While cells can synthesize their own saturated and monounsaturated fatty acids, polyunsaturated fatty acids must be obtained from the diet (79).

To test if CXCL14 from LepR^+^ cells promotes polyunsaturated fatty acid uptake by megakaryocytes, we infused *Lepr-cre; Cxcl14^fl/fl^* and littermate control mice for 24 hours with ¹³C_18_-linoleic acid, then isolated CD42d⁺ megakaryocytes from the bone marrow using paramagnetic beads. We observed no significant difference in the fractional enrichment of ¹³C_18_-linoleic acid (¹³C_18_-linolaic acid/all linoleic acid) in blood plasma from *Lepr-cre; Cxcl14^fl/fl^*and control mice (Fig. 4I) but *Lepr-cre; Cxcl14^fl/fl^* megakaryocytes exhibited significantly reduced fractional enrichment as compared to control megakaryocytes (Fig. 4J). This suggests that bone marrow megakaryocytes have less capacity to take up polyunsaturated fatty acids from circulation in the absence of CXCL14 from LepR^+^ cells.

CXCL14 is considered an orphan chemokine, with unknown receptor (52). Nonetheless, evidence has been presented for several potential receptors including CXCR4 (46, 47), Low-density lipoprotein receptor-related protein 1 (LRP1 (50)), Insulin-like growth factor 1 receptor (IGF1R (48)), Atypical chemokine receptor 2 (ACKR2 (51)), and G protein coupled receptor 85 (GPR85 (49)). By RNA sequencing, only *Cxcr4*, *Lrp1*, and *Igf1r* transcripts were detected in bone marrow megakaryocytes (*SI Appendix,* Fig. S8M). By flow cytometric analysis, we detected CXCR4 on 52±6.5% of CD42d^+^ megakaryocytes, while LRP1 and IGF1R were present on 47±4.8% and 12±2.2% of CD42d^+^ megakaryocytes, respectively (*SI Appendix,* Fig. S8N-O). The percentage of CD42d^+^ megakaryocytes that were positive for these candidate receptors did not significantly differ between *Lepr-Cre;Cxcl14^fl/fl^* and control mice (*SI Appendix,* Fig. S8O). Future studies will be required to assess which candidate receptors mediate the effect of CXCL14 on megakaryocyte maturation.

### A high-fat diet rescued platelet formation in *Lepr-cre; Cxcl14^fl/fl^* mice

If the defects in megakaryocyte maturation and platelet formation/function were caused by reduced fatty acid uptake in megakaryocytes, we hypothesized these defects would be rescued by feeding a high fat diet, which would increase the abundance of circulating fatty acids. To test this, we fed *Lepr-cre; Cxcl14^fl/fl^* and littermate control mice normal chow (10% of calories from fat) or either of two high fat diets (both with 60% of calories from fat; Fig. 5A). Both high fat diets significantly increased body mass in *Lepr-cre; Cxcl14^fl/fl^* and control mice as compared to mice fed normal chow (Fig. 5B). Neither diet nor mouse genotype significantly altered white (Fig. 5C) or red blood cell counts (Fig. 5D), or bone marrow cellularity (Fig. 5F). Platelets were depleted in the blood of *Lepr-cre; Cxcl14^fl/fl^* as compared to control mice fed normal chow but not between *Lepr-cre; Cxcl14^fl/fl^* and control mice fed either high fat diet (Fig. 5E). High fat diets thus rescued platelet production in *Lepr-cre; Cxcl14^fl/fl^* mice.

**Figure 5:**
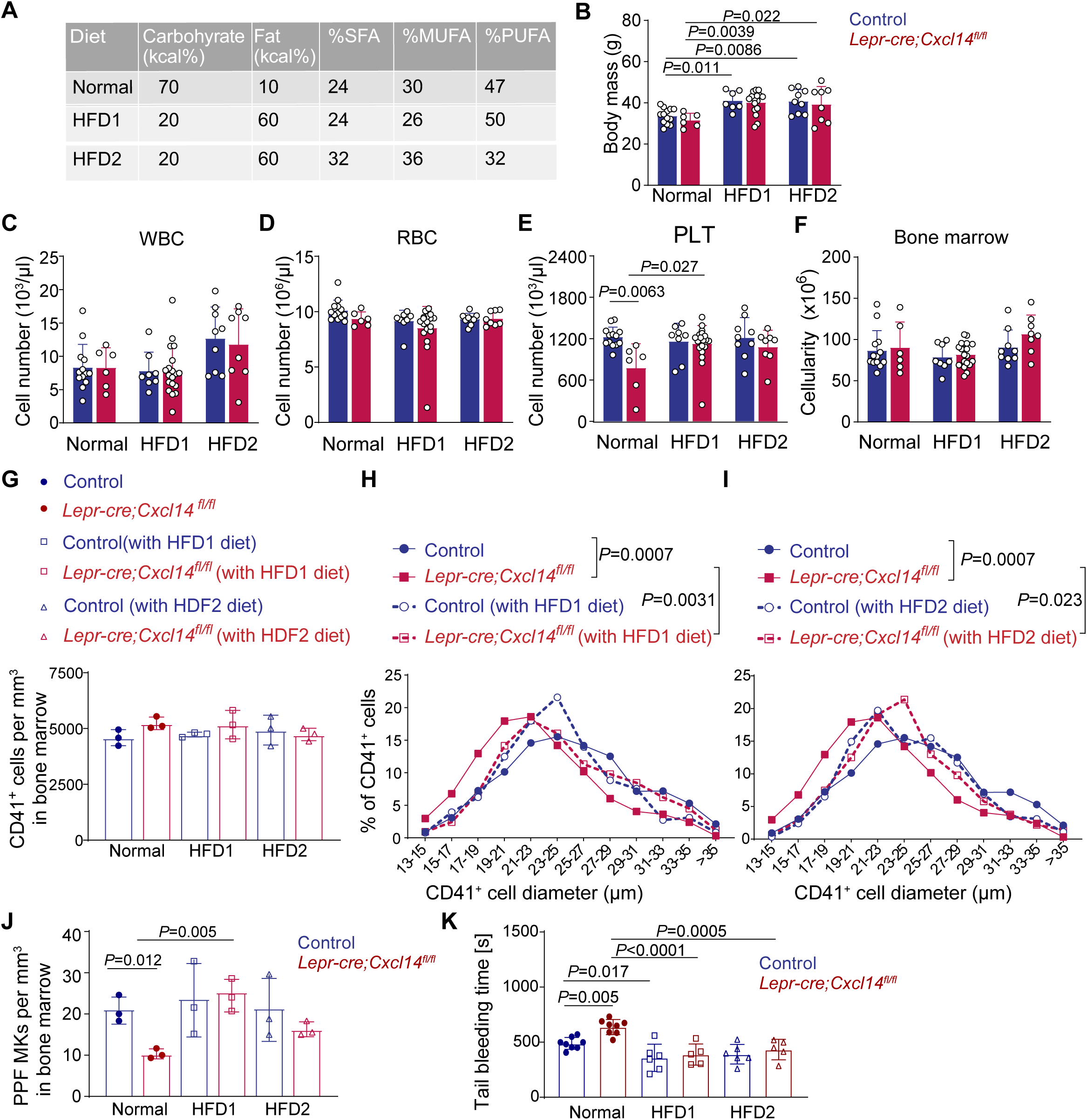
High-fat diet feeding rescued thrombopoiesis in *Lepr-cre; Cxcl14^fl/fl^* mice. (A) *Lepr-cre; Cxcl14^fl/fl^* and littermate control mice were fed normal chow (10% of calories from fat) or either of two high fat diets (HFD1 and HFD2, both of which had 60% of calories from fat) for four weeks. (B) Body mass of *Lepr-cre; Cxcl14^fl/fl^*and control mice after 4 weeks on these diets (a total of 6-16 mice per genotype per diet from 2-3 independent experiments). (C-F) Blood cell counts and bone marrow cellularity (F) in *Lepr-cre; Cxcl14^fl/fl^* and control mice fed each diet (a total of 6-19 mice per genotype per diet from 2-3 independent experiments). (G-I) The number of CD41^+^ megakaryocytes per mm^3^ in sections through the bone marrow (G) and the cell diameter distribution of these cells in mice fed normal chow versus HFD1 (H) or normal chow versus HFD2 (I). (J) The number of CD41^+^ proplatelet-forming megakaryocytes per mm^3^ in sections through the bone marrow of mice on control, HFD1, or HFD2 diets (3 mice per genotype from 3 independent experiments in G-J). (K) Bleeding time after severing the tail 1 mm from the end in *Lepr-cre; Cxcl14^fl/fl^* and control mice fed normal chow, HFD1, or HFD2 (a total of 5-8 mice per genotype per diet from 2-3 independent experiments). All data represent mean ± standard deviation and each dot represents a different mouse except in panels H and I. All statistical tests were two-sided. Statistical significance was assessed using two-way ANOVAs followed by Sidak’s or Dunnett’s multiple comparisons adjustments (B, C, F, G, J and K), Kruskal-Wallis tests followed by Dunn’s multiple comparisons adjustments (D and E), or nested one-way ANOVAs followed by Sidak’s multiple comparisons adjustments (H and I).

*Lepr-cre; Cxcl14^fl/fl^* and control mice did not exhibit significant differences in the frequencies of any hematopoietic stem or progenitor cell population in the bone marrow by flow cytometry, irrespective of diet (*SI Appendix,* Fig. S9A-M). Neither diet nor mouse genotype significantly affected the number of CD41^+^ megakaryocytes in bone marrow sections (Fig. 5G). The size of these megakaryocytes was significantly reduced in *Lepr-cre; Cxcl14^fl/fl^* as compared to control mice fed normal chow but no statistically significant difference was observed in mice fed either high fat diet (Fig. 5H-I). The number of proplatelet-forming megakaryocytes was significantly reduced in *Lepr-cre; Cxcl14^fl/fl^* as compared to control mice fed normal chow but not in mice fed a high fat diet (Fig. 5J). A high fat diet did not significantly alter *Cxcl14* expression in the bone marrow or CXCL14 levels in bone marrow or blood (*SI Appendix,* Fig.S 9Q-T).

The triglycerides that were depleted in *Lepr-cre; Cxcl14^fl/fl^* megakaryocytes from mice fed normal chow (*SI Appendix,* Fig. S9N) were not depleted in *Lepr-cre; Cxcl14^fl/fl^* megakaryocytes from mice fed a high fat diet (*SI Appendix,* Fig. S9O-P; Table S3-5 for the lipids detected). A high-fat diet also rescued the prolonged tail bleeding in *Lepr-cre; Cxcl14^fl/fl^*mice fed normal chow (Fig. 5K) and the levels of granule proteins in platelets from these mice (*SI Appendix,* Fig. S7H). A high fat diet thus eliminated the depletion of triglycerides containing polyunsaturated fatty acids in megakaryocytes in the bone marrow in addition to rescuing the formation of proplatelet-forming megakaryocytes, platelets, and platelet function. This suggests that the defects in platelet function within *Lepr-cre; Cxcl14^fl/fl^* mice arise during platelet formation in the bone marrow as a consequence of reduced fatty acid uptake in bone marrow megakaryocytes.

### Recombinant CXCL14 is sufficient to promote thrombopoiesis

We cultured CD42d^+^ bone marrow cells with or without recombinant CXCL14 (100 ng/ml). CXCL14 significantly increased the numbers of proplatelet-forming megakaryocytes that formed in culture after 16 to 40 hours (Fig. 6A-B). Since only megakaryocytes were present in these cultures, this observation suggests that CXCL14 acts directly on megakaryocytes to promote their maturation into proplatelet-forming megakaryocytes.

**Figure 6:**
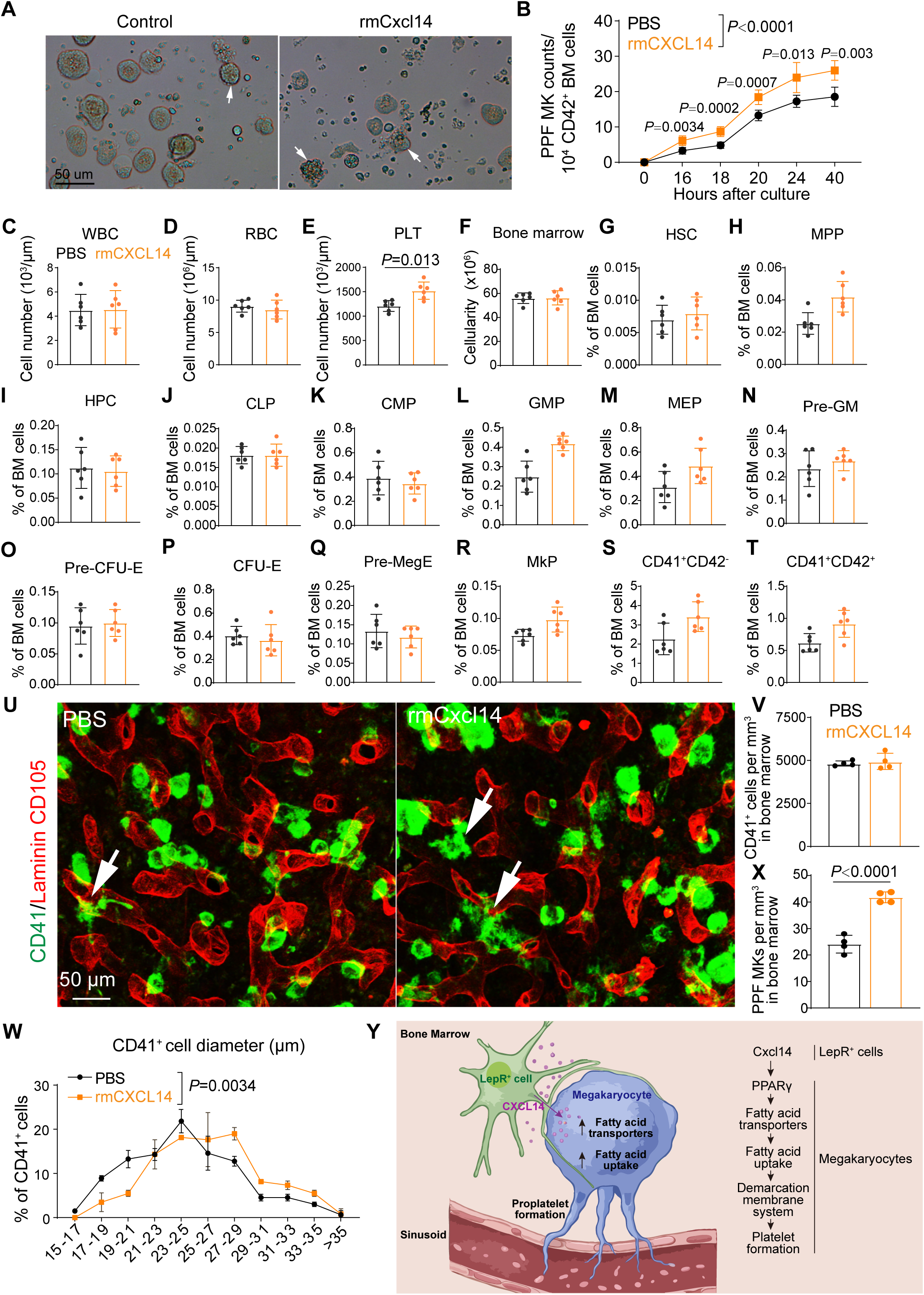
Recombinant CXCL14 is sufficient to increase the production of proplatelet-forming megakaryocytes and platelets. (A, B) Proplatelet-forming megakaryocytes (arrows) formed by CD42⁺ bone marrow cells cultured with or without 100 ng/mL recombinant mouse CXCL14 (rmCXCL14) (images are representative of 5 independent experiments). (C-X) Wild-type mice were intravenously injected with recombinant mouse CXCL14 and then analyzed 16 hours later. Panels C-T show 6 mice per treatment from 3 independent experiments. (C-E) Blood cell counts. (F-T) Bone marrow cellularity (F) and the frequencies of HSCs (G), MPPs (H), HPCs (I), CLPs (J), CMPs (K), GMPs (L), MEPs (M), Pre-GMs (N), Pre-CFU-Es (O), CFU-Es (P), Pre-MegEs (Q), megakaryocyte progenitors (R), CD41^+^CD42d^-^ megakaryocytes (S) and CD41^+^CD42d^+^ megakaryocytes (T) in the bone marrow. (U) CD41^+^ megakaryocytes in femur diaphysis bone marrow (arrows point to proplatelet-forming megakaryocytes; images are representative of 4 mice per treatment from 3 independent experiments). (V and W) Number of CD41^+^ megakaryocytes per mm^3^ in diaphysis bone marrow sections (V) as well as cell diameter distribution (W) (4 mice per group from 3 independent experiments). (X) Number of CD41^+^ proplatelet-forming megakaryocytes per mm^3^ in diaphysis bone marrow sections (4 mice per treatment from 3 independent experiments). (Y) Schematic summary. All data represent mean ± standard deviation and each dot represents a different mouse except in panels B and W. All statistical tests were two-sided. Statistical significance was assessed using matched samples two-way ANOVAs followed by Sidak’s multiple comparisons adjustments (B, G-T, V, and X), Student’s *t*-tests followed by Holm-Sidak’s multiple comparisons adjustments (C-E), a Student’s *t*-test (F), or a nested *t*-test (W).

We also intravenously injected wild-type mice with 20 µg/kg recombinant mouse CXCL14, then analyzed the bone marrow 16 hours later. CXCL14-treated and control mice did not significantly differ in terms of white or red blood cell counts but the CXCL14-treated mice had significantly higher platelet counts (Fig. 6C-E). CXCL14-treated and control mice also did not significantly differ in terms of bone marrow cellularity or the frequencies of HSCs, MPPs, restricted progenitors, or megakaryocytes in the bone marrow by flow cytometry (Fig. 6F-T) or in bone marrow sections (Fig. 6U-V). However, CXCL14-treated mice had significantly larger megakaryocytes (Fig. 6W) and more proplatelet-forming megakaryocytes in the bone marrow as compared to control mice (Fig. 6X). Treatment of wild-type mice with CXCL14 was thus sufficient to increase the formation of proplatelet-forming megakaryocytes and platelets.

## DISCUSSION

CXCL14 appeared to promote the formation of proplatelet-forming megakaryocytes and platelets by remodeling lipid metabolism in megakaryocytes. Increased polyunsaturated fatty acid uptake is necessary in megakaryocytes to form the demarcation membrane system (72, 73). The triglyceride depletion in megakaryocytes from *Lepr-cre; Cxcl14^fl/fl^*mice (Fig. 4H) may reflect the consumption of triglycerides in an effort to sustain demarcation membrane synthesis in the absence of CXCL14. Megakaryocytes in the bone marrow of *Lepr-cre; Cxcl14^fl/fl^* mice had fewer demarcation membranes and the platelets they produced had fewer granules and impaired thrombus growth under flow as compared to control cells. High-fat diet feeding rescued lipid abundance in megakaryocytes (*SI Appendix,* Fig. S9O and S9P), the numbers of proplatelet-forming megakaryocytes (Fig. 5J) and platelets (Fig. 5E) in *Lepr-cre; Cxcl14^fl/fl^* mice, as well as bleeding time (Fig. 5K) and granule protein levels (*SI Appendix,* Fig. S7H) in platelets from *Lepr-cre; Cxcl14^fl/fl^* mice. To our knowledge, these data provide the first evidence that LepR^+^ cells regulate metabolism and terminal differentiation in some hematopoietic cells.

Recent studies have debated whether the lung is a major site of platelet production (80, 81). Our study has not explored whether there are CXCL14-expressing cells in the lung that regulate thrombopoiesis.

The fatty acid transporters whose expression declined in megakaryocytes from *Lepr-cre; Cxcl14^fl/fl^* mice as compared to littermate controls (CD36, FATP4, and FABP5) are not specific for polyunsaturated fatty acids (82). Yet polyunsaturated fatty acid-containing lipids were most prominently depleted from these megakaryocytes (Fig. 4H). Cells can synthesize their own saturated and monounsaturated fatty acids, but not polyunsaturated fatty acids, potentially compensating for the reduced uptake of saturated and monounsaturated fatty acids (79, 83). It is also possible that the depletion of polyunsaturated fatty acid-containing triglycerides in megakaryocytes from *Lepr-cre; Cxcl14^fl/fl^* mice reflects broader defects in lipid uptake or lipid metabolism, not just reduced expression of fatty acid transporters.

## MATERIALS AND METHODS

### Mice

All mice were maintained on a C57BL/6 background. *Scf^GFP^* (*Kitl^tm1.1Sjm^*; a GFP knockin allele that reports *Scf* expression) was generated in our laboratory. *Lepr^cre^* (B6.129(Cg)-Lepr^tm2(cre)Rck^; a Cre allele expressed by *Lepr*^+^ cells) (84) was obtained from Jackson Laboratory. *Prx1-cre* (B6.Cg-Tg(Prrx1-cre)1Cjt; a Cre allele expressed by limb mesenchymal cells) (85). *Cxcl14^dsRed^* and *Cxcl14^flox^* mice were generated in this study. Male and female mice were used in most experiments. C57BL/Ka-Thy-1.1/ C57BL/Ka-Thy-1.2 (CD45.2/CD45.1) mice or C57BL/Ka-Thy-1.2 (CD45.1) mice were used as recipients or as a source of competitor bone marrow cells in transplantation experiments. Recipient mice were given Baytril (0.1 mg/ml in the drinking water) for 1 week prior to, and one month after, irradiation.

Mice were housed in the Animal Resource Center at UT Southwestern in a specific-pathogen-free facility under a 12 h:12 h light:dark cycle with a temperature of 18–24 °C and humidity of 35–60%. For experiments with high-fat diets, 4-5 month-old control and *Lepr-cre; Cxcl14^fl/fl^* mice were placed on the following diets: normal chow (Research Diets, D12450Ji), HFD1 diet (Research Diets, D22050405i), HFD2 diet (Research Diets, D12492i). All procedures were approved by the UT Southwestern Institutional Animal Care and Use Committee.

### Measuring fatty acid uptake in vivo using ¹³C_18_-linoleic acid

Jugular catheters were implanted in *Lepr-cre; Cxcl14^fl/fl^* and control mice. After a week of recovery, the mice were fasted for 4 hours, infused intravenously with ¹³C_18_-linoleic acid conjugated to BSA for 24 hours, then blood and bone marrow were collected. CD42d⁺ megakaryocytes were isolated from bone marrow using paramagnetic beads, then lipids were extracted and analyzed by LC/MS/MS (86). We calculated the theoretical mass of isotopically labeled lipid species (87) then used these values to derive a library of isotopically labeled lipid species that included phospholipids, triglycerides, and free fatty acids. Lipidomic data were searched against this library with 5 ppm mass accuracy. Peaks that appeared to be isotopically labeled lipids were confirmed through MS/MS analysis. All data was corrected for naturally abundant isotopes and the fractional enrichments and pool sizes of each lipid were quantified.

### Recombinant mouse CXCL14 treatment

Recombinant mouse CXCL14 (R&D, Catalog #: 730-XC) was intravenously injected into mice at a dose of 20 µg/kg body weight in a volume of 150 μl per mouse. The equivalent volume of sterile PBS was injected into control mice. The mice were analyzed 16 hours after injection.

## Additional methods can be found in Supporting information

### Materials availability

A material transfer agreement is required to obtain *Cxcl14^flox^* or *Cxcl14^dsRed^*mice in accordance with the regulations of the University of Texas Southwestern Medical Center.

### Data and code availability

All data reported in this paper will be shared by the lead contact upon reasonable request. The RNA sequencing data generated in this paper have been deposited at the Sequence Read Archive (SRA) with a BioProject ID PRJNA1225166. Source code related to the analysis of single-cell RNA sequencing, bulk RNA sequencing, and lipidomics data can be found at https://git.biohpc.swmed.edu/CRI/morrison-lab/scRNASeq, https://git.biohpc.swmed.edu/CRI/morrison-lab/RNASeq, and https://git.biohpc.swmed.edu/CRI/ODA, respectively. Any additional information required to reanalyze the data reported in this paper is available from the lead contact upon request.

### Data sharing statement

For original data, contact

## ACKNOWLEDGEMENTS

S.J.M. is a Howard Hughes Medical Institute (HHMI) Investigator, the Mary McDermott Cook Chair in Pediatric Genetics, the Kathryn and Gene Bishop Distinguished Chair in Pediatric Research, the director of the Hamon Laboratory for Stem Cells and Cancer, and a Cancer Prevention and Research Institute of Texas Scholar. This study was supported partly by funding from the National Institutes of Health (DK118745 to S.J.M.) and the Moody Medical Research Institute (to S.J.M.). R.J.D. is an HHMI Investigator and is supported by NIH grant R35CA220449. D.L. is a scholar of Blood Cancer United. TPM and the CRI Metabolomics Facility are supported by funding from the Cancer Prevention Research Institute of Texas (CPRIT Core Facilities Support Award RP240494). K.M.D. is supported by funding from the Cancer Prevention and Research Institute of Texas (CPRIT; RP250571) and the National Institutes of Health (NIH; U54 CA268072). We thank the BioHPC high-performance computing cloud at UT Southwestern Medical Center for providing computational resources and the Moody Foundation Flow Cytometry Facility. We thank the electron microscopy core facility, supported by NIH grant 1S10OD021685. This article is subject to HHMI’s Open Access to Publications policy. HHMI lab heads have previously granted a nonexclusive CC BY 4.0 license to the public and a sublicensable license to HHMI in their research articles. Pursuant to those licenses, the author-accepted manuscript of this article can be made freely available under a CC BY 4.0 license immediately upon publication.

## Author Contributions

Y.X. and S.J.M. conceived the project and designed and interpreted experiments. Y.X. performed most of the experiments, with technical assistance from A.R.R. and R.Z. and advice related to lipidomic analysis from S.M. and G.G.. M.L. performed the western blots and T.S.T. implanted jugular catheters in mice for isotope-labelled linoleic acid infusion. T.P.M., Y.L., W.G., and R.J.D. helped to perform and analyze the isotope tracing experiments. T.N., Z.S., E.C.J., and K.M.D. provided the light-sheet microscopy platform, performed the imaging, and analyzed the light-sheet imaging data. N.S.T. and D.W.S. helped to perform microfluidic assays for platelet thrombus formation. N.J., T.W., and L.D. helped to prepare samples and perform flow cytometry in experiments that involved many mice. Y.Z. and D.L. observed the fine processes from LepR^+^ cells that envelop megakaryocytes and suggested we test if this was observed in *CXCL14*-expressing cells. Z.Z. performed statistical analyses. Y.X. and S.J.M. wrote the manuscript.

