## Supplemental Data for "Leptin Receptor^+^ cells create a perisinusoidal niche for thrombopoiesis in the bone marrow by synthesizing CXCL14"

**This PDF file includes:**

Supplementary Materials and Methods

Figures S1-S9

Tables S1 to S6

Legend for Movie S1

SI References

**Other supporting material for this manuscript includes the following:**

Movie S1

### Supplementary Materials and Methods

#### Flow cytometric analysis and isolation of stromal cells and megakaryocytes

Bone marrow hematopoietic cells were isolated by flushing the long bones using staining medium ($Ca^{2+}$- and $Mg^{2+}$-free HBSS (HBSS-free) with 2% bovine serum). Spleen cells were obtained by crushing the spleen between two glass slides. The cells were dissociated into a single cell suspension by gently triturating with a 25-gauge needle and then filtering through 40 μm nylon mesh. For analysis of HSCs, cells were stained with fluorophore-conjugated antibodies against lineage markers (CD2 (RM2-5), CD3 (17A2), CD5 (53-7.3), CD8a (53-6.7), Gr1 (RB6-8C5), Ter119 (TER-119) and B220 (RA3-6B2)), c-kit (2B8), Sca1 (D7), CD150 (TC15-12F12.2), and CD48 (HM48-1). For analysis of restricted hematopoietic progenitors, cells were stained with fluorophore-conjugated antibodies against lineage markers, c-kit (2B8), Sca1 (D7), CD16/32 (93), CD34 (RAM34), CD127 (A7R34) and CD135 (A2F10). For analysis of differentiated cells, cells were stained with fluorophore-conjugated antibodies against Mac-1 (M1/70), Gr1 (RB6-8C5), B220 (RA3-6B2), CD3 (17A2), Ter119 (TER-119) and CD71

(R17217). Cells were analyzed using a FACS Canto (BD Biosciences), a FACSAria II (BD Biosciences) or a FACSAria Fusion (BD Biosciences) flow cytometer. Dead cells were identified and gated out of all analyses by including 1 μg/mL 4′,6-diamidino-2-phenylindole (DAPI) or propidium iodide in the staining medium used to resuspend cells for flow cytometry. Flow cytometry data were analyzed using Flowjo (BD Biosciences). The markers used to identify each hematopoietic cell population in this study are shown in Supplemental Table 1 and the flow cytometry gates are shown in Supplemental Figure1E-H.

For flow cytometric analysis of stromal cells, whole femurs and tibias were minced with scissors and enzymatically dissociated in HBSS (with $Ca^{++}$ and $Mg^{++}$) with DNaseI (200U/ml) and LiberaseTL (250mg/ml) by agitating for 30 minutes at 37°C. Bone marrow fragments were allowed to sediment for 1 minute, then the cell suspension was collected and the enzymes were quenched by adding staining medium with 7 mM EDTA. Cells were washed once in staining medium and incubated with biotin-conjugated goat anti-LepR antibody for 90 minutes at 4°C. Cells were washed and then incubated with fluorophore-conjugated antibodies against CD45, Ter119, CD31 and BV421-conjugated streptavidin. For endothelial cell analyses, the cells were stained with anti-CD31 antibody. Dead cells and debris were excluded by gating on forward scatter, side scatter, and the viability dye Ghost Red 780.

To isolate megakaryocytes, tibias and femurs were crushed using a mortar and pestle. The bone marrow cells were then gently triturated and filtered, as described above, to obtain a single-cell suspension. Cells were stained with APC-conjugated anti-CD42d antibody for 20 minutes, followed by enrichment of $CD42d^+$ cells using anti-APC paramagnetic microbeads (Miltenyi Biotec), which were incubated for 30 minutes.

**Generation of mutant mice**

To generate the *Cxcl14*$^{flox}$ and *Cxcl14*$^{dsRed}$ mice, Alt-R S.p. Cas9 Nuclease V3 (Integrated DNA Technologies), sgRNA (Synthego, Supplemental Table S6), tracrRNAs and donor oligos (Azenta, Supplemental Table S6) were microinjected into C57BL/6 zygotes. Chimeric mice were genotyped by sequencing the targeted locus, confirming the correct insertion of loxP or dsRed sequences, and then by PCR analysis. Founders were backcrossed with C57BL/Ka mice for at least three generations prior to analysis.

**Lipid extraction and lipidomic data analysis**

Bone marrow megakaryocytes were transferred to glass vials for Bligh-Dyer lipid extraction using methanol:PBS:chloroform (1:1:2) containing 0.1% formic acid. Mass spectrometry was performed as described (1). Positive and negative lipid intensity data from the experiments were combined, normalized using the relative log expression (RLE) method (2) and log-transformed. Lipid species that significantly differed among treatments (log2 fold change > 0.5 in either direction and false discovery rate (FDR) < 0.05) were identified using Omics Data Analyzer (ODA) V2.2 (3) with generalized linear modelling, taking batch effects into account.

**Primary cell culture**

To assess the frequencies of colony-forming cells, 10,000 bone marrow cells or 300,000 spleen cells were plated per well in Methocult GM M3434 medium supplemented with 10 ng/mL of thrombopoietin and 1x penicillin/streptomycin in six well plates (3 wells per sample). Colonies were counted after 12-14 days using an inverted microscope. To assess the frequency of megakaryocyte colony-forming cells (CFU-Mk), 100,000 bone marrow cells or $1.0 \times 10^6$ spleen cells were plated per well in MegaCult-C in collagen gel supplemented with 10 ng/mL recombinant mouse IL-3, 20 ng/mL of recombinant human (rh) IL-6, 50 ng/mL of rhIL-11 and 50 ng/mL of rhTPO. After 6-8 days of culture, colonies were dehydrated, fixed, and stained for acetylcholinesterase (AChE) activity, which marks mouse megakaryocyte lineage cells, then counted based on morphology and size using an inverted microscope. To assess the formation of proplatelet-forming megakaryocytes (PPF MK), $CD42d^+$ bone marrow cells were isolated using paramagnetic beads and 10,000 cells were plated in 12-well plates coated with poly-L-lysine. The cells were cultured for 2 days in DMEM-low supplemented with 2 mM L-glutamine, 100 U/ml penicillin, 50 mg/ml streptomycin, with or without 100 ng/ml rmCXCL14.

**Megakaryocyte DNA ploidy analysis**

Whole bone marrow cells (~1x$10^6$ cells per sample) were stained with anti-CD42d antibody on ice for 20 minutes followed by Hoechst 33342 (10 µg/ml) plus Verapamil (50µM) in HBSS at 37°C for 30 min, protected from light. After incubation, the cells were washed with ice cold HBSS and resuspend in 300 µl of staining medium for flow cytometric analysis.

**Electron microscopy of bone marrow megakaryocytes and platelets from the blood**

For transmission electron microscopy analysis of megakaryocytes and platelets, bone marrow plugs or blood platelet were collected from *Lepr-cre; Cxcl14$^{fl/fl}$* and control mice. For platelets, blood was collected into sodium citrate anticoagulant. Platelet-rich plasma was obtained by centrifuging at 250 x g for 5 min at room temperature. The platelet-rich plasma was carefully transferred without disturbing the red blood cell layer and centrifuged at approximately 650 x g for 5 min to pellet the platelets. The bone marrow plugs and platelet pellets were fixed in 4% paraformaldehyde, 2.5% glutaraldehyde plus 0.2% picric acid in 0.01M phosphate buffer. After three rinses in 0.1 M sodium cacodylate buffer, the plugs were embedded in 3% low melting agarose, then washed three times in 0.1M sodium cacodylate buffer then post-fixed two times in 1% osmium tetroxide with 1.5 % K3[Fe(CN)6] in 0.1 M sodium cacodylate buffer for 1.5 hours each at room temperature. Tissues were rinsed with water and en bloc stained with 0.5% aqueous uranyl acetate in 25% methanol overnight at 4°C. After five rinses with water, specimens were stained with 0.02M lead nitrate in 0.03M L-aspartate for 30 minutes at 60°C. Samples were dehydrated with increasing concentrations of ethanol and infiltrated with Embed-812 resin and polymerized in a 60°C oven overnight. Blocks were sectioned with a diamond knife (Diatome) on a Leica Ultracut UCT (7) ultramicrotome (Leica Microsystems) and collected onto formvar coated copper grids. Images were acquired on a JEOL JEM-1400 Plus transmission electron microscope operated at 80 kV using an AMT-BioSprint 16M CCD camera.

Transmission electron microscopy images were analyzed using ImageJ and QuPath. For megakaryocyte analysis, the cell boundary was manually delineated to determine total megakaryocyte area. Megakaryocyte nuclei, cytoplasm, demarcation membrane system (DMS), and platelet territories were manually delineated, and the area of each compartment within images was quantified. Platelet territory fraction was calculated by dividing platelet territory by total megakaryocyte area. Individual platelet boundaries were manually annotated to determine platelet area. α-granules and vacuolar structures were identified based on their morphology and manually annotated. The number of α-granules and vacuoles per platelet was counted, and their densities were calculated as the number of structures per $\mu m^2$ of platelet area. The area occupied by α-granules or vacuoles was also measured, and the corresponding area fraction was calculated by dividing the summed area of each structure by the total platelet area.

**Quantitative reverse transcription PCR**

Cells were flow cytometrically sorted from enzymatically dissociated bone marrow into Trizol (Invitrogen). RNA was extracted and reverse transcribed into cDNA using iScript Reverse Transcription Supermix for RT-qPCR. The PCR primers are listed in Supplemental Table 6. Transcript levels were normalized to Actin (Actb) and fold change was calculated based on ΔCt.

**Competitive reconstitution assays**

Adult recipient mice were irradiated using an XRAD 320 X-ray irradiator (Precision X-Ray Inc.) with two doses of 540 rad at least 4 hours apart (1080 rads total). C57BL/Ka (CD45.1/CD45.2 heterozygous) mice were used as recipients. 500,000 unfractionated bone marrow cells from donor (CD45.2) and competitor (CD45.1) mice were mixed and injected intravenously through the tail vein. Recipient mice were bled from 4 to 16 weeks after transplantation to examine the levels of donor-derived Mac1$^{+}$ myeloid cells, B220$^{+}$ B cells, and CD3$^{+}$ T cells in the blood. Red blood cells were lysed with ammonium chloride potassium buffer before antibody staining. The antibodies used to analyze donor chimerism in the blood were anti-CD45.1 (A20), anti-CD45.2 (104), anti-Gr1 (8C5), anti-Mac1 (M1/70), anti-B220 (6B2) and anti-CD3 (KT31.1). For sublethal irradiation, mice were irradiated with one dose of 650 rads.

**RNA-Seq library preparation and data analysis**

Bone marrow CD42d$^{+}$ megakaryocytes were isolated into tubes containing 300 ul of RLT buffer (Qiagen RNeasy Micro kit) and RNA was purified according to the manufacturer's instructions. RNA integrity and concentration were measured using a Pico Bioanalyzer. cDNA libraries were generated using the SMARTer Stranded Total RNA-Seq Kit v2 - Pico Input Mammalian (Clontech). Library fragment size was measured using D1000 Screen Tape (Agilent) and libraries were quantified using the Qubit dsDNA high sensitivity assay kit (Life Technologies). Libraries were sequenced using an Illumina NextSeq 2000. The quality of RNA-seq raw reads was checked using FastQC 0.11.8. Raw reads were mapped to the Ensembl GRCm38 mouse reference genome using STAR 2.7.9a. Mapped reads were quantified using HTSeq 0.9.1. Quantified mapped reads were normalized and gene expression levels were measured as fragments per 1000 exonic bases per million mapped reads (FPKMs) using DESeq2 1.30.0 with R 4.5.0. Differential expression was assessed using DESeq2 1.30.0 and heatmaps were generated using pheatmap 1.0.12 with R 4.5.0. Gene set over-representation analyses were performed and visualized using clusterProfiler 4.10.1 (4) with R 4.5.0.

**Deep imaging of bone marrow and image processing**

Sample preparation and immunostaining of bisected (half) femurs were performed as previously described (5). Femurs were longitudinally cut in half, then stained, and deep imaged.

The staining solution contained 10% DMSO, 0.5% IgePal630 (Sigma), and 5% donkey serum (Jackson Immuno) in PBS. Briefly, femurs were fixed in 4% paraformaldehyde in PBS for 6 hours with gentle rocking at 4°C. After fixation, the bones were embedded in OCT. Each femur was trimmed to a half bone using a cryostat and then washed three times in PBS. Non-specific antibody binding was blocked by incubating half bones in whole mount staining medium (PBS with 5% donkey serum, 0.5% IgePal630, and 10% DMSO) overnight at room temperature. Half bones were then stained in whole mount staining medium with primary antibodies against CD105, laminin, GFP, DsRed/Tomato, LepR, and CD41 or CD45 for three days at room temperature. The samples were washed three times for 5 minutes in PBS and then overnight in PBS. Half bones were then stained with secondary antibodies (Alexa Fluor 647-AffiniPure F(ab′)2 fragment donkey anti-chicken IgY, Alexa Fluor 488-AffiniPure F(ab′)2 fragment donkey anti-rabbit IgG, and 555-conjugated donkey anti-goat antibodies) for three days at room temperature. The samples were washed three times for 5 minutes in PBS and then overnight in PBS. Bone clearing was performed in eppendorf tubes, gently rotating at room temperature, first by dehydrating through a methanol series, before clearing with Murray's clear (1:2 Benzyl Alcohol: Benzyl Benzoate; BABB) solution (5). Half bones were mounted in BABB and three-dimensional images were acquired using a Leica SP8 confocal microscope.

During imaging, $CD41^+$ cells with a shortest axis greater than 15 μm were identified as megakaryocytes. The numbers of megakaryocytes and proplatelet-forming megakaryocytes were quantified in regions distributed across femur diaphysis bone marrow. Confocal z-stacks of 20 μm and 100 μm were acquired from thick bone marrow sections to count $CD41^+$ megakaryocytes and proplatelet-forming megakaryocytes per $mm^3$ of bone marrow, respectively. Three to five regions were analyzed per mouse, encompassing a combined projected bone marrow volume of approximately 0.2-1 $mm^3$ per mouse. The diameter of each megakaryocyte was calculated as the average of the longest and shortest axes of each cell in an optical section through the bone marrow. The distance from each CD41⁺ cell or CD45⁺ cell to the nearest arteriole, sinusoid, and *Cxcl14*-dsRed⁺ cell in the selected region of the diaphysis was measured manually in Imaris slice mode.

**Light-sheet imaging of megakaryocytes in the bone marrow**

Bone marrow plugs were flushed from femurs and tibias and stained as described above for deep imaging. All specimens were imaged using a custom-built cleared tissue axially swept light-sheet microscope (CT-ASLM) (6). In aqueous imaging conditions (refractive index ≈ 1.33), the system provided a field of view of 468 × 468 µm² with an isotropic diffraction-limited resolution of ~470 nm. Following deconvolution, the effective lateral and axial resolutions improved to ~380 nm and ~333 nm, respectively. Imaging in higher refractive index environments, such as in benzyl alcohol/benzyl benzoate (BABB; refractive index = 1.56), further improved the spatial resolution of the microscope at the expense of a reduced effective field of view. Fluorescence images were acquired using a Hamamatsu Fusion camera with a 2048 × 2048 pixel region of interest (full camera chip 2304 x 2304, with a voxel size of 167 × 167 × 200 nm (x, y, z)). Excitation was provided by laser lines at 405, 488, 562, and 652 nm. Emitted fluorescence was spectrally filtered using the following bandpass filters: ET450/50m (Chroma) for 405 nm excitation, FF01-515/30-32 (Semrock) for 488 nm excitation, FF01-595/31-32 (Semrock) for 562 nm excitation, and BLP01-647R/31-32 (Semrock) for 652 nm excitation. Microscope control, synchronization of illumination and detection, and automated volumetric acquisition were performed using navigate, an open-source, Python-based microscope control platform (7).

Image processing and analysis was performed using custom Python-based workflows unless otherwise noted. Raw volumetric datasets were deconvolved using PyPetaKit5D using experimentally measured point spread functions, following established procedures (8).

Deconvolution was applied only to the *Cxcl14* $^{+}$ cells prior to segmentation and quantitative analysis. Megakaryocytes were segmented using u-Segment3D (9). To improve elineation of fine membrane features, particularly in regions of low contrast or complex morphology, a guided filter was applied. For downstream analyses, megakaryocytes were defined as segmented objects with an effective diameter greater than 15 µm.

Cell diameter and surface area were estimated from the total number of voxels comprising each segmented object, assuming a spherical geometry. *Cxcl14*$^{+}$ cells were segmented using a custom pipeline optimized for detecting fine, tubular morphologies. This workflow included intensity normalization, denoising, background subtraction, and multi-scale vesselness filtering using a Sato filter implemented in scikit-image. The vesselness response was thresholded to generate an initial binary mask, followed by morphological refinement steps including binary closing, removal of small objects, and hole filling to improve continuity and suppress spurious detections. For each megakaryocyte, a cropped subvolume encompassing the segmented cell and an additional buffer of 64 pixels was extracted. Within each cropped region, the Euclidean distance from the surface of the cell of interest to voxels identified as *Cxcl14*$^{+}$ cell structures was computed, up to a maximum distance of 50 voxels. For quality control, maximum intensity projections of each cropped region, including the segmented target cell and adjacent *Cxcl14*$^{+}$ cell structures were exported to verify segmentation accuracy. We reported *Cxcl14*$^{+}$ voxel counts per $\mu m^2$ of cell surface area and assessed whether megakaryocyte proximity to *Cxcl14*$^{+}$ cell structures correlated with cell size. Pearson correlation coefficients were computed between spherical radius and normalized *Cxcl14*$^{+}$ voxel count.

**Platelet depletion and activation**

Thrombocytopenia was induced by intravenous injection of anti-CD42b antibody (Emfret Analytics, Cat #R300) at a dose of 2 µg/g body mass in a volume of 150 µl per mouse. The control group received an equivalent volume of sterile PBS. Complete blood cell counts were performed at the indicated time points. To induce platelet activation, whole blood was incubated with adenosine diphosphate (Fisher scientific, Cat #22-515-225) or thrombin receptor-activating peptide 4 (Sigma-Aldrich, Cat #A3227-1MG) for 10 minutes at room temperature. Platelets were identified based on forward and side scatter as well as CD41 staining, followed by analysis of JON/A (activated integrin αIIbβ3) and CD62P (P-selectin) antibody staining. Samples not treated with adenosine diphosphate or thrombin receptor-activating peptide 4 served as negative controls.

**Tail vein bleeding time**

One mm of the mouse tail tip was cut off with a scalpel. Tail bleeding was monitored by gently absorbing blood on filter paper without making contact with the wound site. Bleeding was determined to have ceased when no blood was observed on the paper. Experiments were stopped manually after 20 min by cauterization to prevent excessive blood loss.

**Microfluidic thrombus formation assay**

The adhesion of platelets under shear flow to micropatterns of collagen was tested using methods previously described (10-12). Initially, micropatterns of 250 µm wide lines of type I bovine collagen were deposited on to Aquasil (Thermo Scientific, PI42799) coated glass coverslips (#1.5, 50 x 45 mm, Brain Research Laboratories, #4550-1.5D) by attaching a straight channel microfluidic device (width = 250 µm, height = 100 µm, length = 22 mm) to the glass coverslip following treatment with air plasma in a plasma cleaner (PDC-32G, Harrick Plasma) for 1 minute on high RF at 400 mTorr pressure, resulting in a reversible bond. Next a solution of type I bovine collagen (0.05 mg/mL) was perfused through the microchannel and incubated at 37°C for 1 hour. After perfusing buffer through the microchannel to remove any unbound protein, the microfluidic stamp was removed from the glass coverslip, and the collagen modified substrate was dried under air. Next the region of the glass coverslip containing the micropatterned collagen was protected by covering it with a small slab of PDMS before being placed in a plasma cleaner alongside a second straight channel microfluidic device (width = 750 µm, height = 80 µm, length = 22 mm) and exposed to air plasma on high RF for 60 seconds at 400 mTorr. The second straight channel was then irreversibly sealed to the glass coverslip containing the line of micropatterned collagen by placing the straight channel on top of the collagen line, intersecting the micropattern at a right angle. Next a solution of 2% bovine serum albumin (BSA) in PBS was perfused through the microfluidic channel and allowed to adsorb to the uncoated regions of the glass coverslip for 30 minutes to reduce non-specific adhesion. After the BSA blocking step, the microfluidic chamber was placed on the stage of an inverted microscope (Nikon), and the chamber was filled with Hank's balanced salt solution (HBSS). An inlet reservoir of the microfluidic device was filled with mouse blood that had been labeled with 10 µM mepacrine for 10 minutes at 37°C prior to perfusion. The fluorescently labeled blood was then pulled through the channel over the micropatterned collagen line using a syringe pump (PHD ULTRA CP Syringe Pump, Harvard Apparatus) at a flow rate of 5.04 µL/min which corresponded to a shear rate of 100 $s^{-1}$. Fluorescent images of the mepacrine labeled platelets were acquired with a 20x objective (NA = 0.5) and recorded with Olympus DP71 digital camera every 30 seconds for 5 minutes. The amount of platelet adhesion to the micropatterned collagen line was measured and quantified with ImageJ analysis software using methods similar to those described previously. Individual platelets and thrombi were identified by thresholding and image segmentation (10, 13).

**Protein extraction and western blot analysis**

Bone marrow $CD42d^+$ megakaryocytes or circulating platelets were washed twice in HBSS then lysed in 4x Laemmli buffer (Biorad, 161-0747) with 355 mM 2-mercaptoethanol (Sigma, M7522) and vortexed. Samples were then boiled at 100°C for 10 minutes, centrifuged for 10 minutes at 16,000 x g, and the supernatants were loaded on 4-20% Mini-PROTEAN TGX gels (Biorad, 4561094). The gels were transferred onto 0.2 µm PVDF membranes (BioRad, 1704156) by semi-dry transfer using BioRad TransBlot Turbo transfer. Signals were detected using Pierce ECL Western Blotting Substrate chemiluminescence (Thermo Scientific, 32016).

**ELISA analysis**

Plasma was separated from whole blood collected in EDTA-containing tubes by centrifugation at 900 x g for 10 minutes. Bone marrow serum from long bones was collected by centrifugation at 2200 x g for 5 minutes in 10 µl PBS. CXCL14 levels were measured using a mouse CXCL14 ELISA Kit (Invitrogen, Catalog # EM526RB) according to the instructions.

**Statistical methods**

In each type of experiment, multiple mice were tested in multiple independent experiments performed on different days. Mice were allocated to experiments randomly and samples processed in an arbitrary order, but formal randomization techniques were not used. No formal blinding was applied when performing the experiments or analyzing the data. Sample sizes were not pre-determined based on statistical power calculations but were based on our experience with these assays. No data were excluded.

Prior to analyzing the statistical significance of differences among treatments, we tested whether data were normally distributed and whether variance was similar among groups. To test for normality, we performed the Shapiro–Wilk tests when $3 \le n < 20$ or D'Agostino Omnibus tests when $n \ge 20$. To test whether variability significantly differed among groups, we performed *F*-tests (for experiments with two groups) or Levene's median tests (for experiments with more than two groups). When the data significantly deviated from normality or variability significantly differed among groups, we log2-transformed the data and tested again for normality and variability. If the transformed data no longer significantly deviated from normality and equal variability, we performed parametric tests on the transformed data. If log2-transformation was not possible or the transformed data still significantly deviated from normality or equal variability, we performed non-parametric tests on the non-transformed data. For large datasets (n > 50 for each group), we applied parametric tests according to the Central Limit Theorem (CLT) if data were not normally distributed but variance was similar among groups.

When data or log2-transformed data were normal and equally variable, statistical analyses were performed using Student's *t*-tests (when there were two groups), one-way ANOVAs (when there were more than two groups), or two-way ANOVAs/matched samples two-way ANOVAs (when there were two or more groups with multiple diets, cell populations, or time points). When the data or log2-transformed data were normally distributed but unequally variable, statistical analysis was performed using Welch's *t*-tests (when there were two groups) or Welch's one-way ANOVAs (when there were more than two groups). When the data or log2-transformed data were not normally distributed, statistical analysis was performed using Mann-Whitney tests (when there were two groups) or Kruskal-Wallis tests (when there were more than two groups). After ANOVAs, P-values from multiple comparisons were adjusted using Dunnett's (when there were more than two groups and comparisons were between a control group and other groups) or Sidak's method (when there were more than two groups and planned comparisons). Holm-Sidak's method was used to adjust comparisons involving multiple Student's *t*-tests, Welch's *t*-tests, or Mann-Whitney tests. Dunnet's T3 method was applied after Welch's one-way ANOVAs and Dunn's method was applied after Kruskal-Wallis tests for multiple comparisons adjustments. Nested *t*-tests or nested one-way ANOVAs were used when multiple measurements were obtained from the same mice. Nested one-way ANOVAs were followed by Sidak's multiple comparisons adjustments. All statistical tests were two-sided. All data represent mean ± standard deviation where applicable. Statistical tests were performed using GraphPad Prism V10.4.1 or R 4.5.0.

### Supplemental Figures and Legends

#### Fig. S1

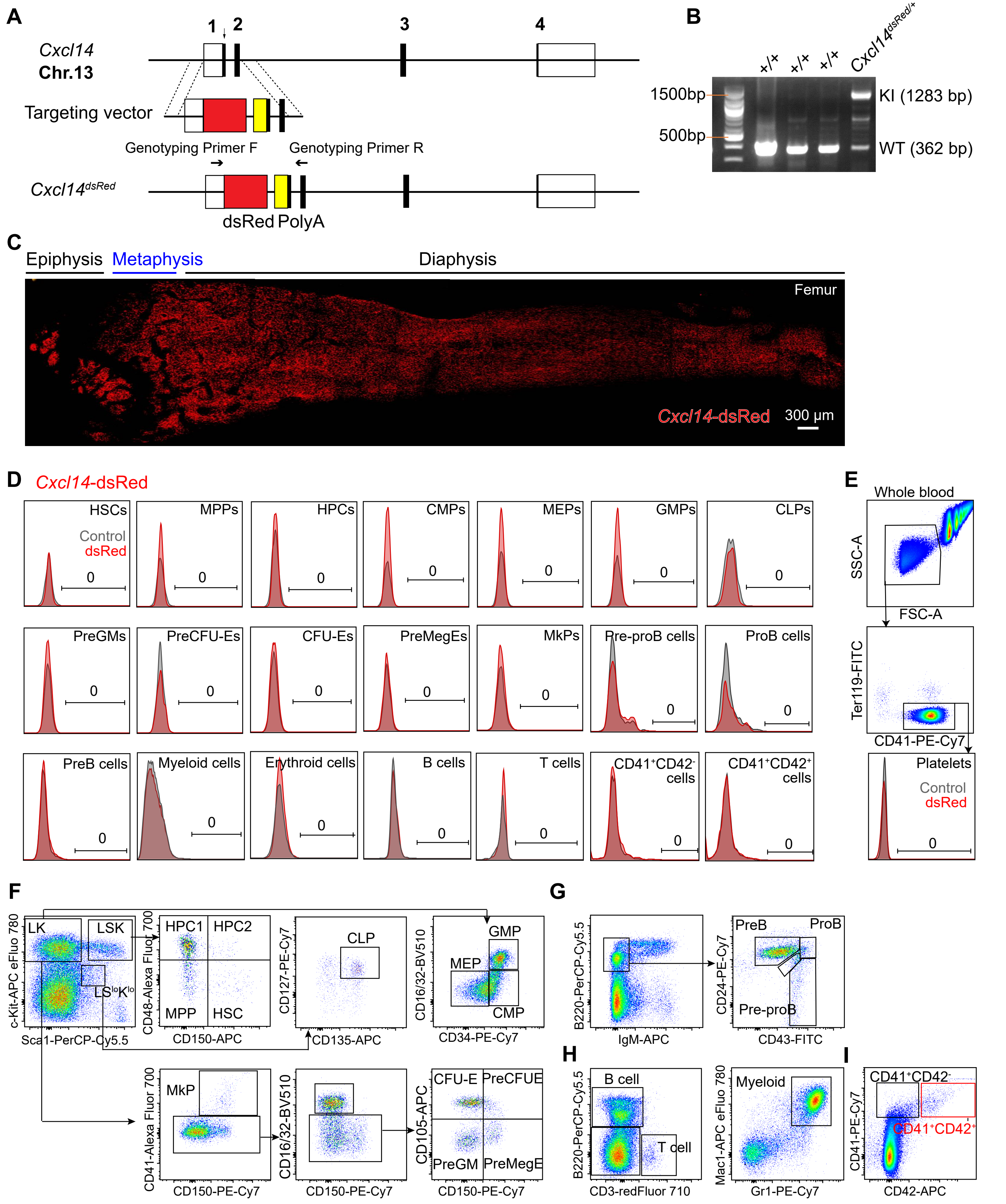

**Fig. S1, Related to Figure 1. Generation of a *Cxcl14*$^{dsRed}$ mouse reporter allele and the flow cytometry gating strategy for the isolation of hematopoietic cell populations.** (A) We inserted a *Cxcl14-dsRed* cassette after the start codon in exon 1 of the *Cxcl14* gene to create a *Cxcl14*$^{dsRed}$ reporter allele. Open boxes indicate untranslated sequences in exons and black boxes indicate translated sequences in exons. Correctly targeted founder mice were identified by long-range PCR with primers flanking the targeted region, followed by Sanger sequencing of the amplicon. (B) PCR genotyping of genomic DNA confirmed germline transmission of the *Cxcl14-dsRed* allele. Mice were backcrossed at least three times onto a C57BL/Ka background before analysis. (C) Representative low magnification image of femur bone marrow from an adult *Cxcl14*$^{dsRed}$ mouse (image is representative of 6 mice). (D-E) We observed no *Cxcl14*$^{dsRed}$ expression by hematopoietic stem and progenitor cells or differentiated hematopoietic cells from the bone marrow or platelets from the blood of *Cxcl14*$^{dsRed}$ mice (data are representative of 3 independent experiments). (F-I) Representative flow cytometry gates used to identify hematopoietic stem and progenitor cell populations analyzed in this study. The markers for each cell population are listed in Table S2.

Fig. S2

A

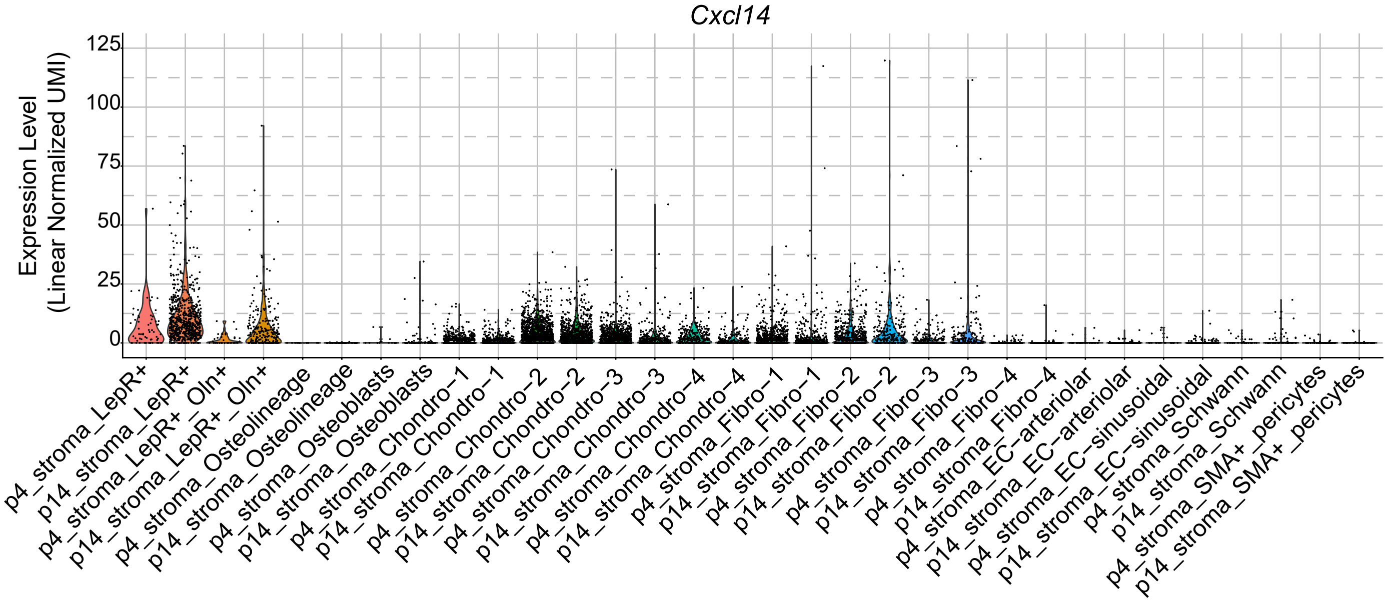
Cxcl14
Expression Level
(Linear Normalized UMI)
125
100
75
50
25
0
p4_stroma_LepR+
p14_stroma_LepR+
p4_stroma_LepR+_Oln+
p14_stroma_LepR+_Oln+
p4_stroma_Osteolineage
p14_stroma_Osteolineage
p4_stroma_Osteoblasts
p14_stroma_Osteoblasts
p4_stroma_Chondro−1
p14_stroma_Chondro−1
p4_stroma_Chondro−2
p14_stroma_Chondro−2
p4_stroma_Chondro−3
p14_stroma_Chondro−3
p4_stroma_Chondro−4
p14_stroma_Chondro−4
p4_stroma_Fibro−1
p14_stroma_Fibro−1
p4_stroma_Fibro−2
p14_stroma_Fibro−2
p4_stroma_Fibro−3
p14_stroma_Fibro−3
p4_stroma_Fibro−4
p14_stroma_Fibro−4
p4_stroma_EC−arteriolar
p14_stroma_EC−arteriolar
p4_stroma_EC−sinusoidal
p14_stroma_EC−sinusoidal
p4_stroma_Schwann
p14_stroma_Schwann
p4_stroma_SMA+_pericytes
p14_stroma_SMA+_pericytes

B

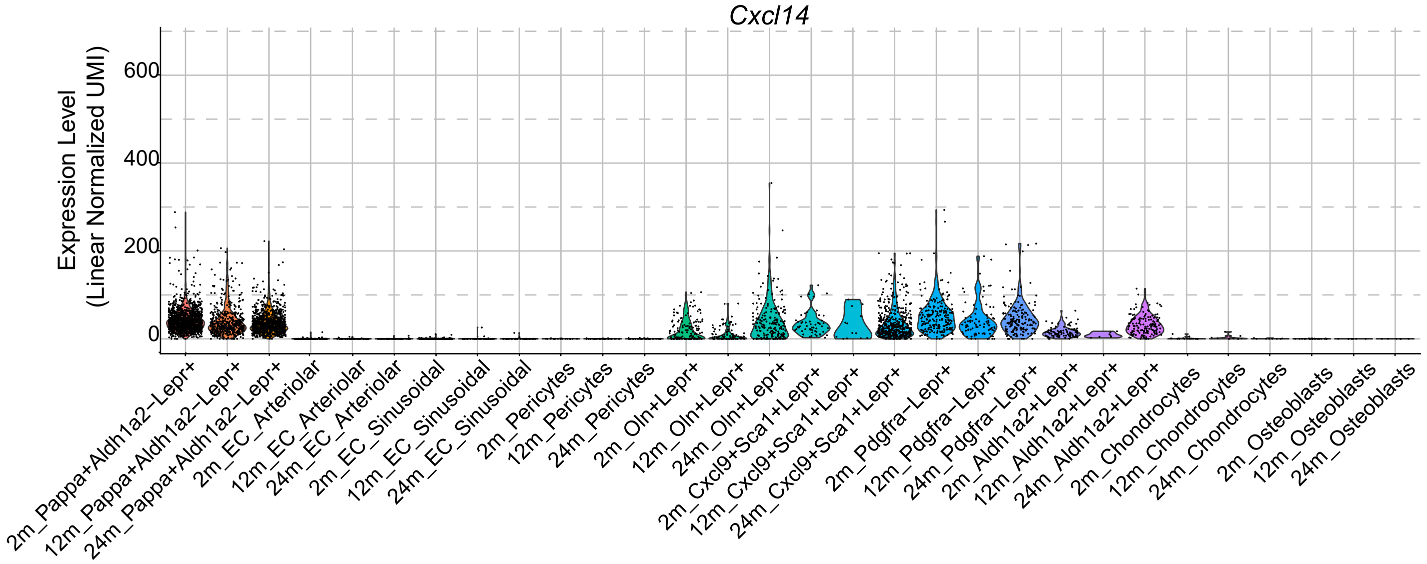
Cxcl14
Expression Level
(Linear Normalized UMI)
600
400
200
0
2m_Pappa+Aldh1a2−Lepr+
12m_Pappa+Aldh1a2−Lepr+
24m_Pappa+Aldh1a2−Lepr+
2m_EC_Arteriolar
12m_EC_Arteriolar
24m_EC_Arteriolar
2m_EC_Sinusoidal
12m_EC_Sinusoidal
24m_EC_Sinusoidal
2m_Pericytes
12m_Pericytes
24m_Pericytes
2m_Oln+Lepr+
12m_Oln+Lepr+
24m_Oln+Lepr+
2m_Cxcl9+Sca1+Lepr+
12m_Cxcl9+Sca1+Lepr+
24m_Cxcl9+Sca1+Lepr+
2m_Pdgfra−Lepr+
12m_Pdgfra−Lepr+
24m_Pdgfra−Lepr+
2m_Aldh1a2+Lepr+
12m_Aldh1a2+Lepr+
24m_Aldh1a2+Lepr+
2m_Chondrocytes
12m_Chondrocytes
24m_Chondrocytes
2m_Osteoblasts
12m_Osteoblasts
24m_Osteoblasts

**Fig. S2, Related to Figure 1. Single cell RNA sequencing analysis of *Cxcl14* expression by bone marrow stromal cells in early postnatal (day 4 and 14) and adult (2, 12, 24 months of age) bone marrow.** These data were obtained from published studies ref (14) in A and ref (15) in B.

### Fig. S3

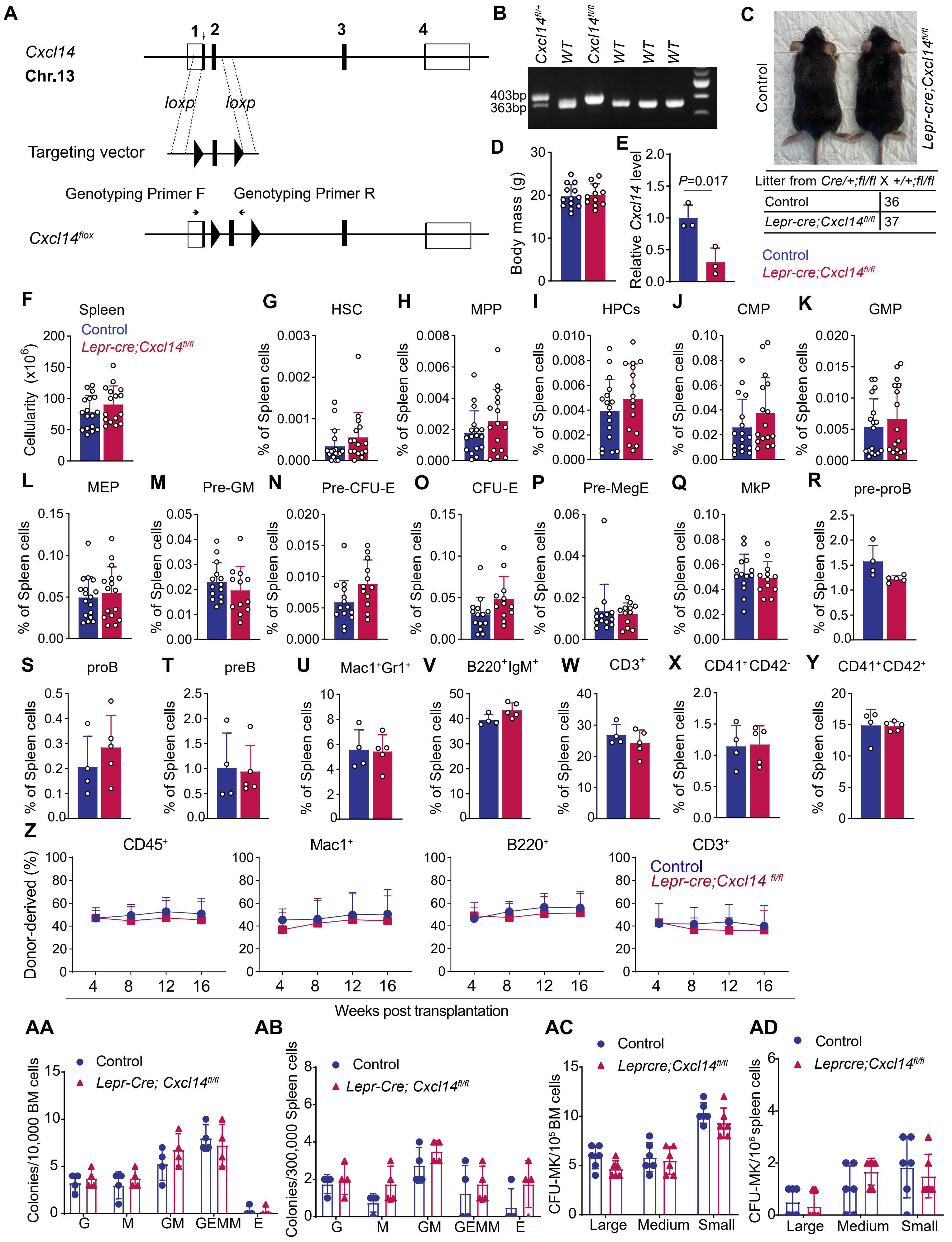

A
Cxcl14
Chr.13
1 2 3 4
loxp
loxp
Targeting vector
Genotyping Primer F
Genotyping Primer R
Cxcl14flox
B
Cxcl14fl/+ WT Cxcl14fl/fl WT WT WT
403bp
363bp
C
Control
Lepr-cre;Cxcl14fl/fl
Litter from Cre/+;fl/fl X +/+;fl/fl
Control 36
Lepr-cre;Cxcl14fl/fl 37
Control
Lepr-cre;Cxcl14fl/fl
D
Body mass (g)
E
Relative Cxcl14 level
P=0.017
F
Spleen
Control
Lepr-cre;Cxcl14fl/fl
Cellularity (x10^6)
G HSC
H MPP
I HPCs
J CMP
K GMP
L MEP
M Pre-GM
N Pre-CFU-E
O CFU-E
P Pre-MegE
Q MkP
R pre-proB
S proB
T preB
U Mac1+Gr1+
V B220+IgM+
W CD3+
X CD41+CD42-
Y CD41+CD42+
% of Spleen cells
Z
CD45+
Mac1+
B220+
CD3+
Control
Lepr-cre;Cxcl14 fl/fl
Donor-derived (%)
Weeks post transplantation
AA
Control
Lepr-Cre; Cxcl14fl/fl
Colonies/10,000 BM cells
G M GM GEMM E
AB
Control
Lepr-Cre; Cxcl14fl/fl
Colonies/300,000 Spleen cells
G M GM GEMM E
AC
Control
Leprcre;Cxcl14fl/fl
CFU-MK/10^5 BM cells
Large Medium Small
AD
Control
Leprcre;Cxcl14fl/fl
CFU-MK/10^6 spleen cells
Large Medium Small

**Fig. S3, Related to Figure 2. Generation and characterization of *Lepr-cre; Cxcl14*$^{fl/fl}$ mice.** (A) Targeting strategy to generate the *Cxcl14*$^{flox}$ allele. The donor ssDNA contained loxp sequences on both sides of exon 2. The loxp insertion sites were chosen to avoid disrupting sequences conserved among species. Cre mediated recombination of this allele would be expected to cause a strong loss of CXCL14 function as it would cause a frame-shift. The *Cxcl14* floxed allele was generated by injecting C57BL/6 zygotes with donor ssDNA, Cas9 protein and 2 sgRNAs targeting the sequences where loxp sites were inserted. Open boxes represent untranslated sequences and black boxes indicate translated sequences in exons. (B) PCR genotyping of genomic DNA confirmed germline transmission of the *Cxcl14*$^{flox}$ allele using the genotyping primers showed in panel (A). (C and D) *Lepr-cre; Cxcl14* $^{fl/fl}$ mice were born in normal numbers and were grossly normal in size and appearance (C) as well as body mass (D) at fourteen weeks of age (11-13 mice per genotype from 5 independent experiments). (E) *Cxcl14* transcript levels in LepR$^{+}$ cells from the bone marrow of *Lepr-cre; Cxcl14* $^{fl/fl}$ mice and littermate controls (3 samples per genotype, each from 1-2 mice, from 3 independent experiments). (F-Y) Spleen cellularity and the frequencies of hematopoietic stem and progenitor cells in the spleens of *Lepr-cre; Cxcl14* $^{fl/fl}$ and littermate control mice (a total of 4-17 mice per genotype from 3-7 independent experiments). (Z) Donor-derived CD45$^{+}$ (overall), myeloid, B, and T cells in the blood of recipient mice competitively transplanted with 500,000 *Lepr-cre; Cxcl14* $^{fl/fl}$ donor bone marrow cells along with 500,000 competing wild-type bone marrow cells (4 donors per genotype were transplanted into a total of 16-17 irradiated recipients per genotype in 4 independent experiments). (AA and AB) CFU-G, CFU-M, CFU-GM, CFU-GEMM and BFU-E colonies formed by 10,000 whole bone marrow cells (AA) or 300,000 spleen cells (AB) in Methocult GM M3434 (4 mice per genotype from 2 independent experiments). (AC and AD) CFU-Mk colonies formed by 100,000 bone marrow cells (AC) or 1,000,000 spleen cells (AD) in Methocult-C medium in collagen gel (6 mice per genotype from 3 independent experiments). All data represent mean ± standard deviation. Each dot represents a different mouse except in panel Z. All statistical tests were two-sided. Statistical significance was assessed using Student’s *t*-tests (D-F), Student’s *t*-tests (AA and AC) or Mann-Whitney tests (G-Y, AB, and AD) followed by Holm-Sidak’s multiple comparisons adjustments, or matched samples two-way ANOVAs followed by Sidak's multiple comparisons adjustments (Z).

### Fig. S4

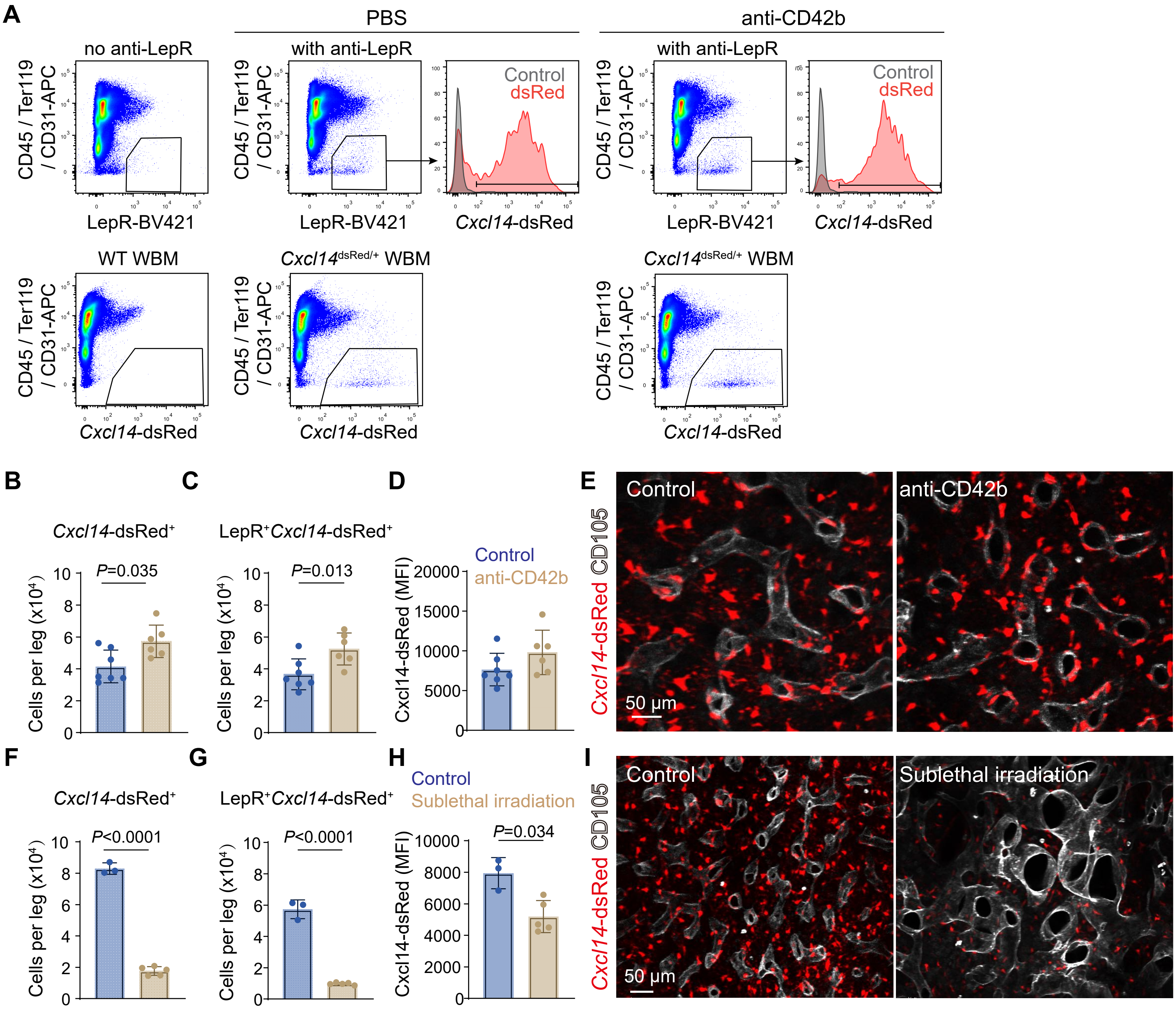

A
PBS
anti-CD42b
no anti-LepR
with anti-LepR
with anti-LepR
CD45 / Ter119 / CD31-APC
LepR-BV421
Control
dsRed
Cxcl14-dsRed
WT WBM
Cxcl14dsRed/+ WBM
Cxcl14dsRed/+ WBM
B
C
D
E
Cxcl14-dsRed+
LepR+Cxcl14-dsRed+
Cells per leg (x10^4)
P=0.035
P=0.013
Control
anti-CD42b
Cxcl14-dsRed (MFI)
Control
anti-CD42b
50 μm
Cxcl14-dsRed CD105
F
G
H
I
P<0.0001
P<0.0001
P=0.034
Control
Sublethal irradiation
Control
Sublethal irradiation
50 μm

**Fig. S4, Related to Figure 3. Dynamics of bone marrow *Cxcl14* expression after platelet depletion and sublethal irradiation.** (A) Flow cytometric analysis of enzymatically dissociated bone marrow from *Cxcl14*-dsRed mice 3 days after treatment with PBS control or platelet depletion with anti-CD42b antibody. (B-D) Number of *Cxcl14*-dsRed$^+$ stromal cells (B) or LepR$^+$*Cxcl14*-dsRed$^+$ cells (C) in the bone marrow per leg (femur and tibia) and mean *Cxcl14*-dsRed fluorescence intensity per LepR$^+$ cell (D) in *Cxcl14*-dsRed mice treated with anti-CD42b antibody or PBS control (6-7 mice per treatment in 3 independent experiments). (E) Femur bone marrow from *Cxcl14*-dsRed mice 3 days after treatment with anti-CD42b antibody or PBS control (images are representative of 3 independent experiments). (F-I) Number of *Cxcl14*-dsRed$^+$ stromal cells (F) or LepR$^+$*Cxcl14*-dsRed$^+$ cells (G) in the bone marrow per leg (femur and tibia) and mean *Cxcl14*-dsRed fluorescence intensity per LepR$^+$ cell (H) in *Cxcl14*-dsRed mice with or without sublethal irradiation (a total of 3-5 mice per treatment in 2 independent experiments). (I) Femur bone marrow from unirradiated (left) or sublethally irradiated (right; 10 days after irradiation) *Cxcl14*-dsRed mice (images are representative of 2 independent experiments). All data represent mean ± standard deviation. Each dot represents a different mouse. All statistical tests were two-sided. Statistical significance was assessed using two-way ANOVAs followed by Sidak's multiple comparisons adjustments (B-D and F-H).

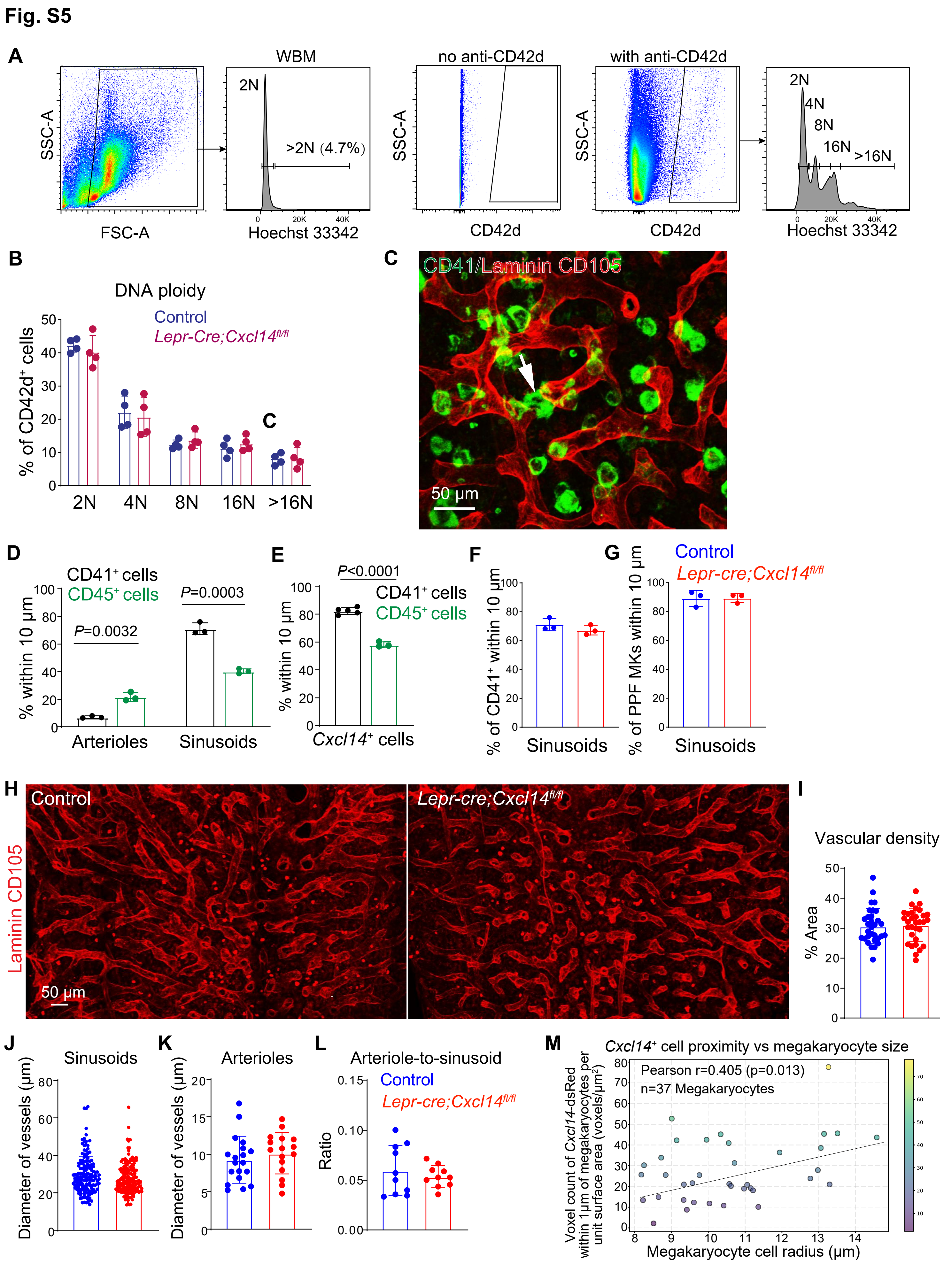

Fig. S5
A
SSC-A
FSC-A
WBM
2N
>2N (4.7%)
Hoechst 33342
no anti-CD42d
CD42d
with anti-CD42d
4N
8N
16N
>16N
B
DNA ploidy
Control
Lepr-Cre;Cxcl14fl/fl
% of CD42d+ cells
C
CD41/Laminin CD105
50 µm
D
CD41+ cells
CD45+ cells
% within 10 µm
P=0.0032
P=0.0003
Arterioles
Sinusoids
E
P<0.0001
Cxcl14+ cells
F
% of CD41+ within 10 µm
G
% of PPF MKs within 10 µm
Lepr-cre;Cxcl14fl/fl
H
Laminin CD105
I
Vascular density
% Area
J
Diameter of vessels (µm)
K
Arterioles
L
Arteriole-to-sinusoid
Ratio
M
Cxcl14+ cell proximity vs megakaryocyte size
Pearson r=0.405 (p=0.013)
n=37 Megakaryocytes
Voxel count of Cxcl14-dsRed within 1µm of megakaryocytes per unit surface area (voxels/µm2)
Megakaryocyte cell radius (µm)

**Fig. S5, Related to Figure 3. Megakaryocyte ploidy and blood vessels in the bone marrow of *Lepr-cre; Cxcl14$^{fl/fl}$* and control mice**. (A, B) DNA content (ploidy) of CD42d$^{+}$ megakaryocytes from *Lepr-cre;Cxcl14$^{fl/fl}$* and control bone marrow. Cells were stained with anti-CD42d antibody and Hoechst 33342 to assess DNA content in whole bone marrow cells and CD42d$^{+}$ megakaryocytes (A). Distribution of DNA content in megakaryocytes from *Lepr-cre; Cxcl14$^{fl/fl}$* and control bone marrow (B) (4 mice per genotype from 3 independent experiments). (C) CD41$^{+}$ megakaryocytes and proplatelet-forming megakaryocytes (arrow) localized adjacent to sinusoids in femur bone marrow from both *Lepr-cre; Cxcl14 $^{fl/fl}$* and control mice (3 mice per genotype from 3 independent experiments). (C) Representative image of femur bone marrow showing CD41$^{+}$ proplatelet-forming megakaryocytes (arrow) and sinusoidal blood vessels (Laminin/CD105). (D) CD41$^{+}$ megakaryocytes were significantly more likely than all CD45$^{+}$ cells to localize adjacent to sinusoids and less likely to localize adjacent to arterioles (a total of 3 mice per group from 3 independent experiments). (E) CD41$^{+}$ megakaryocytes were significantly more likely than all CD45$^{+}$ cells to localize adjacent to *Cxcl14*-dsRed$^{+}$ stromal cells (3 or 5 mice per group from 3 independent experiments). (F-G) *Cxcl14* deletion did not significantly alter the localization of CD41$^{+}$ megakaryocytes or proplatelet-forming megakaryocytes relative to sinusoids in femur bone marrow (3 mice per genotype from 3 independent experiments). (H-L) *Cxcl14* deletion did not significantly alter the vasculature in femur bone marrow of *Lepr-cre;Cxcl14$^{fl/fl}$* as compared to control mice: representative images (H), vascular density (I), sinusoid and arteriole diameter (J, K) and arteriole-to-sinusoid ratio (L) (3 mice per genotype from 3 independent experiments). (M) Linear regression fit of the number of *Cxcl14*$^{+}$ voxels per unit surface area of megakaryocytes within 1µm of the megakaryocyte cell surface versus megakaryocyte size (n = 37 cells; Pearson r = 0.405, p = 0.013). Each dot represents a different mouse in panels B and D-G. Data represent mean ± standard deviation. All statistical tests were two-sided. Statistical significance was assessed using Student's *t*-tests (B and D) or Mann-

Whitney tests (E) followed by Holm-Sidak’s multiple comparisons adjustments, a matched samples two-way ANOVA (F and G) or a two-way ANOVA (J and K) followed by Sidak's multiple comparisons adjustments, a Student’s *t*-test (I), or a Welch’s *t*-test (L).

### Fig. S6

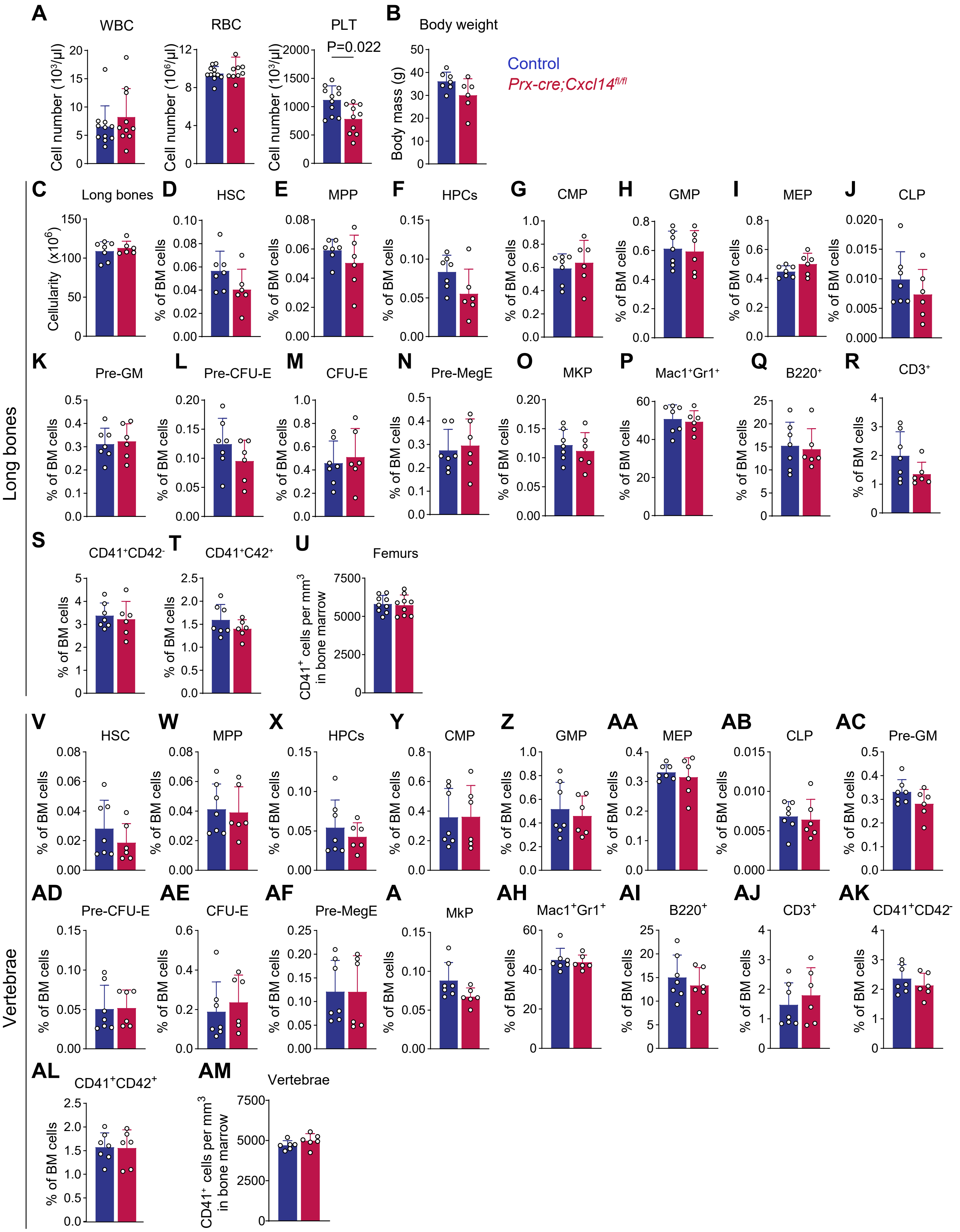

Control
Prx-cre;Cxcl14fl/fl
A
WBC
RBC
PLT
P=0.022
Cell number (10³/µl)
Cell number (10⁶/µl)
B
Body weight
Body mass (g)
C
Long bones
Cellularity (x10⁶)
D
HSC
E
MPP
F
HPCs
G
CMP
H
GMP
I
MEP
J
CLP
K
Pre-GM
L
Pre-CFU-E
M
CFU-E
N
Pre-MegE
O
MKP
P
Mac1⁺Gr1⁺
Q
B220⁺
R
CD3⁺
S
CD41⁺CD42⁻
T
CD41⁺C42⁺
U
Femurs
CD41⁺ cells per mm³ in bone marrow
% of BM cells
Long bones
V
HSC
W
MPP
X
HPCs
Y
CMP
Z
GMP
AA
MEP
AB
CLP
AC
Pre-GM
AD
Pre-CFU-E
AE
CFU-E
AF
Pre-MegE
A
MkP
AH
Mac1⁺Gr1⁺
AI
B220⁺
AJ
CD3⁺
AK
CD41⁺CD42⁻
AL
CD41⁺CD42⁺
AM
Vertebrae
Vertebrae

**Fig. S6, Related to Figure 3. Characterization of *Prx1-cre; Cxcl14$^{fl/fl}$* mice.** (A) WBC, RBC and PLT counts in the blood of *Prx1-cre; Cxcl14 $^{fl/fl}$* mice and littermate controls (10-11 mice per genotype from 3 independent experiments). (B) Body mass in *Prx1-cre; Cxcl14 $^{fl/fl}$* and littermate control mice at 6-9 months of age (6-7 mice per genotype from 2 independent experiments). (C-U) In the long bones (femurs and tibia) of *Prx1-cre; Cxcl14 $^{fl/fl}$* and littermate control mice we observed no differences in bone marrow cellularity (C) or the frequencies of HSCs (D), MPPs (E), HPCs (F), CMPs (G), GMPs (H), MEPs (I), CLPs (J), Pre-GMs (K), Pre-CFUEs (L), CFU-Es (M), Pre-MegEs (N), megakaryocyte progenitors (O), Mac-1$^+$Gr-1$^+$ myeloid cells (P), B220$^+$IgM$^+$ B cells (Q), CD3$^+$ T cells (R) CD41$^+$CD42d$^-$ megakaryocytes (S), CD41$^+$CD42d$^+$ megakaryocytes (T), or the numbers of CD41$^+$ megakaryocytes per mm$^3$ in bone marrow sections (U) (6-9 mice per genotype from 2-4 independent experiments). (V-AM) We also did not observe any significant differences in these hematopoietic parameters in the vertebral bone marrow of *Prx1-cre; Cxcl14 $^{fl/fl}$* and littermate control mice (6-7 mice per genotype from 2-3 independent experiments). All data represent mean ± standard deviation. Each dot represents a different mouse. All statistical tests were two-sided. Statistical significance was assessed using Student's *t*-tests followed by Holm-Sidak's multiple comparisons adjustments (A and D-AM) or Mann-Whitney tests followed by Holm-Sidak's multiple comparisons adjustments (A), or Student's *t*-tests (B and C).

**Fig. S7**

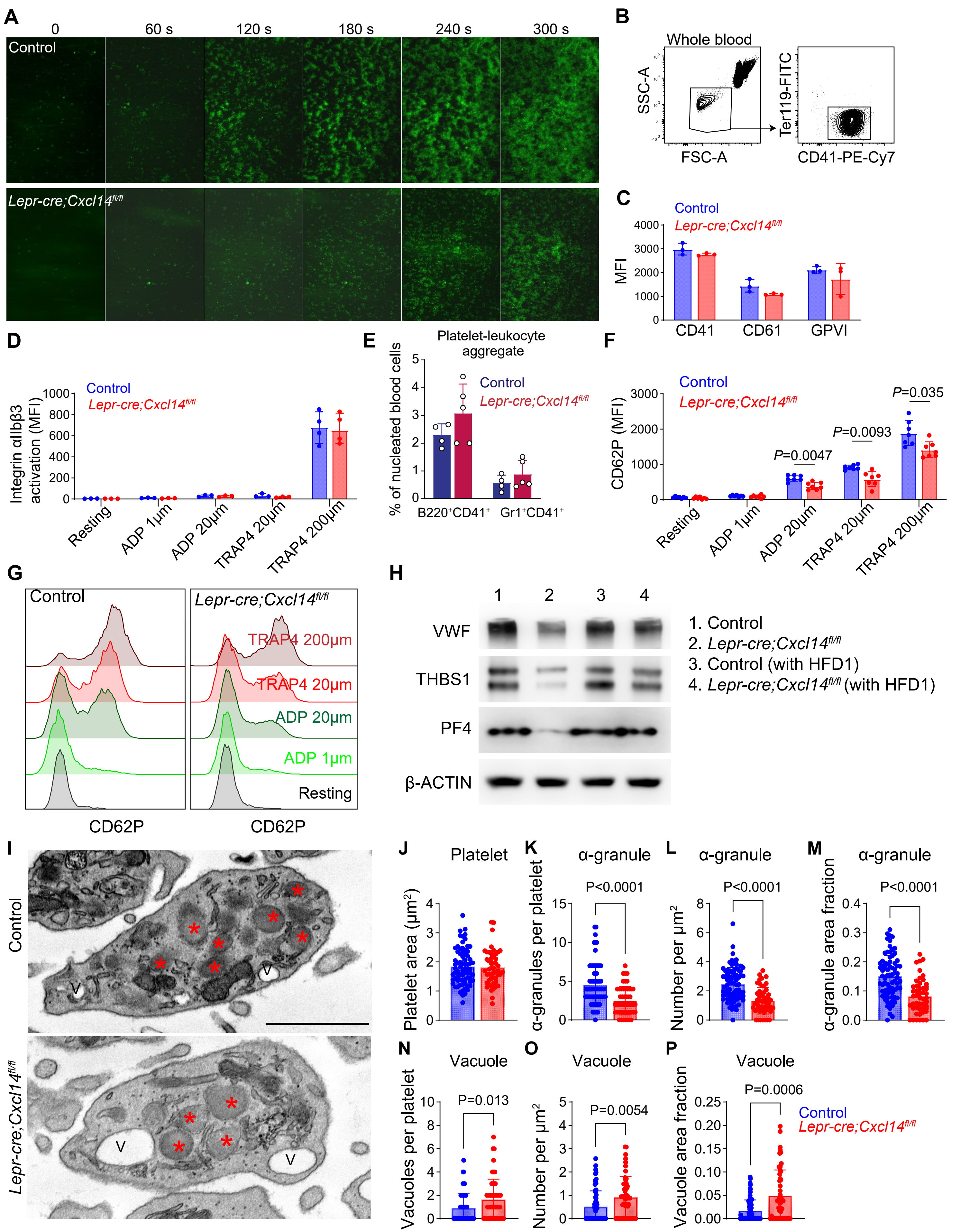
A
0
60 s
120 s
180 s
240 s
300 s
Control
Lepr-cre;Cxcl14fl/fl
B
Whole blood
SSC-A
FSC-A
Ter119-FITC
CD41-PE-Cy7
C
Control
Lepr-cre;Cxcl14fl/fl
MFI
CD41
CD61
GPVI
D
Control
Lepr-cre;Cxcl14fl/fl
Integrin αIIbβ3 activation (MFI)
Resting
ADP 1μm
ADP 20μm
TRAP4 20μm
TRAP4 200μm
E
Platelet-leukocyte aggregate
% of nucleated blood cells
Control
Lepr-cre;Cxcl14fl/fl
B220+CD41+
Gr1+CD41+
F
Control
Lepr-cre;Cxcl14fl/fl
CD62P (MFI)
P=0.0047
P=0.0093
P=0.035
Resting
ADP 1μm
ADP 20μm
TRAP4 20μm
TRAP4 200μm
G
Control
Lepr-cre;Cxcl14fl/fl
TRAP4 200μm
TRAP4 20μm
ADP 20μm
ADP 1μm
Resting
CD62P
CD62P
H
1
2
3
4
VWF
THBS1
PF4
β-ACTIN
1. Control
2. Lepr-cre;Cxcl14fl/fl
3. Control (with HFD1)
4. Lepr-cre;Cxcl14fl/fl (with HFD1)
I
Control
Lepr-cre;Cxcl14fl/fl
J
Platelet
Platelet area (μm²)
K
α-granule
P<0.0001
α-granules per platelet
L
α-granule
P<0.0001
Number per μm²
M
α-granule
P<0.0001
α-granule area fraction
N
Vacuole
P=0.013
Vacuoles per platelet
O
Vacuole
P=0.0054
Number per μm²
P
Vacuole
P=0.0006
Vacuole area fraction
Control
Lepr-cre;Cxcl14fl/fl

**Fig. S7, Related to Figure 3. Platelet function in *Lepr-cre; Cxcl14$^{fl/fl}$* and control mice.** (A) Representative images of platelet adhesion and thrombus formation under flow at the indicated time points on collagen-coated surfaces using blood from *Lepr-cre;Cxcl14$^{fl/fl}$* and control mice. (B) Representative flow cytometry gates used to obtain platelets from whole blood. (C) CD41, CD61, and GPVI mean fluorescence intensity (MFI) on platelets (3 mice per genotype from 2 independent experiments). (D) Integrin αIIbβ3 activation on platelets under resting conditions or after stimulation with adenosine diphosphate (ADP) (1 or 20 μM) or thrombin receptor–activating peptide 4 (TRAP4) (20 or 200 μM), assessed by flow cytometry (3-4 mice per genotype from 2 independent experiments). (E) Platelet aggregation with B220$^{+}$ B cells or Gr1$^{+}$ myeloid cells from the blood of *Lepr-cre;Cxcl14$^{fl/fl}$* or control mice (4-5 mice per genotype from 2 independent experiments). (F-G) Representative flow cytometry histograms (G) and quantification (F) of platelet CD62P expression under resting conditions or after stimulation with ADP (1 or 20 μM) or TRAP4 (20 or 200 μM) (7 mice per genotype from 3 independent experiments). (H) Western blot analysis of VWF, Thrombospondin1 (THBS1), and PF4 protein levels in washed platelets from *Lepr-cre;Cxcl14$^{fl/fl}$* or control mice fed normal chow or a high fat diet (HFD1). β-ACTIN was used as a loading control. (I) Representative transmission electron microscopy images of platelets from *Lepr-cre;Cxcl14$^{fl/fl}$* and control mice. Red asterisks indicate α-granules and V indicates vacuoles. Scale bar, 1 μm. (J-P) Quantification of platelet ultrastructural features from electron microscopy images including platelet area (J), number of α-granules per platelet (K), α-granule number per μm² of platelet area (L), α-granule area fraction (M), number of vacuoles per platelet (N), vacuole number per μm² of platelet area (O), and vacuole area fraction (P) (3 mice per genotype from 3 independent experiments; each dot represents a megakaryocyte; 50 or 76 platelets were analyzed per group). All data represent mean ± standard deviation. In panels C-F, each dot represents a different mouse. All statistical tests were two-sided. Statistical significance was assessed using a two-way ANOVA (C) or a matched samples two-way ANOVA

(E) followed by Sidak's multiple comparisons adjustments, Student's *t*-tests followed by Holm-Sidak's multiple comparisons adjustments (D, F), Mann-Whitney tests followed by Holm-Sidak's multiple comparisons adjustments (F and K-P), or a Student's *t*-test (J).

**Fig. S8**

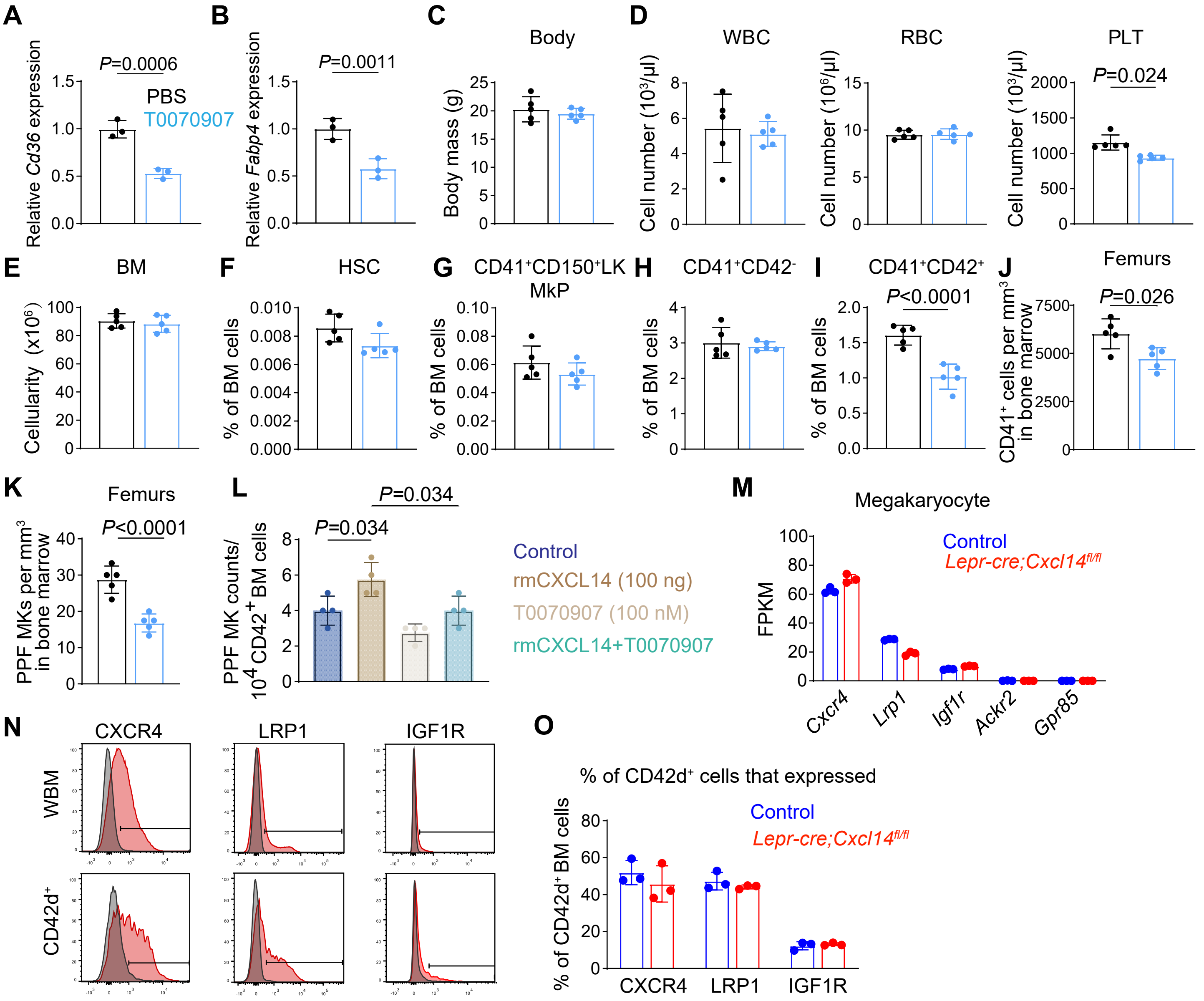
A
B
C
D
P=0.0006
PBS
T0070907
Relative Cd36 expression
P=0.0011
Relative Fabp4 expression
Body
Body mass (g)
WBC
Cell number (10³/µl)
RBC
Cell number (10⁶/µl)
PLT
P=0.024
Cell number (10³/µl)
E
F
G
H
I
J
BM
Cellularity (x10⁶)
HSC
% of BM cells
CD41⁺CD150⁺LK MkP
% of BM cells
CD41⁺CD42⁻
% of BM cells
CD41⁺CD42⁺
P<0.0001
% of BM cells
Femurs
P=0.026
CD41⁺ cells per mm³ in bone marrow
K
Femurs
P<0.0001
PPF MKs per mm³ in bone marrow
L
P=0.034
P=0.034
PPF MK counts/ 10⁴ CD42⁺ BM cells
Control
rmCXCL14 (100 ng)
T0070907 (100 nM)
rmCXCL14+T0070907
M
Megakaryocyte
FPKM
Control
Lepr-cre;Cxcl14fl/fl
Cxcr4
Lrp1
Igf1r
Ackr2
Gpr85
N
CXCR4
LRP1
IGF1R
WBM
CD42d⁺
O
% of CD42d⁺ cells that expressed
% of CD42d⁺ BM cells
Control
Lepr-cre;Cxcl14fl/fl
CXCR4
LRP1
IGF1R

**Fig. S8, Related to Figure 4. PPARγ inhibition impairs megakaryocyte maturation.** (A-B) *Cd36* and *Fabp4* expression by quantitative reverse-transcription PCR in $CD42^+$ bone marrow cells cultured for 12 hours with T0070907 (PPARγ inhibitor) or PBS control (a total of 3 cultures per group from 3 independent experiments). (C-K) Mice were intraperitoneally injected daily with T0070907 (1 mg/kg,day) or PBS for one week: body mass (C), WBC, RBC and PLT counts in the blood (D), bone marrow cellularity (E), frequencies of HSCs (F), megakaryocyte progenitors (G), $CD41^+CD42^-$ cells (H) and $CD41^+CD42^+$ cells (I) in the bone marrow by flow cytometry, as well as the numbers of $CD41^+$ megakaryocytes (J) and proplatelet-forming megakaryocytes (K) in bone marrow sections by immunofluorescence analysis (5 mice per genotype from 2 independent experiments). (L) $CD42^+$ bone marrow cells were cultured for 10 hours with T0070907 (100 nM) and/or recombinant mouse CXCL14 (rmCXCL14; 100 ng/ml) then proplatelet-forming (PPF) megakaryocytes were counted (a total of 4 samples per group from 2 independent experiments). (M) Expression of potential CXCL14 receptors, including *Cxcr4*, *Lrp1*, *Igf1r*, *Ackr2* and *Gpr85*, by RNA sequencing analysis of megakaryocytes from *Lepr-cre; Cxcl14*$^{fl/fl}$ and control mice (RNAseq from Figure 4) (3 mice per genotype from 3 independent experiments). (N) Flow-cytometric analysis of CXCR4, LRP1 and IGF1R staining on the surface of whole bone marrow cells or $CD42^+$ bone marrow cells. (O) The percentages of $CD42^+$ bone marrow cells that were $CXCR4^+$, $LRP1^+$ or $IGF1R^+$ (3 mice per genotype from 2 independent experiments). All data represent mean ± standard deviation and each dot represents a different mouse. All statistical tests were two-sided. Statistical significance was assessed using matched samples two-way ANOVAs followed by Sidak's multiple comparisons adjustments (A, B, F-I, O, J and K), Student's *t*-tests (C and E), Students' *t*-tests or Mann-Whitney test followed by Holm-Sidak's multiple comparisons adjustment (D), or a one-way ANOVA followed by Sidak's multiple comparisons adjustment (L).

**Fig. S9**

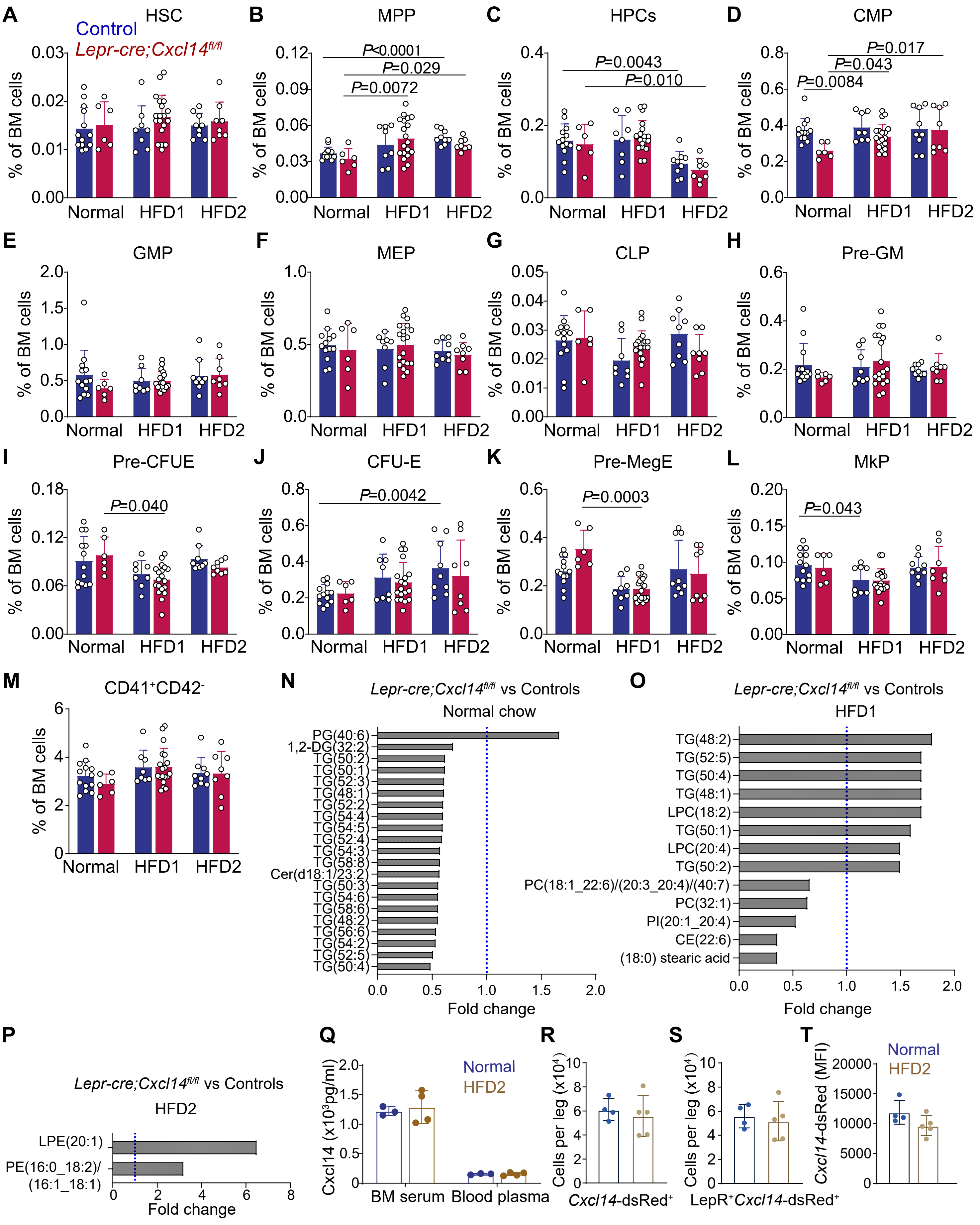
A
HSC
Control
Lepr-cre;Cxcl14fl/fl
% of BM cells
Normal
HFD1
HFD2
B
MPP
P<0.0001
P=0.029
P=0.0072
C
HPCs
P=0.0043
P=0.010
D
CMP
P=0.017
P=0.043
P=0.0084
E
GMP
F
MEP
G
CLP
H
Pre-GM
I
Pre-CFUE
P=0.040
J
CFU-E
P=0.0042
K
Pre-MegE
P=0.0003
L
MkP
P=0.043
M
CD41+CD42-
N
Lepr-cre;Cxcl14fl/fl vs Controls
Normal chow
PG(40:6)
1,2-DG(32:2)
TG(50:2)
TG(50:1)
TG(52:3)
TG(48:1)
TG(52:2)
TG(54:4)
TG(54:5)
TG(52:4)
TG(54:3)
TG(58:8)
Cer(d18:1/23:2)
TG(50:3)
TG(54:6)
TG(58:6)
TG(48:2)
TG(56:6)
TG(54:2)
TG(52:5)
TG(50:4)
Fold change
O
Lepr-cre;Cxcl14fl/fl vs Controls
HFD1
TG(48:2)
TG(52:5)
TG(50:4)
TG(48:1)
LPC(18:2)
TG(50:1)
LPC(20:4)
TG(50:2)
PC(18:1_22:6)/(20:3_20:4)/(40:7)
PC(32:1)
PI(20:1_20:4)
CE(22:6)
(18:0) stearic acid
Fold change
P
Lepr-cre;Cxcl14fl/fl vs Controls
HFD2
LPE(20:1)
PE(16:0_18:2)/
(16:1_18:1)
Fold change
Q
Cxcl14 (x10³pg/ml)
Normal
HFD2
BM serum
Blood plasma
R
Cells per leg (x10⁴)
Cxcl14-dsRed+
S
Cells per leg (x10⁴)
LepR+Cxcl14-dsRed+
T
Cxcl14-dsRed (MFI)
Normal
HFD2

**Fig. S9, Related to Figure 5. The frequencies of hematopoietic stem and progenitor cells did not significantly differ between *Lepr-cre; Cxcl14$^{fl/fl}$* and control mice, irrespective of whether they were fed normal chow or a high fat diet.** (A-M) *Lepr-cre; Cxcl14$^{fl/fl}$* and littermate control mice were fed normal chow (10% of calories from fat) or either of two high fat diets (HFD1 and HFD2, both of which had 60% of calories from fat) for four weeks. The frequencies of HSCs (A), MPPs (B), HPCs (C), CMPs (D), GMPs (E), MEPs (F), CLPs (G), Pre-GMs (H), Pre-CFUEs (I), CFU-Es (J), Pre-MegEs (K), megakaryocyte progenitors (L), and $CD41^+CD42d^-$ megakaryocytes (M) from the bone marrow of *Lepr-cre; Cxcl14$^{fl/fl}$* and control mice (a total of 6-19 mice per genotype per diet from 2-3 independent experiments). (N-P) Lipidomic analysis of $CD42d^+$ megakaryocytes from *Lepr-cre; Cxcl14$^{fl/fl}$* versus control bone marrow from mice fed normal chow (N), high fat diet 1 (O) or high fat diet 2 (P). Each panel shows all lipid species that significantly differed (FDR<0.05 and log2 FC>0.5 in either direction) between megakaryocytes from *Lepr-cre; Cxcl14$^{fl/fl}$* as compared to control bone marrow in mice fed each diet (a total of 3-13 mice per genotype in 2 experiments plus 2 experiments with mice on normal chow from the experiment shown in Figure 4H). (Q) ELISA analysis of the levels of CXCL14 in bone marrow serum and blood plasma from 5-month-old wild-type mice after one month of normal chow or a high fat diet (HFD2) (a total of 3-4 mice per diet from 2 independent experiments). (R-T) Number of *Cxcl14*-dsRed$^+$ stromal cells (R) or LepR$^+$*Cxcl14*-dsRed$^+$ cells (S) in the bone marrow per leg (femur and tibia) and mean *Cxcl14*-dsRed fluorescence intensity per LepR$^+$ cell (T) in *Cxcl14*-dsRed mice fed normal chow or a high fat diet (a total of 4-5 mice per diet from 2 independent experiments). All data represent mean ± standard deviation and each dot represents a different mouse. All statistical tests were two-sided. Statistical significance was assessed using two-way ANOVAs followed by Sidak's or Dunnett's multiple comparisons adjustments (A, C-E, G, L-M and R-T), Welch's one-way ANOVAs followed by Dunnett's T3 multiple comparisons adjustments (B, I, and J), Kruskal-Wallis tests followed by Dunn's multiple comparisons adjustments (F,H, and K), Omics Data Analyzer (see lipidomics methods; N-P), or a matched samples two-way ANOVA followed by Sidak's multiple comparisons adjustment (Q).

**Table S1. Markers used to identify each cell population by flow cytometry.**

| Cell population | Abbreviation | Markers | Ref |
|---|---|---|---|
| Hematopoietic stem cells | HSC | $CD150^{+}CD48^{-}Lin^{-}Sca1^{+}c\text{-}kit^{+}$ | (16) |
| Multipotent progenitors | MPP | $CD150^{-}CD48^{-}Lin^{-}Sca1^{+}c\text{-}kit^{+}$ | (16) |
| Hematopoietic progenitor cells | HPC | $CD48^{+}Lin^{-}Sca1^{+}c\text{-}kit^{+}$ | (17) |
| Common myeloid progenitors | CMP | $Lin^{-}$ $Sca1^{-}c\text{-}kit^{+}CD34^{+}CD16/32^{-}$ | (18) |
| Common lymphoid progenitors | CLP | $Lin^{-}$ $Sca1^{low}c\text{-}kit^{low}CD135^{+}CD127^{-}$ | |
| Granulocyte-macrophage progenitors | GMP | $Lin^{-}$ $Sca1^{-}c\text{-}kit^{+}CD34^{+}$ $CD16/32^{-}$ | (18) |
| Megakaryocyte-erythrocyte progenitors | MEP | $Lin^{-}$ $Sca1^{-}c\text{-}kit^{+}CD34^{-}CD16/32^{-}$ | (18) |
| Pre-Megakaryocyte-erythrocyte progenitors | PreMegE | $Lin^{-}$ $Sca1^{-}c\text{-}kit^{+}CD41^{-}$ $CD16/32^{-}$ $CD150^{+}CD105^{-}$ | (19) |
| Pre-Megakaryocyte-erythrocyte progenitors | PreCFU-E | $Lin^{-}$ $Sca1^{-}c\text{-}kit^{+}CD41^{-}$ $CD16/32^{-}$ $CD150^{+}CD105^{+}$ | (19) |
| CFU-E progenitors | CFU-E | $Lin^{-}c\text{-}kit^{+}Sca1^{-}CD41^{-}$ $CD16/32^{-}$ $CD150^{-}CD105^{+}$ | (19) |
| Megakaryocyte progenitors | MkP | $Lin^{-}c\text{-}kit^{+}Sca1^{-}CD41^{-}CD150^{+}$ | (19) |
| Pre-proB precursors | Pre-proB | $B220^{+}IgM^{-}CD24^{-}CD43^{+}$ | |
| ProB precursors | ProB | $B220^{+}IgM^{-}CD24^{+}CD43^{+}$ | |
| PreB precursors | PreB | $B220^{+}IgM^{-}CD24^{+}CD43^{-}$ | |
| $CD41^{+}CD42d^{-}$ megakaryocytes | $CD41^{+}CD42d^{-}$Mk | $CD41^{+}CD42d^{-}$ | |
| $CD41^{+}CD42d^{+}$ megakaryocytes | $CD41^{+}CD42d^{+}$Mk | $CD41^{+}CD42d^{+}$ | |
| Myeloid cells | $Mac1^{+}Gr1^{+}$ | $Mac1(CD11b)^{+}$ $Gr1^{+}$ | |
| Erythroid cells | $Ter119^{+}$ | $Ter119^{+}$ | |
| B cells | $B220^{+}$ | $B220^{+}$ | |
| T cells | $CD3^{+}$ | $CD3^{+}$ | |
| $LepR^{+}$ cells | $LepR^{+}$ cells | $CD45^{-}Ter119^{-}CD31^{-}LepR^{+}$ | |
| Endothelial cells | EC | $CD45^{-}Ter119^{-}CD31^{+}$ | |

**Table S2. Lipid species detected in megakaryocytes from the bone marrow of *Lepr-cre; Cxcl14*$^{fl/fl}$ and littermate control mice (a total of 7-9 mice per genotype from 2 independent experiments)**. Fold change (FC) reflects values from megakaryocytes from *Lepr-cre; Cxcl14* $^{fl/fl}$ mice divided by megakaryocytes from control mice. Lipid species highlighted in red have an FDR < 0.05 and a log2 FC > 0.5 in either direction.

| **Lipid species** | **Control** | | ***Lepr-cre;Cxcl14***$^{fl/fl}$ | | | |
|---|---|---|---|---|---|---|
| | **Log2-mean** | **Log2-SD** | **Log2-mean** | **Log2-SD** | **FDR** | **FC** |
| **Free Fatty acids (FFA)** | | | | | | |
| (16:0) palmitic acid | 21.4 | 0.764 | 21.7 | 0.704 | 0.32 | 1.2 |
| (18:0) stearic acid | 21.3 | 0.696 | 21.9 | 0.805 | 0.033 | 1.6 |
| (18:1) oleic acid | 18.8 | 0.246 | 19.2 | 1.4 | 0.25 | 1.4 |
| (18:2) linoleic acid | 16 | 0.845 | 16.1 | 0.808 | 0.93 | 1.1 |
| (20:4) arachidonic acid | 17.8 | 0.378 | 18.5 | 1.31 | 0.075 | 1.7 |
| (21:0) heneicosanoic acid | 16.8 | 1.04 | 17.2 | 1.1 | 0.57 | 1.4 |
| (22:6) docosahexaenoic acid | 15 | 1.53 | 14.8 | 1.18 | 0.48 | 0.86 |
| **Diacylglycerols (DG)** | | | | | | |
| 1,2-DG(18:1/18:3)_(18:2/18:2) | 20.6 | 1.42 | 20.6 | 1.79 | 0.81 | 1 |
| 1,2-DG(20:4/16:0) | 21.3 | 1.19 | 21.4 | 0.901 | 0.85 | 1 |
| 1,2-DG(32:0) | 20.4 | 0.813 | 20.8 | 0.433 | 0.3 | 1.4 |
| 1,2-DG(32:1) | 20.2 | 1.51 | 20.1 | 1.31 | 0.41 | 0.91 |
| 1,2-DG(32:2) | 20 | 1.19 | 19.7 | 0.908 | 0.16 | 0.8 |
| 1,2-DG(34:0) | 22.4 | 0.353 | 23.1 | 0.778 | 0.16 | 1.5 |
| 1,2-DG(34:1) | 22.3 | 1.2 | 22.3 | 1.05 | 0.66 | 0.98 |
| 1,2-DG(34:2) | 22.8 | 1.2 | 22.7 | 1.2 | 0.57 | 0.94 |
| 1,2-DG(34:3) | 20.9 | 1.46 | 20.7 | 1.59 | 0.26 | 0.84 |
| 1,2-DG(36:0) | 24.2 | 0.351 | 24.8 | 0.802 | 0.13 | 1.6 |
| 1,2-DG(36:1) | 20.7 | 0.934 | 20.7 | 0.79 | 0.79 | 1 |
| 1,2-DG(36:2) | 22.5 | 1.19 | 22.6 | 1.12 | 0.89 | 1.1 |
| 1,2-DG(36:3) | 22.7 | 1.31 | 22.8 | 1.52 | 0.8 | 1 |
| 1,2-DG(38:1) | 26.2 | 0.921 | 26.5 | 0.961 | 0.77 | 1.2 |
| 1,2-DG(38:4) | 23.4 | 1.11 | 23.3 | 0.89 | 0.75 | 0.94 |
| 1,2-DG(38:5) | 21.1 | 1.15 | 21 | 0.908 | 0.62 | 0.91 |
| 1,2-DG(38:6) | 18.7 | 1.66 | 18.4 | 1.67 | 0.2 | 0.78 |
| 1,2-DG(40:6) | 18.4 | 1.18 | 18.3 | 0.935 | 0.77 | 0.91 |
| 1,2-DG(40:7) | 17.7 | 1.2 | 17.4 | 1.04 | 0.31 | 0.81 |
| **Cholesteryl esters (CE)** | | | | | | |
| CE(20:4) | 17.8 | 0.453 | 17.8 | 0.379 | 0.93 | 1 |
| CE(22:6) | 17.1 | 0.866 | 16.9 | 0.718 | 0.34 | 0.88 |
| **Ceramides (Cer)** | | | | | | |
| Cer(40:2) | 24.9 | 0.617 | 24.7 | 0.636 | 0.73 | 0.86 |
| Cer(d18:0/16:0) | 21.1 | 0.699 | 21.2 | 0.288 | 0.92 | 1.1 |

| | | | | | | |
|---|---|---|---|---|---|---|
| Cer(d18:0/18:0) | 19.7 | 0.485 | 20.1 | 0.319 | 0.14 | 1.4 |
| Cer(d18:0/20:0) | 19.6 | 0.872 | 20 | 1.06 | 0.61 | 1.3 |
| Cer(d18:0/22:0) | 22.6 | 0.627 | 22.5 | 0.67 | 0.93 | 0.91 |
| Cer(d18:0/22:1) | 23.2 | 0.998 | 22.9 | 0.948 | 0.83 | 0.84 |
| Cer(d18:0/24:0) | 22.6 | 0.678 | 22.3 | 0.637 | 0.65 | 0.82 |
| Cer(d18:0/24:1) | 22.7 | 0.678 | 22.6 | 0.615 | 0.92 | 0.91 |
| Cer(d18:0/26:0) | 17.6 | 1.6 | 17.7 | 1.86 | 0.51 | 1.1 |
| Cer(d18:0/26:1) | 16.8 | 0.748 | 16.5 | 0.604 | 0.66 | 0.83 |
| Cer(d18:0/26:2)/(18:2/26:0) | 19.5 | 1.16 | 19.1 | 1.44 | 0.77 | 0.77 |
| Cer(d18:1/16:0) | 23.5 | 0.519 | 23.3 | 0.39 | 0.51 | 0.86 |
| Cer(d18:1/18:0) | 22.1 | 0.688 | 21.9 | 0.478 | 0.65 | 0.85 |
| Cer(d18:1/18:1) | 18.4 | 1.19 | 18.2 | 0.774 | 0.16 | 0.86 |
| Cer(d18:1/20:0) | 19.7 | 1.08 | 19.8 | 0.889 | 0.85 | 1.1 |
| Cer(d18:1/22:0) | 21.3 | 1.02 | 21.5 | 0.883 | 0.89 | 1.1 |
| Cer(d18:1/23:0) | 20.1 | 0.864 | 20 | 0.807 | 0.93 | 0.89 |
| Cer(d18:1/23:2) | 17.7 | 0.51 | 16.5 | 0.64 | 0.041 | 0.45 |
| Cer(d18:1/23:2)/(d18:2/23:1) | 18.7 | 0.487 | 18.4 | 0.845 | 0.84 | 0.84 |
| Cer(d18:1/24:0) | 20.7 | 0.74 | 20.5 | 0.918 | 0.92 | 0.88 |
| Cer(d18:1/24:1)/(d18:2/24:0) | 24.5 | 0.332 | 24.4 | 0.401 | 0.89 | 0.94 |
| Cer(d18:1/25:0) | 17.8 | 1.71 | 17.8 | 1.56 | 0.66 | 1 |
| Cer(d18:1/26:0) | 17.8 | 1.59 | 17.9 | 1.64 | 0.68 | 1 |
| Cer(d18:1/26:1) | 17.1 | 0.958 | 17 | 0.741 | 0.47 | 0.94 |
| Cer(d18:2/16:0) | 18.4 | 1.03 | 17.5 | 1.28 | 0.033 | 0.54 |
| Cer(d18:2/18:0) | 17.5 | 1.28 | 17.4 | 0.983 | 0.52 | 0.93 |
| Cer(d18:2/23:0)/(18:1/23:1) | 21.5 | 0.979 | 21.1 | 1.07 | 0.57 | 0.77 |
| Cer(d18:2/24:1)(18:1/24:2) | 22.5 | 1.56 | 22.3 | 1.31 | 0.95 | 0.86 |
| **lyso-phosphatidylcholines (LPC)** | | | | | | |
| LPC(16:0) | 23.3 | 0.355 | 23.5 | 0.348 | 0.57 | 1.1 |
| LPC(18:1) | 20.9 | 0.368 | 20.9 | 0.321 | 0.89 | 1 |
| LPC(18:2) | 19.1 | 0.428 | 19.3 | 0.558 | 0.64 | 1.1 |
| LPC(20:0) | 17.4 | 0.624 | 17.6 | 0.833 | 0.84 | 1.1 |
| LPC(20:1) | 17.3 | 0.582 | 17.3 | 0.198 | 0.89 | 0.97 |
| LPC(20:4) | 17.9 | 0.38 | 17.8 | 0.573 | 0.92 | 0.95 |
| LPC(22:0) | 17.1 | 0.543 | 17.2 | 0.639 | 0.93 | 1 |
| LPC(22:6) | 15.7 | 1.24 | 15.5 | 1.04 | 0.49 | 0.91 |
| LPC(24:0) | 18.2 | 0.344 | 18.1 | 0.487 | 0.83 | 0.95 |
| **lyso-phosphatidylethanolamines (LPE)** | | | | | | |
| LPE(16:0) | 20 | 0.433 | 20.3 | 0.406 | 0.57 | 1.2 |
| LPE(18:0) | 21.5 | 0.484 | 21.8 | 0.328 | 0.32 | 1.3 |
| LPE(18:1) | 20.7 | 0.341 | 20.6 | 0.365 | 0.91 | 0.98 |
| LPE(18:2) | 16.7 | 0.652 | 16.1 | 0.927 | 0.4 | 0.69 |
| LPE(20:1) | 18.7 | 0.712 | 18.8 | 0.307 | 0.89 | 1.1 |
| LPE(20:4) | 19.1 | 0.551 | 19.1 | 0.689 | 0.91 | 1.1 |

| | | | | | | |
|---|---|---|---|---|---|---|
| LPE(22:4) | 17.9 | 0.459 | 18 | 0.433 | 0.83 | 1.1 |
| LPE(22:5) | 15.4 | 1.17 | 15.3 | 0.942 | 0.57 | 0.96 |
| LPE(22:6) | 18.3 | 0.881 | 18.2 | 1.01 | 0.83 | 0.93 |
| **lyso-phosphatidylinositol (LPI)** | | | | | | |
| LPI(18:0) | 17.2 | 0.875 | 17.8 | 0.861 | 0.22 | 1.5 |
| **Phosphatidylcholines (PC)** | | | | | | |
| PC(16:0_16:0) | 25.9 | 0.264 | 26.3 | 0.305 | 0.064 | 1.3 |
| PC(16:0_18:0) | 23.4 | 0.299 | 23.6 | 0.219 | 0.033 | 1.2 |
| PC(16:0_18:1) | 26.8 | 0.224 | 26.9 | 0.128 | 0.51 | 1.1 |
| PC(16:0_18:2) | 24.8 | 0.683 | 25.1 | 0.816 | 0.09 | 1.3 |
| PC(16:0_20:2)/(18:1_18:1)/(18:0_18:2) | 25.4 | 0.372 | 25.3 | 0.233 | 0.83 | 0.98 |
| PC(16:0_20:3) | 23 | 0.177 | 23.1 | 0.225 | 0.57 | 1.1 |
| PC(16:0_20:3)/(18:2_18:1) | 24 | 0.394 | 24.1 | 0.291 | 0.85 | 1.1 |
| PC(16:0_20:4) | 25.7 | 0.274 | 25.9 | 0.319 | 0.11 | 1.2 |
| PC(16:0_20:5) | 18.8 | 0.724 | 19 | 0.674 | 0.95 | 1.1 |
| PC(16:0_22:4) | 21.5 | 0.694 | 21.7 | 0.689 | 0.8 | 1.1 |
| PC(16:0_22:5)/(18:0_20:5)/(18:1_20:4) | 23.9 | 0.462 | 24.1 | 0.429 | 0.56 | 1.1 |
| PC(16:0_22:6) | 24.2 | 0.626 | 24.1 | 0.736 | 0.66 | 0.94 |
| PC(16:1_14:0)/(30:1) | 19.6 | 0.286 | 19.7 | 0.283 | 0.89 | 1 |
| PC(16:1_18:1) | 23.2 | 1.96 | 22.9 | 1.96 | 0.95 | 0.79 |
| PC(16:1_18:2) | 21.3 | 0.74 | 21.5 | 0.749 | 0.83 | 1.1 |
| PC(16:1_20:3)/(18:2_18:2) | 22.1 | 0.457 | 22.4 | 0.398 | 0.57 | 1.2 |
| PC(16:1_20:4) | 20.8 | 1.18 | 21.1 | 0.966 | 0.83 | 1.2 |
| PC(18:0_20:4) | 24.8 | 0.449 | 24.9 | 0.378 | 0.92 | 1.1 |
| PC(18:0_22:4) | 20.4 | 0.365 | 20.6 | 0.248 | 0.57 | 1.2 |
| PC(18:0_22:4)/(40:4) | 20.8 | 0.488 | 20.6 | 0.284 | 0.8 | 0.88 |
| PC(18:0_22:5) | 20.4 | 0.994 | 20.4 | 1.25 | 0.85 | 1 |
| PC(18:0_22:6) | 20.9 | 2.06 | 20.8 | 2.65 | 0.57 | 0.96 |
| PC(18:1_22:5) | 18.6 | 0.878 | 18.7 | 0.81 | 0.93 | 1.1 |
| PC(18:1_22:6) | 19.2 | 1.22 | 18.7 | 0.631 | 0.51 | 0.73 |
| PC(18:1_24:0)/(16:0_26:1) | 20.3 | 0.723 | 20.4 | 0.654 | 0.84 | 1.1 |
| PC(18:2_18:1) | 23.5 | 0.223 | 23.6 | 0.276 | 0.71 | 1.1 |
| PC(18:2_18:2) | 21.4 | 0.527 | 21.4 | 0.211 | 0.98 | 1 |
| PC(18:2_20:2) | 17.8 | 1.6 | 18.5 | 1.78 | 0.51 | 1.7 |
| PC(18:2_20:4) | 22.3 | 0.545 | 22.6 | 0.516 | 0.37 | 1.2 |
| PC(18:2_22:4) | 19.2 | 0.755 | 19.1 | 0.822 | 0.64 | 0.97 |
| PC(18:3_16:0) | 20.1 | 0.722 | 20.5 | 0.855 | 0.34 | 1.3 |
| PC(20:3_20:4) | 18.5 | 0.633 | 18.6 | 0.623 | 0.81 | 1.1 |
| PC(20:4_20:1) | 19.5 | 2.21 | 19.8 | 2.09 | 0.8 | 1.2 |
| PC(20:4_20:2) | 19.8 | 0.548 | 20.1 | 0.488 | 0.22 | 1.3 |
| PC(30:1) | 18.6 | 0.531 | 18.8 | 0.192 | 0.8 | 1.1 |
| PC(32:1) | 24.3 | 0.346 | 24.5 | 0.262 | 0.6 | 1.1 |

| | | | | | | |
|---|---|---|---|---|---|---|
| PC(32:2) | 20.8 | 0.744 | 21 | 0.747 | 0.81 | 1.1 |
| PC(36:0) | 22.2 | 1.33 | 22.2 | 1.07 | 0.42 | 1 |
| PC(36:1) | 25.1 | 0.415 | 25.1 | 0.149 | 0.83 | 0.94 |
| PC(38:2) | 22 | 0.484 | 22 | 0.212 | 0.93 | 0.99 |
| PC(40:2) | 20.3 | 0.812 | 20.2 | 0.323 | 0.92 | 0.91 |
| PC(40:5) | 21.3 | 0.651 | 21.3 | 0.75 | 0.29 | 0.99 |
| PC(40:7) | 21.3 | 0.792 | 21.1 | 1.04 | 0.33 | 0.84 |
| PC(40:8) | 21.2 | 0.624 | 21.3 | 0.581 | 0.91 | 1 |
| **Phosphatidylcholine-ether lipids (PC(O))** | | | | | | |
| PC(O-16:0) | 20.1 | 0.53 | 20.3 | 0.415 | 0.57 | 1.1 |
| PC(O-16:0_16:0) | 23.8 | 0.413 | 24.1 | 0.615 | 0.62 | 1.2 |
| PC(O-16:0_20:3) | 19.4 | 0.814 | 19.7 | 0.87 | 0.62 | 1.2 |
| PC(O-16:0_20:4) | 23.6 | 0.204 | 23.8 | 0.388 | 0.64 | 1.1 |
| PC(O-16:1_18:1) | 20.1 | 0.823 | 20.1 | 0.619 | 0.93 | 0.96 |
| PC(O-16:1_20:4) | 21 | 0.702 | 20.9 | 0.569 | 0.92 | 0.97 |
| PC-(O-18:0_18:2) | 21.9 | 0.237 | 22.3 | 1.08 | 0.79 | 1.3 |
| PC(O-18:0_18:3) | 21.5 | 0.381 | 21.7 | 0.728 | 0.6 | 1.2 |
| PC(O-18:1_22:6) | 19.4 | 0.567 | 19.4 | 0.3 | 0.92 | 0.94 |
| PC(O-20:0_16:0) | 19 | 1.1 | 18.6 | 0.906 | 0.65 | 0.8 |
| PC(O-30:0) | 19.8 | 0.406 | 20 | 0.628 | 0.66 | 1.2 |
| PC(O-32:1) | 21 | 0.702 | 21.1 | 0.965 | 0.64 | 1 |
| PC(O-32:2) | 18.1 | 1.66 | 17.9 | 1.62 | 1 | 0.83 |
| PC(O-34:0) | 21.4 | 0.434 | 21.6 | 0.63 | 0.62 | 1.1 |
| PC(O-34:1) | 24 | 0.318 | 24.1 | 0.524 | 0.67 | 1.1 |
| PC(O-34:2) | 22.4 | 0.296 | 22.7 | 0.792 | 0.65 | 1.2 |
| PC(O-36:1) | 19.6 | 0.76 | 19.8 | 0.778 | 0.91 | 1.1 |
| PC-(O-36:2) | 20.8 | 0.15 | 20.7 | 0.51 | 0.79 | 0.89 |
| PC(O-38:5) | 23.2 | 0.306 | 23.3 | 0.299 | 0.57 | 1.1 |
| PC(O-38:6) | 20 | 0.732 | 20.1 | 0.691 | 0.95 | 1.1 |
| PC(O-40:7) | 20.5 | 0.205 | 21 | 0.178 | 0.012 | 1.4 |
| **Plasmenylcholines (PC(P))** | | | | | | |
| PC(P-16:0_16:0) | 19.5 | 0.37 | 19.5 | 0.496 | 0.89 | 0.98 |
| PC(P-18:0_18:1) | 22.6 | 0.388 | 22.8 | 0.777 | 0.89 | 1.1 |
| **Phosphatidylethanolamines (PE)** | | | | | | |
| PE(16:0_16:1) | 18.7 | 0.941 | 18.8 | 0.82 | 0.92 | 1.1 |
| PE(16:0_18:0) | 19.2 | 0.177 | 19.4 | 0.202 | 0.34 | 1.1 |
| PE(16:0_18:2) | 22.1 | 0.111 | 22.3 | 0.148 | 0.064 | 1.2 |
| PE(16:0_18:2)/(16:1_18:1) | 21.8 | 0.392 | 22.1 | 0.264 | 0.66 | 1.2 |
| PE(16:0_20:2)/(18:1_18:1)/(18:0_18:2) | 24.7 | 0.433 | 24.5 | 0.219 | 0.62 | 0.88 |
| PE(16:0_20:3) | 20.4 | 0.0875 | 20.7 | 0.21 | 0.14 | 1.2 |
| PE(16:0_20:3)/(18:1_18:2) | 22.1 | 0.46 | 22.3 | 0.146 | 0.79 | 1.1 |
| PE(16:0_20:4) | 23.5 | 0.396 | 23.8 | 0.227 | 0.19 | 1.2 |
| PE(16:0_22:4) | 20.9 | 0.62 | 21.2 | 0.725 | 0.16 | 1.2 |

| | | | | | | |
|---|---|---|---|---|---|---|
| PE(16:0_22:5) | 21.2 | 0.387 | 21.3 | 0.315 | 0.43 | 1.1 |
| PE(16:0_22:6) | 23 | 0.198 | 23 | 0.307 | 0.83 | 0.98 |
| PE(16:1_18:1) | 19.6 | 0.296 | 19.4 | 0.308 | 0.51 | 0.84 |
| PE(16:1_20:4) | 20.1 | 0.546 | 20.1 | 0.419 | 0.99 | 1 |
| PE(18:0_20:3) | 19.2 | 0.327 | 19 | 0.255 | 0.62 | 0.91 |
| PE(18:0_20:4) | 25.7 | 0.308 | 25.7 | 0.151 | 0.89 | 1 |
| PE(18:0_22:4) | 22.5 | 0.135 | 22.6 | 0.155 | 0.32 | 1.1 |
| PE(18:0_22:5) | 21.8 | 0.357 | 21.8 | 0.204 | 0.92 | 1 |
| PE(18:0_22:6) | 24.6 | 0.643 | 24.3 | 0.521 | 0.09 | 0.81 |
| PE(18:1_18:2) | 22.4 | 0.0975 | 22.5 | 0.214 | 0.78 | 1.1 |
| PE(18:1_20:4) | 24.2 | 0.264 | 24.3 | 0.193 | 0.83 | 1.1 |
| PE(18:1_22:4) | 21.1 | 1.36 | 20.6 | 0.319 | 0.39 | 0.69 |
| PE(18:1_22:6) | 22.5 | 0.428 | 22.4 | 0.599 | 0.51 | 0.92 |
| PE(18:2_20:4) | 20.4 | 0.718 | 20.5 | 0.706 | 0.93 | 1.1 |
| PE(20:4_20:0) | 18.2 | 0.618 | 18.4 | 0.683 | 0.32 | 1.2 |
| PE(20:4_20:1) | 21 | 0.367 | 21.2 | 0.246 | 0.6 | 1.1 |
| PE(34:1) | 23.9 | 1.11 | 23.5 | 0.271 | 0.32 | 0.74 |
| PE(36:1) | 23.8 | 0.136 | 23.8 | 0.15 | 0.93 | 0.99 |
| PE(38:2) | 22.3 | 0.301 | 22.3 | 0.264 | 0.72 | 1 |
| PE(38:3) | 21.4 | 0.298 | 21.6 | 0.161 | 0.32 | 1.1 |
| PE(42:1) | 19 | 0.568 | 18.6 | 0.385 | 0.13 | 0.74 |
| **Phosphatidylethanolamines-ether lipids (PE (O))** | | | | | | |
| PE(O-16:0_18:1) | 20.7 | 0.309 | 20.8 | 0.443 | 0.52 | 1.1 |
| PE(O-16:1_22:4) | 23.7 | 0.374 | 23.8 | 0.342 | 0.62 | 1.1 |
| PE(O-18:0_18:2) | 19.9 | 0.761 | 19.9 | 1.12 | 0.69 | 1 |
| PE(O-18:1_18:1) | 18.7 | 0.781 | 18.9 | 0.952 | 0.78 | 1.2 |
| PE(O-38:5) | 24 | 0.148 | 24.3 | 0.203 | 0.19 | 1.2 |
| **Plasmenylethalomines (PE (P))** | | | | | | |
| PE(P-16:0_16:0) | 19.6 | 0.31 | 19.8 | 0.389 | 0.32 | 1.2 |
| PE(P-18:0_18:1) | 25.4 | 0.427 | 25.4 | 0.363 | 0.8 | 0.99 |
| PE(P-18:0_20:4) | 24.5 | 0.186 | 24.7 | 0.228 | 0.43 | 1.2 |
| PE(P-18:0_22:6) | 22.9 | 0.222 | 22.9 | 0.221 | 0.89 | 1 |
| **Phosphatidylglycerols (PG)** | | | | | | |
| PG(34:1) | 22.7 | 0.333 | 22.7 | 0.479 | 0.73 | 1 |
| PG(34:2) | 18.9 | 0.803 | 18.7 | 0.961 | 0.84 | 0.83 |
| PG(36:1) | 18.2 | 1.08 | 17.9 | 0.816 | 0.78 | 0.78 |
| PG(36:2) | 22 | 0.974 | 21.9 | 0.81 | 0.93 | 0.89 |
| PG(36:3) | 22.6 | 0.709 | 22.6 | 0.617 | 0.81 | 1 |
| PG(36:4) | 21.5 | 0.789 | 21.7 | 0.583 | 0.58 | 1.2 |
| PG(38:4) | 20.2 | 1 | 20.2 | 0.779 | 0.95 | 0.95 |
| PG(38:5) | 23.1 | 0.821 | 23 | 0.73 | 0.91 | 0.99 |
| PG(38:6) | 22.9 | 0.74 | 23 | 0.63 | 0.77 | 1.1 |
| PG(40:5) | 21.7 | 0.856 | 21.7 | 0.761 | 0.79 | 1.1 |

| | | | | | | |
|---|---|---|---|---|---|---|
| PG(40:6) | 21.7 | 0.88 | 22 | 0.788 | 0.41 | 1.2 |
| PG(40:7) | 20.7 | 1.37 | 21 | 1.13 | 0.65 | 1.2 |
| PG(40:8) | 22.8 | 0.642 | 22.8 | 0.581 | 0.89 | 1 |
| PG(40:9) | 16.9 | 0.865 | 16.8 | 0.885 | 0.92 | 0.91 |
| **Phosphatidylinositols (PI)** | | | | | | |
| PI(16:0_18:1) | 22.9 | 1.43 | 22.9 | 1.65 | 0.32 | 1 |
| PI(16:0_20:4) | 23.9 | 0.552 | 24.3 | 0.7 | 0.033 | 1.3 |
| PI(16:0_22:6) | 21.3 | 0.659 | 21.5 | 0.638 | 0.32 | 1.1 |
| PI(18:0_16:0) | 20.4 | 0.819 | 20.6 | 0.874 | 0.65 | 1.2 |
| PI(18:0_18:2) | 22 | 0.628 | 22.4 | 0.804 | 0.22 | 1.3 |
| PI(18:0_20:3) | 22.6 | 0.285 | 22.9 | 0.291 | 0.2 | 1.2 |
| PI(18:0_20:4) | 26.9 | 0.684 | 27.3 | 0.826 | 0.033 | 1.3 |
| PI(18:0_22:4) | 21.7 | 0.976 | 22 | 1.16 | 0.15 | 1.3 |
| PI(18:0_22:5) | 18.8 | 0.636 | 19.1 | 0.733 | 0.41 | 1.2 |
| PI(18:0_22:6) | 22.6 | 0.773 | 22.7 | 0.705 | 0.57 | 1.1 |
| PI(18:1_18:0) | 21.9 | 0.979 | 22.1 | 1.01 | 0.32 | 1.1 |
| PI(18:1_18:1) | 22.1 | 1.76 | 22.2 | 2.07 | 0.3 | 1.1 |
| PI(18:1_20:3) | 17.2 | 3.44 | 17.7 | 3 | 0.23 | 1.4 |
| PI(18:1_20:4) | 24.2 | 0.44 | 24.6 | 0.66 | 0.063 | 1.3 |
| PI(20:1_20:4) | 20.9 | 0.688 | 21.3 | 0.966 | 0.06 | 1.3 |
| PI(34:2) | 21.6 | 0.439 | 22 | 0.897 | 0.15 | 1.3 |
| PI(36:3) | 21.4 | 0.588 | 21.7 | 0.723 | 0.2 | 1.2 |
| **Sphingomyelins (SM)** | | | | | | |
| SM(18:0/16:0) | 22.1 | 0.571 | 22.5 | 0.404 | 0.37 | 1.3 |
| SM(18:0/18:0) | 21.1 | 0.491 | 20.9 | 0.281 | 0.32 | 0.85 |
| SM(18:0/20:0) | 20.4 | 0.459 | 20.3 | 0.226 | 0.79 | 0.92 |
| SM(18:0/22:0) | 21.6 | 0.702 | 21.5 | 0.295 | 0.95 | 0.94 |
| SM(18:0/22:2)/(18:1/22:1)/(18:2/22:0) | 21.3 | 0.325 | 21.6 | 0.158 | 0.17 | 1.3 |
| SM(18:0/24:1) | 22.5 | 0.655 | 22.5 | 0.418 | 0.81 | 0.99 |
| SM(18:1/14:0) | 18.1 | 0.271 | 18.1 | 0.695 | 0.79 | 0.95 |
| SM(18:1/15:0) | 19.1 | 0.386 | 19.4 | 0.416 | 0.32 | 1.3 |
| SM(18:1/16:0) | 25.2 | 0.432 | 25.6 | 0.425 | 0.24 | 1.3 |
| SM(18:1/17:0) | 18.3 | 0.47 | 18.5 | 0.264 | 0.33 | 1.2 |
| SM(18:1/18:0) | 23.8 | 0.373 | 23.9 | 0.203 | 0.66 | 1.1 |
| SM(18:1/20:0) | 22.4 | 0.629 | 22.4 | 0.262 | 0.73 | 1 |
| SM(18:1/21:0) | 18.8 | 0.533 | 18.9 | 0.403 | 0.77 | 1 |
| SM(18:1/22:0) | 23.9 | 0.706 | 24.1 | 0.429 | 0.19 | 1.2 |
| SM(18:1/22:1)/(18:2/22:0)/(18:0/22:2) | 22.5 | 0.28 | 22.5 | 0.234 | 0.95 | 1 |
| SM(18:1/23:0) | 21.5 | 0.773 | 21.6 | 0.571 | 0.57 | 1 |
| SM(18:1/23:1)/(18:2/23:0) | 20 | 0.479 | 20.1 | 0.371 | 0.51 | 1.1 |
| SM(18:1/24:0) | 24.2 | 0.786 | 24.3 | 0.483 | 0.32 | 1.1 |
| SM(18:1/24:1)_(18:2/24:0) | 25.5 | 0.704 | 25.7 | 0.509 | 0.17 | 1.1 |

| | | | | | | |
|---|---|---|---|---|---|---|
| SM(18:1/24:2)_(18:2/24:1) | 23.5 | 0.537 | 23.8 | 0.575 | 0.22 | 1.3 |
| SM(18:1/25:0)_(18:0/25:1) | 16 | 1.05 | 16.3 | 1.1 | 0.91 | 1.2 |
| SM(18:2/25:0)_(18:1/25:1) | 18.5 | 0.856 | 18.2 | 0.662 | 0.57 | 0.84 |
| SM(36:2) | 19.4 | 0.302 | 19.4 | 0.219 | 0.95 | 1 |
| SM(40:3) | 17.6 | 0.701 | 17.3 | 0.742 | 0.72 | 0.81 |
| **Triglycerides (TG)** | | | | | | |
| TG(48:1) | 27 | 0.746 | 26.4 | 0.644 | 0.19 | 0.66 |
| TG(48:2) | 27.4 | 0.797 | 26.5 | 0.729 | 0.063 | 0.56 |
| TG(50:0) | 24.2 | 1.02 | 24 | 0.531 | 0.79 | 0.82 |
| TG(50:1) | 27.8 | 0.683 | 27.2 | 0.448 | 0.14 | 0.66 |
| TG(50:2) | 28.6 | 0.63 | 28 | 0.548 | 0.072 | 0.64 |
| TG(50:3) | 28.3 | 0.717 | 27.5 | 0.591 | 0.033 | 0.56 |
| TG(50:4) | 26.8 | 0.816 | 25.7 | 0.709 | 0.033 | 0.47 |
| TG(52:1) | 26.2 | 0.797 | 25.5 | 0.597 | 0.26 | 0.62 |
| TG(52:2) | 28.5 | 0.589 | 27.8 | 0.444 | 0.05 | 0.62 |
| TG(52:3) | 29 | 0.542 | 28.3 | 0.452 | 0.033 | 0.63 |
| TG(52:4) | 28.5 | 0.566 | 27.7 | 0.402 | 0.033 | 0.6 |
| TG(52:5) | 26.9 | 0.664 | 25.9 | 0.561 | 0.033 | 0.51 |
| TG(54:2) | 26.4 | 0.602 | 25.5 | 0.572 | 0.064 | 0.55 |
| TG(54:3) | 28.1 | 0.542 | 27.3 | 0.443 | 0.033 | 0.59 |
| TG(54:4) | 28.3 | 0.492 | 27.6 | 0.421 | 0.033 | 0.62 |
| TG(54:5) | 27.7 | 0.532 | 27 | 0.444 | 0.05 | 0.6 |
| TG(54:6) | 26.5 | 0.6 | 25.6 | 0.508 | 0.033 | 0.53 |
| TG(56:6) | 25.1 | 0.558 | 24.1 | 0.466 | 0.017 | 0.52 |
| TG(58:10) | 20.6 | 0.665 | 19.8 | 0.565 | 0.07 | 0.55 |
| TG(58:6) | 22.5 | 0.539 | 21.6 | 0.476 | 0.012 | 0.53 |
| TG(58:8) | 22.9 | 0.535 | 22.1 | 0.476 | 0.033 | 0.55 |
| TG(60:10) | 19.2 | 0.464 | 18.6 | 0.748 | 0.22 | 0.62 |
| TG(60:11) | 18.5 | 0.629 | 17.6 | 0.915 | 0.14 | 0.51 |
| TG(60:12) | 16.6 | 1.52 | 15.9 | 0.671 | 0.34 | 0.59 |

**Table S3. Lipid species detected in megakaryocytes from the bone marrow of *Lepr-cre; Cxcl14*$^{fl/fl}$ and littermate control mice fed normal chow in the experiments in Figure 5 (a total of 11-13 mice per genotype in 2 independent experiments plus 2 additional experiments with mice fed normal chow from the experiment shown in Figure 4h)**. Fold change (FC) reflects values from megakaryocytes from *Lepr-cre; Cxcl14* $^{fl/fl}$ mice divided by megakaryocytes from control mice. Lipid species highlighted in red have an FDR < 0.05 and a log2 FC > 0.5 in either direction.

| **Lipid species** | **Control** | | ***Lepr-cre;Cxcl14***$^{fl/fl}$ | | | |
|---|---|---|---|---|---|---|
| | **Log2-mean** | **Log2-SD** | **Log2-mean** | **Log2-SD** | **FDR** | **FC** |
| **Free Fatty acids (FFA)** | | | | | | |
| (16:0) palmitic acid | 21.1 | 0.72 | 21.4 | 0.693 | 0.38 | 1.2 |
| (18:0) stearic acid | 21 | 0.821 | 21.5 | 0.952 | 0.15 | 1.4 |
| (18:1) oleic acid | 18.7 | 0.349 | 19 | 1.13 | 0.39 | 1.2 |
| (18:2) linoleic acid | 15.9 | 1.05 | 15.9 | 0.902 | 0.78 | 1 |
| (20:4) arachidonic acid | 17.7 | 0.302 | 18.2 | 1.12 | 0.15 | 1.4 |
| (21:0) heneicosanoic acid | 16.4 | 0.933 | 16.8 | 1.1 | 0.61 | 1.3 |
| (22:6) docosahexaenoic acid | 14.7 | 1.42 | 14.6 | 1.2 | 0.57 | 0.94 |
| **Diacylglycerols (DG)** | | | | | | |
| 1,2-DG(18:1/18:3)_(18:2/18:2) | 21 | 1.35 | 20.9 | 1.63 | 0.68 | 0.92 |
| 1,2-DG(20:4/16:0) | 21.5 | 1.02 | 21.5 | 0.929 | 0.87 | 0.99 |
| 1,2-DG(32:0) | 20.5 | 0.803 | 20.9 | 0.514 | 0.2 | 1.3 |
| 1,2-DG(32:1) | 20.7 | 1.43 | 20.4 | 1.32 | 0.24 | 0.82 |
| 1,2-DG(32:2) | 20 | 1.58 | 19.5 | 1.4 | 0.045 | 0.7 |
| 1,2-DG(34:0) | 22.5 | 0.695 | 23 | 0.689 | 0.069 | 1.4 |
| 1,2-DG(34:1) | 22.7 | 1.2 | 22.6 | 1.17 | 0.66 | 0.93 |
| 1,2-DG(34:2) | 23.2 | 1.19 | 22.9 | 1.19 | 0.3 | 0.84 |
| 1,2-DG(34:3) | 21.2 | 1.27 | 20.9 | 1.42 | 0.15 | 0.8 |
| 1,2-DG(36:0) | 24.2 | 0.769 | 24.8 | 0.77 | 0.052 | 1.5 |
| 1,2-DG(36:1) | 21.2 | 1.04 | 21.2 | 1.03 | 0.92 | 1 |
| 1,2-DG(36:2) | 23.1 | 1.36 | 23.1 | 1.35 | 0.89 | 0.97 |
| 1,2-DG(36:3) | 23.2 | 1.36 | 23.1 | 1.49 | 0.68 | 0.92 |
| 1,2-DG(38:0) | 12.2 | 2.22 | 12.8 | 2.84 | 0.68 | 1.6 |
| 1,2-DG(38:1) | 19.4 | 5.98 | 21.3 | 5.93 | 0.22 | 3.9 |
| 1,2-DG(38:4) | 23.5 | 0.928 | 23.5 | 0.928 | 0.85 | 0.96 |
| 1,2-DG(38:5) | 21.4 | 1.02 | 21.3 | 1.04 | 0.76 | 0.93 |
| 1,2-DG(38:6) | 18.9 | 1.36 | 18.6 | 1.45 | 0.2 | 0.8 |
| 1,2-DG(40:6) | 19 | 1.32 | 18.8 | 1.25 | 0.76 | 0.88 |
| 1,2-DG(40:7) | 18.1 | 1.22 | 17.9 | 1.32 | 0.62 | 0.88 |
| **Cholesteryl esters (CE)** | | | | | | |
| CE(20:4) | 17.6 | 0.704 | 17.9 | 0.505 | 0.57 | 1.2 |
| CE(22:6) | 17.1 | 0.94 | 17.2 | 0.958 | 0.95 | 1.1 |

| | | | | | | |
|---|---|---|---|---|---|---|
| **Ceramides (Cer)** | | | | | | |
| Cer(40:2) | 25 | 0.734 | 24.9 | 0.752 | 0.96 | 0.92 |
| Cer(d18:0/16:0) | 20.9 | 0.588 | 21.1 | 0.336 | 0.78 | 1.1 |
| Cer(d18:0/18:0) | 18.9 | 0.941 | 19.4 | 1.07 | 0.18 | 1.3 |
| Cer(d18:0/20:0) | 19.2 | 0.794 | 19.5 | 1.02 | 0.61 | 1.2 |
| Cer(d18:0/22:0) | 22.4 | 0.711 | 22.4 | 0.885 | 0.89 | 0.99 |
| Cer(d18:0/22:1) | 22.7 | 1.1 | 22.7 | 1.07 | 0.87 | 0.99 |
| Cer(d18:0/24:0) | 21.8 | 1.04 | 21.8 | 0.822 | 0.87 | 0.98 |
| Cer(d18:0/24:1) | 23.1 | 1.09 | 22.9 | 0.995 | 1 | 0.91 |
| Cer(d18:0/26:0) | 17.9 | 1.59 | 18.1 | 1.86 | 0.31 | 1.2 |
| Cer(d18:0/26:1) | 16.4 | 0.655 | 16.3 | 0.553 | 0.69 | 0.9 |
| Cer(d18:0/26:2)/(18:2/26:0) | 19.6 | 1.02 | 19.3 | 1.29 | 0.87 | 0.83 |
| Cer(d18:1/16:0) | 23.2 | 0.559 | 23.2 | 0.415 | 0.89 | 0.99 |
| Cer(d18:1/18:0) | 22.2 | 0.587 | 22 | 0.541 | 0.61 | 0.86 |
| Cer(d18:1/18:1) | 18.4 | 1.15 | 18.1 | 0.741 | 0.11 | 0.85 |
| Cer(d18:1/20:0) | 20.2 | 1.16 | 20.2 | 1.02 | 0.87 | 0.97 |
| Cer(d18:1/22:0) | 21.9 | 1.08 | 21.9 | 1.07 | 0.85 | 1.1 |
| Cer(d18:1/23:0) | 19.9 | 0.727 | 19.9 | 0.699 | 0.89 | 0.99 |
| Cer(d18:1/23:2) | 17.6 | 1.38 | 16.8 | 1.25 | 0.047 | 0.57 |
| Cer(d18:1/23:2)/(d18:2/23:1) | 18.5 | 0.348 | 18.3 | 0.947 | 0.86 | 0.83 |
| Cer(d18:1/24:0) | 20.8 | 0.661 | 20.7 | 0.888 | 0.92 | 0.94 |
| Cer(d18:1/24:1)/(d18:2/24:0) | 24.5 | 0.248 | 24.5 | 0.48 | 0.78 | 1 |
| Cer(d18:1/25:0) | 17.9 | 1.4 | 18 | 1.34 | 0.52 | 1.1 |
| Cer(d18:1/26:0) | 17.9 | 1.45 | 17.9 | 1.62 | 0.66 | 1 |
| Cer(d18:1/26:1) | 17.5 | 1.05 | 17.4 | 1.04 | 0.83 | 0.97 |
| Cer(d18:2/16:0) | 17.6 | 1.42 | 17.2 | 1.33 | 0.2 | 0.75 |
| Cer(d18:2/18:0) | 17.5 | 1.26 | 17.4 | 0.954 | 0.47 | 0.92 |
| Cer(d18:2/23:0)/(18:1/23:1) | 20.9 | 1.2 | 20.8 | 1.2 | 0.85 | 0.91 |
| Cer(d18:2/24:1)(18:1/24:2) | 22.7 | 1.68 | 22.4 | 1.51 | 0.92 | 0.84 |
| Cer(d36:2) | 18.6 | 0.665 | 18.7 | 0.447 | 0.91 | 1 |
| Cholesterol | 17.2 | 1.55 | 16.8 | 1.29 | 0.61 | 0.74 |
| **lyso-phosphatidylcholines (LPC)** | | | | | | |
| LPC(16:0) | 22.5 | 1.09 | 22.6 | 1.21 | 0.96 | 1.1 |
| LPC(16:1) | 15.8 | 2.35 | 15.4 | 1.73 | 0.68 | 0.75 |
| LPC(18:1) | 20.1 | 1.07 | 20.1 | 1.2 | 0.64 | 0.96 |
| LPC(18:2) | 18.3 | 1.29 | 18.4 | 1.39 | 0.93 | 1.1 |
| LPC(20:0) | 16.5 | 1.27 | 16.7 | 1.35 | 0.87 | 1.2 |
| LPC(20:1) | 16.6 | 0.942 | 16.7 | 0.786 | 0.87 | 1 |
| LPC(20:4) | 17.5 | 0.77 | 17.4 | 0.859 | 0.72 | 0.93 |
| LPC(22:0) | 16.4 | 0.91 | 16.5 | 0.931 | 0.92 | 1.1 |
| LPC(22:6) | 15.2 | 1.14 | 15.1 | 0.986 | 0.44 | 0.98 |
| LPC(24:0) | 17.3 | 1.17 | 17.4 | 1.11 | 0.8 | 1.1 |
| **lyso-phosphatidylethanolamines (LPE)** | | | | | | |

| | | | | | | |
|---|---|---|---|---|---|---|
| LPE(16:0) | 18.9 | 1.57 | 19.1 | 1.7 | 0.93 | 1.2 |
| LPE(18:0) | 20.1 | 2.1 | 20.8 | 1.52 | 0.37 | 1.6 |
| LPE(18:1) | 20 | 0.959 | 20.3 | 0.602 | 0.8 | 1.2 |
| LPE(18:2) | 15.8 | 1.26 | 15.5 | 1.15 | 0.39 | 0.82 |
| LPE(20:1) | 18.3 | 1 | 18.6 | 0.728 | 0.68 | 1.3 |
| LPE(20:4) | 18.5 | 0.988 | 19 | 1.08 | 0.4 | 1.5 |
| LPE(22:4) | 17.4 | 0.89 | 17.6 | 0.848 | 0.72 | 1.2 |
| LPE(22:5) | 14.9 | 1.07 | 14.9 | 0.904 | 0.52 | 1 |
| LPE(22:6) | 17.5 | 1.21 | 18 | 1.2 | 0.69 | 1.4 |
| **lyso-phosphatidylinositol (LPI)** | | | | | | |
| LPI(18:0) | 16.6 | 1.18 | 16.8 | 1.61 | 0.91 | 1.1 |
| **Phosphatidylcholines (PC)** | | | | | | |
| PC(16:0_16:0) | 25.8 | 0.264 | 26.1 | 0.382 | 0.095 | 1.3 |
| PC(16:0_18:0) | 23.4 | 0.393 | 23.6 | 0.242 | 0.14 | 1.1 |
| PC(16:0_18:1) | 26.9 | 0.363 | 26.9 | 0.23 | 0.95 | 0.99 |
| PC(16:0_18:2)/(16:1_18:1) | 25.8 | 0.211 | 25.5 | 0.293 | 0.34 | 0.8 |
| PC(16:0_18:2) | 24.7 | 0.663 | 25.1 | 0.794 | 0.099 | 1.3 |
| PC(16:0_20:2)/(18:1_18:1)/(18:0_18:2) | 25.4 | 0.375 | 25.3 | 0.285 | 0.45 | 0.92 |
| PC(16:0_20:3)/(18:2_18:1) | 24.4 | 0.565 | 24.2 | 0.488 | 0.68 | 0.91 |
| PC(16:0_20:3) | 22.9 | 0.231 | 23 | 0.23 | 0.67 | 1.1 |
| PC(16:0_20:4) | 25.7 | 0.34 | 25.8 | 0.362 | 0.74 | 1.1 |
| PC(16:0_20:5) | 19.9 | 1.76 | 19.7 | 1.46 | 0.59 | 0.86 |
| PC(16:0_22:4)/(18:2_20:2)/(18:1_20:3) | 22.9 | 0.298 | 22.4 | 0.132 | 0.22 | 0.72 |
| PC(16:0_22:4) | 21.5 | 0.689 | 21.6 | 0.664 | 0.87 | 1.1 |
| PC(16:0_22:5)/(18:0_20:5)/(18:1_20:4) | 24.1 | 0.623 | 24.2 | 0.47 | 0.96 | 1 |
| PC(16:0_22:6) | 24.3 | 0.693 | 24 | 0.718 | 0.3 | 0.82 |
| PC(16:1_14:0)/(30:1) | 19.6 | 0.362 | 19.6 | 0.283 | 0.93 | 1 |
| PC(16:1_18:1) | 23.1 | 1.91 | 22.8 | 1.9 | 0.99 | 0.78 |
| PC(16:1_18:2)/(18:3_16:0) | 23.7 | 1.36 | 23.2 | 1.53 | 0.23 | 0.71 |
| PC(16:1_18:2) | 21.2 | 0.736 | 21.4 | 0.733 | 0.89 | 1.1 |
| PC(16:1_20:3)/(18:2_18:2) | 22 | 0.439 | 22.3 | 0.434 | 0.62 | 1.2 |
| PC(16:1_20:4) | 20.9 | 1.58 | 21.1 | 1.21 | 0.86 | 1.2 |
| PC(18:0_20:4) | 24.8 | 0.385 | 24.8 | 0.384 | 0.92 | 1 |
| PC(18:0_22:4)/(40:4) | 21.1 | 0.517 | 21.1 | 0.671 | 0.98 | 1 |
| PC(18:0_22:4) | 20.3 | 0.433 | 20.5 | 0.245 | 0.66 | 1.1 |
| PC(18:0_22:5) | 20.3 | 1.01 | 20.3 | 1.31 | 0.86 | 1 |
| PC(18:0_22:5)/(20:4_20:1)/(40:5) | 22.5 | 0.392 | 22.1 | 0.321 | 0.61 | 0.77 |
| PC(18:0_22:6) | 21.5 | 1.93 | 21.2 | 2.3 | 0.33 | 0.81 |
| PC(18:1_22:5) | 18.5 | 0.857 | 18.7 | 0.787 | 0.96 | 1.1 |
| PC(18:1_22:5)/(20:4_20:2)/(18:2_22:4) | 20.6 | 0.911 | 20.7 | 0.274 | 0.93 | 1 |

| | | | | | | |
|---|---|---|---|---|---|---|
| PC(18:1_22:6)/(20:3_20:4)/(40:7) | 21.8 | 0.707 | 21.5 | 0.202 | 0.37 | 0.76 |
| PC(18:1_22:6) | 18.9 | 1.25 | 18.5 | 0.698 | 0.44 | 0.72 |
| PC(18:1_24:0)/(16:0_26:1) | 20 | 0.7 | 20 | 0.678 | 0.85 | 1.1 |
| PC(18:2_18:1) | 23.4 | 0.182 | 23.5 | 0.297 | 0.76 | 1.1 |
| PC(18:2_18:2) | 21.2 | 1.17 | 20.8 | 0.805 | 0.34 | 0.8 |
| PC(18:2_20:2) | 17.4 | 1.49 | 18.1 | 1.81 | 0.52 | 1.6 |
| PC(18:2_20:4) | 22.3 | 0.527 | 22.3 | 0.604 | 0.95 | 1 |
| PC(18:2_22:4) | 19 | 0.718 | 19 | 0.793 | 0.63 | 0.97 |
| PC(18:3_16:0) | 20.1 | 0.699 | 20.4 | 0.833 | 0.35 | 1.3 |
| PC(20:3_20:4) | 18.4 | 0.639 | 18.6 | 0.616 | 0.9 | 1.1 |
| PC(20:4_20:1) | 19.4 | 2.26 | 19.7 | 2.14 | 0.78 | 1.2 |
| PC(20:4_20:2) | 19.7 | 0.573 | 20 | 0.526 | 0.24 | 1.3 |
| PC(30:1) | 18.6 | 1.07 | 18.2 | 1.02 | 0.76 | 0.78 |
| PC(32:1) | 24.3 | 0.449 | 24.3 | 0.337 | 0.91 | 1 |
| PC(32:2) | 19.8 | 1.63 | 20.1 | 1.59 | 0.96 | 1.2 |
| PC(36:0) | 22.9 | 1.52 | 22.8 | 1.4 | 0.38 | 0.97 |
| PC(36:1) | 25.4 | 0.55 | 25.3 | 0.582 | 0.96 | 0.96 |
| PC(38:2) | 22.3 | 0.657 | 22.3 | 0.68 | 0.86 | 0.98 |
| PC(40:2) | 20.2 | 0.768 | 20.2 | 0.874 | 0.87 | 1 |
| PC(40:5) | 19.8 | 1.66 | 19.6 | 1.72 | 0.63 | 0.87 |
| PC(40:7) | 21.2 | 0.804 | 21 | 1.08 | 0.3 | 0.83 |
| PC(40:8) | 21 | 0.661 | 21 | 0.575 | 0.92 | 1 |
| **Phosphatidylcholine-ether lipids (PC(O))** | | | | | | |
| PC(O-16:0) | 19.3 | 1.15 | 19.5 | 1.16 | 0.85 | 1.1 |
| PC(O-16:0_16:0) | 23.8 | 0.391 | 24 | 0.538 | 0.36 | 1.2 |
| PC(O-16:0_20:3) | 19.2 | 0.776 | 19.4 | 0.875 | 0.66 | 1.2 |
| PC(O-16:0_20:4) | 23.5 | 0.393 | 23.6 | 0.368 | 0.56 | 1.1 |
| PC(O-16:1_18:1) | 20.5 | 1.17 | 20.3 | 0.917 | 0.87 | 0.85 |
| PC(O-16:1_20:4) | 21.6 | 1.21 | 21.4 | 1 | 0.74 | 0.82 |
| PC-(O-18:0_18:2) | 21.2 | 0.714 | 21.4 | 1.16 | 0.8 | 1.2 |
| PC(O-18:0_18:3) | 21.7 | 0.726 | 21.7 | 0.691 | 0.93 | 0.99 |
| PC(O-18:1_22:6) | 18.3 | 0.974 | 18.3 | 0.853 | 0.92 | 0.97 |
| PC(O-20:0_16:0) | 18.4 | 1.01 | 18.2 | 0.846 | 0.69 | 0.88 |
| PC(O-30:0) | 19.3 | 1.29 | 19.8 | 0.616 | 0.45 | 1.4 |
| PC(O-32:1) | 21.3 | 1.02 | 21.2 | 1.11 | 0.85 | 0.92 |
| PC(O-32:2) | 19.6 | 2.7 | 19 | 2.57 | 0.83 | 0.67 |
| PC(O-34:0) | 21.2 | 0.396 | 21.5 | 0.516 | 0.26 | 1.2 |
| PC(O-34:1) | 24.1 | 0.369 | 24.3 | 0.623 | 0.24 | 1.1 |
| PC(O-34:2) | 22.7 | 0.823 | 22.8 | 0.829 | 0.85 | 1 |
| PC(O-34:2)/(O-16:1_18:1) | 23.6 | 0.262 | 23 | 0.735 | 0.57 | 0.65 |
| PC(O-36:1) | 20.2 | 1.04 | 20.2 | 1.08 | 0.69 | 1.1 |
| PC-(O-36:2) | 21.6 | 1.29 | 21.4 | 1.53 | 0.98 | 0.85 |
| PC(O-38:5) | 23.3 | 0.488 | 23.3 | 0.284 | 0.98 | 0.99 |

| | | | | | | |
|---|---|---|---|---|---|---|
| PC(O-38:6) | 19.6 | 0.749 | 19.7 | 0.766 | 0.99 | 1.1 |
| PC(O-40:7) | 20.4 | 0.222 | 20.9 | 0.129 | 0.0029 | 1.4 |
| **Plasmenylcholines (PC(P))** | | | | | | |
| PC(P-16:0_16:0) | 19.4 | 0.333 | 19.4 | 0.408 | 0.93 | 0.99 |
| PC(P-18:0_18:1) | 23.1 | 0.963 | 23.1 | 0.861 | 0.93 | 0.95 |
| PC(P-18:0_22:6) | 19.3 | 0.954 | 19.2 | 0.942 | 0.96 | 1 |
| **Phosphatidylethanolamines (PE)** | | | | | | |
| PE(16:0_16:1) | 17.8 | 1.31 | 18 | 1.28 | 0.9 | 1.1 |
| PE(16:0_18:0) | 18.7 | 0.724 | 18.9 | 0.741 | 0.68 | 1.1 |
| PE(16:0_18:2)/(16:1_18:1) | 20.2 | 1.77 | 21 | 0.968 | 0.36 | 1.8 |
| PE(16:0_18:2) | 22 | 0.0774 | 22.2 | 0.195 | 0.2 | 1.2 |
| PE(16:0_20:2)/(18:1_18:1)/(18:0_18:2) | 23.7 | 1.41 | 24 | 0.836 | 0.81 | 1.2 |
| PE(16:0_20:3) | 20.3 | 0.0507 | 20.6 | 0.236 | 0.24 | 1.2 |
| PE(16:0_20:3)/(18:1_18:2) | 19.9 | 2.55 | 21.3 | 0.942 | 0.33 | 2.5 |
| PE(16:0_20:4) | 22.5 | 1.67 | 23.2 | 0.881 | 0.26 | 1.7 |
| PE(16:0_22:4)/(18:0_20:4) | 22.6 | 1.82 | 24.4 | 0.221 | 0.43 | 3.5 |
| PE(16:0_22:4) | 20.7 | 0.625 | 21 | 0.761 | 0.33 | 1.2 |
| PE(16:0_22:5) | 20.4 | 1.41 | 20.7 | 1.21 | 0.69 | 1.2 |
| PE(16:0_22:6) | 21.9 | 1.59 | 22.4 | 0.811 | 0.59 | 1.4 |
| PE(16:1_18:1) | 19.6 | 0.368 | 19.3 | 0.298 | 0.5 | 0.83 |
| PE(16:1_20:4) | 18.8 | 1.86 | 19.3 | 1.31 | 0.61 | 1.4 |
| PE(18:0_20:3) | 17.4 | 2.51 | 18.1 | 1.99 | 0.61 | 1.6 |
| PE(18:0_20:4) | 25 | 1.1 | 25.2 | 0.896 | 0.85 | 1.1 |
| PE(18:0_22:4) | 21.4 | 1.78 | 22.1 | 0.588 | 0.3 | 1.7 |
| PE(18:0_22:5) | 20.6 | 1.85 | 21.2 | 0.804 | 0.44 | 1.5 |
| PE(18:0_22:6) | 23.5 | 1.69 | 23.7 | 0.89 | 0.96 | 1.2 |
| PE(18:1_18:2) | 22.3 | 0.0308 | 22.4 | 0.264 | 0.86 | 1 |
| PE(18:1_20:4) | 23.6 | 1.1 | 23.8 | 0.867 | 0.81 | 1.1 |
| PE(18:1_20:4)/(16:0_22:5) | 20.6 | 2.22 | 23.1 | 0.178 | 0.36 | 5.6 |
| PE(18:1_22:4) | 20.2 | 1.77 | 20.2 | 0.514 | 0.92 | 1 |
| PE(18:1_22:5)/(20:4_20:2) | 17.3 | 1.59 | 18.2 | 0.665 | 0.66 | 1.8 |
| PE(18:1_22:6) | 21.3 | 1.76 | 21.8 | 0.975 | 0.69 | 1.4 |
| PE(18:2_18:2) | 13.5 | 1.26 | 13.8 | 1.48 | 0.63 | 1.2 |
| PE(18:2_20:3) | 14.6 | 1.5 | 15 | 2.16 | 0.45 | 1.3 |
| PE(18:2_20:4) | 17.6 | 2.9 | 18.6 | 2.54 | 0.35 | 2.1 |
| PE(20:4_20:0) | 17.9 | 0.577 | 18.1 | 0.693 | 0.51 | 1.1 |
| PE(20:4_20:1) | 20.1 | 1.32 | 20.6 | 0.949 | 0.43 | 1.4 |
| PE(34:1) | 22.7 | 2.07 | 22.9 | 0.842 | 0.93 | 1.2 |
| PE(36:1) | 22.9 | 1.43 | 23.5 | 0.4 | 0.4 | 1.5 |
| PE(38:2) | 21.6 | 1.36 | 22.2 | 0.452 | 0.35 | 1.5 |
| PE(38:3) | 19.9 | 2.39 | 21 | 0.86 | 0.3 | 2 |
| PE(42:1) | 18.2 | 1.06 | 18 | 0.802 | 0.31 | 0.85 |
| **Phosphatidylethanolamines-ether lipids (PE (O))** | | | | | | |

| | | | | | | |
|---|---|---|---|---|---|---|
| PE(O-16:0_18:1) | 20.5 | 0.752 | 20.5 | 0.72 | 0.78 | 1.1 |
| PE(O-16:1_22:4) | 22.6 | 1.75 | 23.2 | 0.781 | 0.33 | 1.6 |
| PE(O-18:0_18:2) | 19.6 | 1.1 | 19.5 | 1.2 | 0.95 | 0.92 |
| PE(O-18:1_18:1) | 18.4 | 0.742 | 18.7 | 0.952 | 0.81 | 1.2 |
| PE(O-36:2) | 21.9 | 1.73 | 25.6 | 0.805 | 0.07 | 13 |
| PE(O-38:5) | 22.9 | 1.69 | 23.9 | 0.546 | 0.26 | 2 |
| **Plasmenylethalomines (PE (P))** | | | | | | |
| PE(P-16:0_16:0) | 18.4 | 1.71 | 18.8 | 1.63 | 0.34 | 1.3 |
| PE(P-18:0_18:1) | 25.1 | 0.485 | 25.1 | 0.474 | 0.99 | 0.99 |
| PE(P-18:0_20:4) | 24.3 | 0.324 | 24.5 | 0.144 | 0.59 | 1.2 |
| PE(P-18:0_22:6) | 21.9 | 1.65 | 22.4 | 0.673 | 0.46 | 1.5 |
| **Phosphatidylglycerols (PG)** | | | | | | |
| PG(34:1) | 22.6 | 0.309 | 22.5 | 0.452 | 0.75 | 0.92 |
| PG(34:2) | 18.6 | 1.29 | 18.7 | 0.784 | 0.85 | 1.1 |
| PG(36:1) | 18.6 | 1.11 | 18.3 | 0.983 | 0.72 | 0.78 |
| PG(36:2) | 21.4 | 1.14 | 21.5 | 1.3 | 0.92 | 1.1 |
| PG(36:3) | 22 | 0.891 | 22.4 | 0.732 | 0.33 | 1.3 |
| PG(36:4) | 20.6 | 1.23 | 21.2 | 0.969 | 0.2 | 1.5 |
| PG(38:4) | 19.1 | 1.67 | 19.6 | 1.3 | 0.41 | 1.4 |
| PG(38:5) | 22.3 | 1.11 | 22.8 | 0.734 | 0.3 | 1.4 |
| PG(38:6) | 21.9 | 1.4 | 22.5 | 0.923 | 0.2 | 1.5 |
| PG(40:5) | 20.9 | 1.22 | 21.5 | 0.822 | 0.24 | 1.4 |
| PG(40:6) | 20.7 | 1.57 | 21.4 | 1.25 | 0.039 | 1.7 |
| PG(40:7) | 20.2 | 1.29 | 20.9 | 0.983 | 0.31 | 1.6 |
| PG(40:8) | 21.8 | 1.39 | 22.4 | 0.836 | 0.24 | 1.5 |
| PG(40:9) | 16.5 | 0.784 | 16.4 | 0.792 | 0.95 | 0.96 |
| **Phosphatidylinositols (PI)** | | | | | | |
| PI(16:0_18:1) | 21.7 | 1.95 | 21.8 | 2.07 | 0.66 | 1.1 |
| PI(16:0_20:4) | 22.9 | 1.45 | 23.2 | 1.7 | 0.66 | 1.2 |
| PI(16:0_22:6) | 20.5 | 1.3 | 20.5 | 1.53 | 0.89 | 1 |
| PI(18:0_16:0) | 18.6 | 1.97 | 19 | 2.13 | 0.88 | 1.3 |
| PI(18:0_18:2) | 22 | 0.595 | 22.3 | 0.769 | 0.24 | 1.3 |
| PI(18:0_18:2)/(18:1_18:1) | 21.2 | 0.531 | 20.9 | 0.87 | 0.4 | 0.77 |
| PI(18:0_20:3) | 21.5 | 1.44 | 21.7 | 1.8 | 0.98 | 1.2 |
| PI(18:0_20:4) | 26.2 | 1.14 | 26.4 | 1.61 | 0.72 | 1.2 |
| PI(18:0_22:4) | 20.8 | 1.41 | 21 | 1.9 | 0.78 | 1.1 |
| PI(18:0_22:5) | 16.6 | 2.72 | 16.7 | 3.02 | 0.68 | 1.1 |
| PI(18:0_22:6) | 21.7 | 1.46 | 21.8 | 1.62 | 0.93 | 1 |
| PI(18:1_18:0) | 21.2 | 1.21 | 21.3 | 1.31 | 0.69 | 1.1 |
| PI(18:1_18:1) | 22 | 1.71 | 22 | 2.01 | 0.31 | 1.1 |
| PI(18:1_20:3) | 15.7 | 3.27 | 16.3 | 3.18 | 0.3 | 1.5 |
| PI(18:1_20:4) | 23.4 | 1.44 | 23.8 | 1.67 | 0.61 | 1.3 |
| PI(20:1_20:4) | 20.1 | 1.18 | 20.3 | 1.7 | 0.8 | 1.1 |

| | | | | | | |
|---|---|---|---|---|---|---|
| PI(34:2) | 20.2 | 1.88 | 20.6 | 2.17 | 0.45 | 1.3 |
| PI(36:3) | 19 | 2.86 | 19.4 | 2.97 | 0.91 | 1.3 |
| **Sphingomyelins (SM)** | | | | | | |
| SM(18:0/16:0) | 21.9 | 0.491 | 22.2 | 0.487 | 0.47 | 1.2 |
| SM(18:0/18:0) | 20.7 | 0.642 | 20.6 | 0.458 | 0.3 | 0.89 |
| SM(18:0/20:0) | 20 | 0.557 | 20.1 | 0.216 | 0.76 | 1.1 |
| SM(18:0/22:0) | 21.5 | 0.71 | 21.5 | 0.274 | 0.87 | 1 |
| SM(18:0/22:2)/(18:1/22:1)/(18:2/22:0) | 21.3 | 0.322 | 21.7 | 0.428 | 0.0087 | 1.3 |
| SM(18:0/24:1)/(18:1/24:0) | 24.4 | 0.7 | 24.8 | 0.163 | 0.33 | 1.3 |
| SM(18:0/24:1) | 22.4 | 0.607 | 22.4 | 0.395 | 0.87 | 0.98 |
| SM(18:1/14:0) | 17.5 | 0.788 | 17.5 | 0.923 | 0.69 | 1 |
| SM(18:1/15:0) | 18.3 | 0.951 | 18.7 | 1.07 | 0.37 | 1.3 |
| SM(18:1/16:0) | 24.8 | 0.576 | 25.2 | 0.628 | 0.33 | 1.3 |
| SM(18:1/17:0) | 17.6 | 0.805 | 17.9 | 0.832 | 0.38 | 1.2 |
| SM(18:1/18:0) | 23.5 | 0.353 | 23.5 | 0.379 | 0.93 | 1 |
| SM(18:1/20:0) | 22.3 | 0.5 | 22.4 | 0.205 | 0.4 | 1.1 |
| SM(18:1/21:0) | 18.9 | 1.02 | 18.9 | 0.623 | 0.86 | 1 |
| SM(18:1/22:0) | 23.8 | 0.569 | 24.1 | 0.339 | 0.11 | 1.2 |
| SM(18:1/22:1)/(18:2/22:0)/(18:0/22:2) | 22.3 | 0.341 | 22.3 | 0.119 | 0.98 | 1 |
| SM(18:1/23:0) | 21.2 | 0.659 | 21.4 | 0.462 | 0.33 | 1.1 |
| SM(18:1/23:1)/(18:2/23:0) | 20 | 0.469 | 20 | 0.292 | 0.8 | 1 |
| SM(18:1/23:1) | 19 | 0.0974 | 19.5 | 0.631 | 0.61 | 1.4 |
| SM(18:1/24:0) | 24.1 | 0.739 | 24.2 | 0.432 | 0.36 | 1.1 |
| SM(18:1/24:1)_(18:2/24:0) | 25.5 | 0.58 | 25.7 | 0.42 | 0.095 | 1.2 |
| SM(18:1/24:2)_(18:2/24:1) | 23.2 | 0.499 | 23.6 | 0.518 | 0.069 | 1.3 |
| SM(18:1/25:0)_(18:0/25:1) | 15.5 | 1.04 | 15.7 | 1.17 | 0.93 | 1.2 |
| SM(18:2/25:0)_(18:1/25:1) | 18 | 0.766 | 17.9 | 0.586 | 0.45 | 0.91 |
| SM(36:2) | 18.7 | 0.834 | 18.8 | 0.88 | 0.72 | 1 |
| SM(40:3) | 17.1 | 1.08 | 16.6 | 0.88 | 0.34 | 0.7 |
| **Triglycerides (TG)** | | | | | | |
| TG(48:1) | 27.3 | 0.836 | 26.6 | 0.766 | 0.039 | 0.62 |
| TG(48:2) | 27.5 | 0.79 | 26.7 | 0.755 | 0.0086 | 0.56 |
| TG(50:0) | 24.8 | 1.28 | 24.3 | 1.14 | 0.4 | 0.69 |
| TG(50:1) | 28.1 | 0.768 | 27.4 | 0.702 | 0.022 | 0.62 |
| TG(50:2) | 28.8 | 0.668 | 28.1 | 0.648 | 0.0087 | 0.62 |
| TG(50:3) | 28.4 | 0.684 | 27.6 | 0.623 | 0.0029 | 0.56 |
| TG(50:4) | 26.8 | 0.748 | 25.8 | 0.694 | 0.0029 | 0.49 |
| TG(52:1) | 26.6 | 0.989 | 25.8 | 1.09 | 0.095 | 0.58 |
| TG(52:2) | 28.8 | 0.713 | 28.1 | 0.735 | 0.0074 | 0.61 |
| TG(52:3) | 29.2 | 0.574 | 28.5 | 0.599 | 0.0044 | 0.62 |
| TG(52:4) | 28.5 | 0.512 | 27.8 | 0.495 | 0.0029 | 0.59 |
| TG(52:5) | 26.9 | 0.617 | 25.9 | 0.555 | 0.0029 | 0.51 |

| | | | | | | |
|---|---|---|---|---|---|---|
| TG(54:2) | 26.8 | 0.854 | 25.9 | 1.07 | 0.016 | 0.54 |
| TG(54:3) | 28.4 | 0.701 | 27.6 | 0.791 | 0.0055 | 0.58 |
| TG(54:4) | 28.4 | 0.515 | 27.7 | 0.637 | 0.0086 | 0.61 |
| TG(54:5) | 27.8 | 0.457 | 27.1 | 0.599 | 0.012 | 0.6 |
| TG(54:6) | 26.6 | 0.517 | 25.7 | 0.621 | 0.0086 | 0.56 |
| TG(56:6) | 25.1 | 0.503 | 24.3 | 0.643 | 0.0029 | 0.54 |
| TG(58:10) | 20.6 | 0.538 | 20 | 0.745 | 0.11 | 0.64 |
| TG(58:6) | 22.7 | 0.51 | 21.8 | 0.731 | 0.0029 | 0.56 |
| TG(58:8) | 23 | 0.482 | 22.3 | 0.68 | 0.0074 | 0.58 |
| TG(60:10) | 19.4 | 0.526 | 18.9 | 0.943 | 0.35 | 0.72 |
| TG(60:11) | 18.7 | 0.671 | 17.8 | 1.34 | 0.18 | 0.54 |
| TG(60:12) | 16.8 | 1.61 | 16.3 | 1.27 | 0.33 | 0.68 |

**Table S4. Lipid species detected in megakaryocytes from the bone marrow of *Lepr-cre; Cxcl14*$^{fl/fl}$ and littermate control mice fed a high fat diet (HFD1) in Figure 5.** Fold change (FC) reflects values from megakaryocytes from *Lepr-cre; Cxcl14* $^{fl/fl}$ mice divided by megakaryocytes from control mice. Lipid species highlighted in red have an FDR < 0.05 and a log2 FC > 0.5 in either direction.

| Lipid species | Control | | *Lepr-cre;Cxcl14*$^{flfl}$ | | | |
|---|---|---|---|---|---|---|
| | Log2-mean | Log2-SD | Log2-mean | Log2-SD | FDR | FC |
| **Free Fatty acids (FFA)** | | | | | | |
| (16:0) palmitic acid | 20.1 | 0.16 | 20.3 | 0.477 | 0.59 | 1.2 |
| (18:0) stearic acid | 20.8 | 0.363 | 19.3 | 0 | 0 | 0.36 |
| (18:1) oleic acid | 18.8 | 0.48 | 19 | 0.2 | 0.49 | 1.2 |
| (18:2) linoleic acid | 17.1 | 0.521 | 17.4 | 0.471 | 0.49 | 1.2 |
| **Diacylglycerols (DG)** | | | | | | |
| 1,2-DG(18:1/18:3)_(18:2/18:2) | 23.4 | 0.285 | 23.7 | 0.268 | 0.31 | 1.2 |
| 1,2-DG(20:4/16:0) | 22.1 | 0.197 | 22.5 | 0.433 | 0.26 | 1.3 |
| 1,2-DG(32:0) | 20.4 | 0.323 | 20.8 | 0.436 | 0.32 | 1.3 |
| 1,2-DG(32:1) | 19.6 | 0.386 | 20.2 | 0.586 | 0.27 | 1.5 |
| 1,2-DG(32:2) | 19.1 | 0.697 | 19.7 | 0.392 | 0.31 | 1.4 |
| 1,2-DG(34:0) | 22.4 | 0.503 | 22.8 | 0.576 | 0.45 | 1.3 |
| 1,2-DG(34:1) | 22.9 | 0.227 | 23.2 | 0.354 | 0.32 | 1.2 |
| 1,2-DG(34:2) | 23.7 | 0.361 | 24.1 | 0.349 | 0.22 | 1.3 |
| 1,2-DG(34:3) | 21 | 0.0464 | 21.6 | 0.491 | 0.18 | 1.5 |
| 1,2-DG(36:0) | 23.9 | 0.663 | 24.3 | 0.649 | 0.52 | 1.3 |
| 1,2-DG(36:1) | 22.1 | 0.303 | 22.2 | 0.349 | 0.71 | 1.1 |
| 1,2-DG(36:2) | 24 | 0.377 | 24.4 | 0.328 | 0.27 | 1.3 |
| 1,2-DG(36:3) | 24.7 | 0.365 | 25.1 | 0.378 | 0.32 | 1.3 |
| 1,2-DG(38:0) | 11.8 | 0.577 | 12.3 | 1.09 | 0.62 | 1.4 |
| 1,2-DG(38:1) | 18.3 | 0.213 | 18.9 | 1.04 | 0.47 | 1.5 |
| 1,2-DG(38:4) | 24.4 | 0.374 | 24.8 | 0.205 | 0.17 | 1.3 |
| 1,2-DG(38:5) | 22.1 | 0.255 | 22.4 | 0.369 | 0.49 | 1.2 |
| 1,2-DG(38:6) | 19.6 | 0.187 | 19.5 | 0.74 | 0.9 | 0.96 |
| 1,2-DG(40:6) | 20.3 | 0.196 | 20.5 | 0.377 | 0.63 | 1.1 |
| 1,2-DG(40:7) | 19 | 0.704 | 18.8 | 1.6 | 0.88 | 0.88 |
| **Cholesteryl esters (CE)** | | | | | | |
| CE(20:4) | 18.5 | 0.535 | 18.6 | 0.261 | 0.8 | 1.1 |
| CE(22:6) | 17.9 | 0.116 | 16.5 | 0.66 | 0.0083 | 0.36 |
| **Ceramides (Cer)** | | | | | | |
| Cer(40:2) | 24.5 | 0.487 | 25 | 0.297 | 0.21 | 1.4 |
| Cer(d18:0/16:0) | 20.3 | 0.253 | 20.6 | 0.284 | 0.32 | 1.2 |
| Cer(d18:0/22:0) | 23.2 | 0.475 | 23.7 | 0.132 | 0.12 | 1.4 |
| Cer(d18:0/22:1) | 22.7 | 0.536 | 23.1 | 0.361 | 0.32 | 1.3 |

| | | | | | | |
|---|---|---|---|---|---|---|
| Cer(d18:0/24:0) | 20.6 | 0.492 | 20.8 | 0.334 | 0.65 | 1.1 |
| Cer(d18:0/24:1) | 22.7 | 0.296 | 22.9 | 0.396 | 0.44 | 1.2 |
| Cer(d18:0/26:0) | 19.3 | 0.352 | 19.7 | 0.478 | 0.38 | 1.3 |
| Cer(d18:0/26:2)/(18:2/26:0) | 19.3 | 0.583 | 19.6 | 0.207 | 0.44 | 1.2 |
| Cer(d18:1/16:0) | 22.9 | 0.585 | 23 | 0.298 | 0.88 | 1 |
| Cer(d18:1/18:0) | 21.9 | 0.326 | 22.5 | 0.328 | 0.12 | 1.5 |
| Cer(d18:1/20:0) | 20.8 | 0.3 | 21 | 0.365 | 0.54 | 1.2 |
| Cer(d18:1/22:0) | 22.6 | 0.443 | 23 | 0.316 | 0.26 | 1.3 |
| Cer(d18:1/23:0) | 19.9 | 0.368 | 20.1 | 0.342 | 0.61 | 1.1 |
| Cer(d18:1/24:0) | 20.3 | 0.386 | 20.6 | 0.291 | 0.32 | 1.2 |
| Cer(d18:1/24:1)/(d18:2/24:0 ) | 24.3 | 0.256 | 24.6 | 0.297 | 0.32 | 1.2 |
| Cer(d18:1/25:0) | 17.7 | 0.24 | 17.1 | 1.07 | 0.49 | 0.66 |
| Cer(d18:1/26:0) | 17.4 | 0.906 | 18.2 | 0.374 | 0.17 | 1.8 |
| Cer(d18:1/26:1) | 17.1 | 1.02 | 17.3 | 0.918 | 0.81 | 1.2 |
| Cer(d18:2/23:0)/(18:1/23:1) | 21.3 | 0.41 | 21.6 | 0.362 | 0.32 | 1.3 |
| Cer(d18:2/24:1)(18:1/24:2) | 21.1 | 0.105 | 21.3 | 0.229 | 0.32 | 1.1 |
| Cer(d36:2) | 19 | 0.495 | 19.1 | 0.373 | 0.72 | 1.1 |
| Cholesterol | 17.7 | 1.77 | 18 | 0.734 | 0.81 | 1.2 |
| **lyso-phosphatidylcholines (LPC)** | | | | | | |
| LPC(16:0) | 21.9 | 0.128 | 21.8 | 0.547 | 0.84 | 0.94 |
| LPC(16:1) | 20.3 | 0.0876 | 20.1 | 0.24 | 0.32 | 0.87 |
| LPC(18:1) | 19.9 | 0.196 | 19.7 | 0.668 | 0.68 | 0.86 |
| LPC(18:2) | 18.7 | 0.161 | 19.4 | 0.252 | 0.00033 | 1.7 |
| LPC(20:4) | 18.2 | 0.285 | 18.8 | 0.22 | 0.034 | 1.5 |
| **lyso-phosphatidylethanolamines (LPE)** | | | | | | |
| LPE(16:0) | 18.8 | 0.431 | 17.2 | 1.22 | 0.14 | 0.33 |
| LPE(18:0) | 19.4 | 1.22 | 18.3 | 0.389 | 0.17 | 0.48 |
| LPE(18:1) | 21.2 | 0.385 | 20 | 1.04 | 0.17 | 0.44 |
| LPE(20:1) | 19.7 | 0.962 | 17.8 | 1.21 | 0.12 | 0.27 |
| LPE(20:4) | 20.6 | 0.301 | 20 | 0.641 | 0.32 | 0.69 |
| LPE(22:4) | 19.5 | 0.386 | 17.2 | 1.85 | 0.14 | 0.2 |
| LPE(22:6) | 18.3 | 1.72 | 17.4 | 1.5 | 0.56 | 0.54 |
| **lyso-phosphatidylinositol (LPI)** | | | | | | |
| LPI(18:0) | 14.1 | 0 | 15.5 | 1.95 | 0.34 | 2.7 |
| **Phosphatidylcholines (PC)** | | | | | | |
| PC(16:0_16:0) | 25.8 | 0.378 | 25.6 | 0.204 | 0.45 | 0.87 |
| PC(16:0_18:0) | 23.7 | 0.402 | 24.1 | 0.35 | 0.32 | 1.3 |
| PC(16:0_18:1) | 27.2 | 0.0433 | 27.1 | 0.254 | 0.79 | 0.96 |
| PC(16:0_18:2)/(16:1_18:1) | 26.2 | 0.11 | 25.9 | 0.521 | 0.49 | 0.82 |
| PC(16:0_20:2)/(18:1_18:1)/( 18:0_18:2) | 25.9 | 0.132 | 25.9 | 0.171 | 0.96 | 1 |
| PC(16:0_20:3)/(18:2_18:1) | 24.6 | 0.13 | 24.2 | 0.51 | 0.45 | 0.81 |
| PC(16:0_20:4) | 26.4 | 0.18 | 25.9 | 0.4 | 0.14 | 0.69 |

| | | | | | | |
|---|---|---|---|---|---|---|
| PC(16:0_20:5) | 21.8 | 0.535 | 21.7 | 0.366 | 0.7 | 0.89 |
| PC(16:0_22:4)/(18:2_20:2)/(18:1_20:3) | 23.2 | 0.143 | 23.1 | 0.288 | 0.81 | 0.96 |
| PC(16:0_22:5)/(18:0_20:5)/(18:1_20:4) | 25.1 | 0.194 | 24.9 | 0.504 | 0.65 | 0.88 |
| PC(16:0_22:6) | 24.6 | 0.0479 | 24.1 | 0.305 | 0.082 | 0.72 |
| PC(16:1_18:2)/(18:3_16:0) | 21.9 | 0.127 | 18.3 | 4.52 | 0.32 | 0.081 |
| PC(16:1_20:4) | 21.6 | 0.594 | 21.1 | 0.454 | 0.32 | 0.72 |
| PC(18:0_20:4) | 25.7 | 0.233 | 25.7 | 0.181 | 1 | 1 |
| PC(18:0_22:4)/(40:4) | 22 | 0.162 | 22.3 | 0.262 | 0.21 | 1.2 |
| PC(18:0_22:5)/(20:4_20:1)/(40:5) | 22.7 | 0.261 | 22.6 | 0.148 | 0.68 | 0.95 |
| PC(18:0_22:6) | 23 | 0.19 | 22.8 | 0.379 | 0.68 | 0.91 |
| PC(18:1_22:5)/(20:4_20:2)/(18:2_22:4) | 21.7 | 0.146 | 21.4 | 0.54 | 0.43 | 0.79 |
| PC(18:1_22:6)/(20:3_20:4)/(40:7) | 21.9 | 0.114 | 21.3 | 0.259 | 0.0083 | 0.66 |
| PC(18:2_18:2) | 22.7 | 0.133 | 22.4 | 0.399 | 0.32 | 0.8 |
| PC(18:2_20:4) | 23 | 0.183 | 22.5 | 0.342 | 0.12 | 0.71 |
| PC(32:1) | 23.8 | 0.192 | 23.2 | 0.285 | 0.017 | 0.64 |
| PC(32:2) | 20.2 | 0.54 | 19.7 | 0.317 | 0.21 | 0.7 |
| PC(36:0) | 23.7 | 0.425 | 24.2 | 0.163 | 0.11 | 1.4 |
| PC(36:1) | 25.6 | 0.192 | 25.9 | 0.238 | 0.17 | 1.2 |
| PC(38:2) | 22.7 | 0.232 | 23 | 0.252 | 0.17 | 1.3 |
| PC(40:2) | 20.9 | 0.599 | 21.4 | 0.367 | 0.31 | 1.4 |
| PC(40:8) | 21.3 | 0.512 | 20.9 | 0.315 | 0.22 | 0.72 |
| **Phosphatidylcholine-ether lipids (PC(O))** | | | | | | |
| PC(O-16:0) | 18.4 | 0.0171 | 17.7 | 0.53 | 0.14 | 0.63 |
| PC(O-16:0_16:0) | 23.5 | 0.321 | 23.4 | 0.543 | 0.8 | 0.92 |
| PC(O-16:0_20:4) | 23.8 | 0.178 | 23.6 | 0.389 | 0.62 | 0.9 |
| PC(O-16:1_20:4) | 23.1 | 0.509 | 23.3 | 0.369 | 0.75 | 1.1 |
| PC(O-18:0_18:3) | 22.1 | 0.292 | 22.1 | 0.338 | 0.79 | 0.94 |
| PC(O-30:0) | 18.8 | 0.484 | 18.5 | 1 | 0.71 | 0.81 |
| PC(O-32:1) | 21.7 | 0.266 | 21.6 | 0.347 | 0.65 | 0.91 |
| PC(O-32:2) | 21.1 | 0.572 | 21.1 | 0.449 | 0.89 | 1 |
| PC(O-34:0) | 21.3 | 0.604 | 21.7 | 0.507 | 0.38 | 1.4 |
| PC(O-34:1) | 24.5 | 0.171 | 24.5 | 0.224 | 0.95 | 1 |
| PC(O-34:2)/(O-16:1_18:1) | 23.4 | 0.27 | 23.4 | 0.389 | 0.95 | 0.99 |
| PC(O-36:1) | 20.8 | 0.523 | 21.1 | 0.165 | 0.32 | 1.2 |
| PC-(O-36:2) | 23.2 | 0.542 | 23.7 | 0.108 | 0.14 | 1.4 |
| PC(O-38:5) | 23.4 | 0.102 | 23.2 | 0.229 | 0.28 | 0.86 |
| **Plasmenylcholines (PC(P))** | | | | | | |
| PC(P-16:0_16:0) | 19.2 | 0.192 | 19.3 | 0.0857 | 0.56 | 1.1 |
| PC(P-18:0_22:6) | 20.7 | 0.0991 | 20.4 | 0.426 | 0.35 | 0.81 |
| **Phosphatidylethanolamines (PE)** | | | | | | |

| | | | | | | |
|---|---|---|---|---|---|---|
| PE(16:0_18:2)/(16:1_18:1) | 20.9 | 1.27 | 19.4 | 0.532 | 0.12 | 0.36 |
| PE(16:0_20:2)/(18:1_18:1)/(18:0_18:2) | 23.2 | 1.23 | 22.3 | 0.897 | 0.35 | 0.54 |
| PE(16:0_20:3)/(18:1_18:2) | 21.2 | 1.37 | 19.8 | 0.669 | 0.15 | 0.37 |
| PE(16:0_20:4) | 22.4 | 1.34 | 20.7 | 0.959 | 0.14 | 0.31 |
| PE(16:0_22:4)/(18:0_20:4) | 24.4 | 1.28 | 23.5 | 0.759 | 0.32 | 0.52 |
| PE(16:0_22:6) | 21.4 | 1.43 | 19.8 | 0.817 | 0.14 | 0.32 |
| PE(16:1_20:4) | 16.6 | 1.83 | 15.6 | 0 | 0.32 | 0.48 |
| PE(18:0_20:3) | 16.5 | 1.81 | 15.1 | 1.44 | 0.38 | 0.4 |
| PE(18:0_22:4) | 21.4 | 1.38 | 20.6 | 0.834 | 0.45 | 0.59 |
| PE(18:0_22:5) | 19.3 | 1.39 | 18.1 | 0.963 | 0.29 | 0.43 |
| PE(18:0_22:6) | 22.8 | 1.48 | 21.9 | 1 | 0.39 | 0.51 |
| PE(18:1_20:4)/(16:0_22:5) | 22.7 | 1.33 | 21.4 | 0.613 | 0.17 | 0.4 |
| PE(18:1_22:4) | 19 | 1.3 | 18.4 | 1.14 | 0.63 | 0.68 |
| PE(18:1_22:5)/(20:4_20:2) | 19.1 | 1.26 | 17.7 | 0.63 | 0.15 | 0.4 |
| PE(18:1_22:6) | 20.6 | 1.51 | 18.8 | 1.08 | 0.15 | 0.28 |
| PE(18:2_18:2) | 18.3 | 1.47 | 16.6 | 0.884 | 0.15 | 0.31 |
| PE(18:2_20:3) | 19.9 | 1.34 | 18.5 | 0.589 | 0.14 | 0.37 |
| PE(18:2_20:4) | 19.3 | 1.42 | 17.6 | 0.888 | 0.14 | 0.31 |
| PE(20:4_20:1) | 21.1 | 1.57 | 20.1 | 0.738 | 0.34 | 0.51 |
| PE(34:1) | 21.6 | 1.13 | 20.6 | 0.96 | 0.32 | 0.5 |
| PE(36:1) | 22.5 | 0.991 | 22 | 0.909 | 0.56 | 0.69 |
| PE(38:2) | 21.4 | 1.15 | 20.8 | 0.934 | 0.54 | 0.66 |
| PE(38:3) | 18.9 | 1.56 | 17.7 | 1 | 0.32 | 0.44 |
| **Phosphatidylethanolamines-ether lipids (PE (O))** | | | | | | |
| PE(O-16:1_22:4) | 22.5 | 1.27 | 21.4 | 0.963 | 0.32 | 0.47 |
| PE(O-18:0_18:2) | 21 | 0.28 | 20.9 | 0.385 | 0.82 | 0.95 |
| PE(O-36:2) | 24.2 | 0.979 | 23.7 | 0.988 | 0.58 | 0.7 |
| PE(O-38:5) | 23.5 | 1.43 | 22.6 | 0.896 | 0.38 | 0.53 |
| **Plasmenylethalomines (PE (P))** | | | | | | |
| PE(P-18:0_22:6) | 21.8 | 1.39 | 20.6 | 1.12 | 0.32 | 0.45 |
| **Phosphatidylglycerols (PG)** | | | | | | |
| PG(34:1) | 22.2 | 0.0782 | 21.6 | 0.49 | 0.21 | 0.71 |
| PG(34:2) | 19.3 | 0.428 | 19 | 0.389 | 0.6 | 0.87 |
| PG(36:1) | 19.5 | 0.321 | 19.1 | 0.641 | 0.44 | 0.75 |
| PG(36:2) | 20.5 | 0.193 | 20 | 0.492 | 0.26 | 0.72 |
| PG(36:3) | 21.6 | 0.256 | 21 | 0.439 | 0.12 | 0.64 |
| PG(36:4) | 21.3 | 0.429 | 20.7 | 0.277 | 0.12 | 0.67 |
| PG(38:5) | 21.5 | 0.285 | 20.7 | 0.596 | 0.13 | 0.56 |
| PG(38:6) | 22.2 | 0.402 | 21.5 | 0.335 | 0.086 | 0.62 |
| PG(40:5) | 19.7 | 0.36 | 18.9 | 0.527 | 0.14 | 0.59 |
| PG(40:6) | 19.9 | 0.425 | 19.2 | 0.669 | 0.26 | 0.62 |
| PG(40:7) | 20.3 | 0.44 | 19.7 | 0.607 | 0.31 | 0.67 |
| PG(40:8) | 21.5 | 0.606 | 20.8 | 0.548 | 0.21 | 0.61 |

| | | | | | | |
|---|---|---|---|---|---|---|
| **Phosphatidylinositols (PI)** | | | | | | |
| PI(16:0_20:4) | 20.7 | 0.556 | 20 | 0.311 | 0.12 | 0.62 |
| PI(16:0_22:6) | 18.4 | 0.596 | 17.7 | 0.393 | 0.16 | 0.62 |
| PI(18:0_18:2)/(18:1_18:1) | 21.1 | 0.172 | 20.4 | 0.603 | 0.17 | 0.61 |
| PI(18:0_20:3) | 19.3 | 0.441 | 18.4 | 0.525 | 0.12 | 0.55 |
| PI(18:0_20:4) | 25 | 0.325 | 24.2 | 0.452 | 0.077 | 0.57 |
| PI(18:0_22:4) | 19.1 | 0.391 | 18.7 | 0.768 | 0.53 | 0.76 |
| PI(18:0_22:5) | 17 | 0.313 | 16.2 | 0.597 | 0.16 | 0.59 |
| PI(18:0_22:6) | 19.8 | 0.399 | 19 | 0.487 | 0.12 | 0.58 |
| PI(18:1_20:4) | 21 | 0.593 | 20.3 | 0.311 | 0.12 | 0.6 |
| PI(20:1_20:4) | 18 | 0.364 | 17.1 | 0.359 | 0.014 | 0.53 |
| PI(36:3) | 18.6 | 0.538 | 17.8 | 0.552 | 0.17 | 0.59 |
| **Sphingomyelins (SM)** | | | | | | |
| SM(18:0/16:0) | 22.2 | 0.239 | 21.5 | 0.796 | 0.33 | 0.66 |
| SM(18:0/18:0) | 20.1 | 0.162 | 20.2 | 0.317 | 0.79 | 1.1 |
| SM(18:0/20:0) | 19.7 | 0.359 | 20.1 | 0.248 | 0.25 | 1.3 |
| SM(18:0/22:0) | 21.1 | 0.536 | 21.5 | 0.183 | 0.28 | 1.3 |
| SM(18:0/22:2)/(18:1/22:1)/(18:2/22:0) | 21.6 | 0.111 | 22 | 0.272 | 0.15 | 1.3 |
| SM(18:0/24:1)/(18:1/24:0) | 24.1 | 0.463 | 24.5 | 0.218 | 0.17 | 1.4 |
| SM(18:1/16:0) | 24.7 | 0.149 | 24.2 | 0.625 | 0.32 | 0.7 |
| SM(18:1/18:0) | 23.2 | 0.0566 | 23.3 | 0.389 | 0.82 | 1 |
| SM(18:1/20:0) | 22.3 | 0.161 | 22.6 | 0.249 | 0.17 | 1.2 |
| SM(18:1/21:0) | 17.6 | 0.153 | 18 | 0.312 | 0.18 | 1.3 |
| SM(18:1/22:0) | 23.9 | 0.266 | 24.4 | 0.219 | 0.077 | 1.4 |
| SM(18:1/23:0) | 20.5 | 0.362 | 20.9 | 0.292 | 0.17 | 1.4 |
| SM(18:1/23:1) | 18.5 | 0.268 | 19 | 0.291 | 0.093 | 1.4 |
| SM(18:1/24:1)_(18:2/24:0) | 25 | 0.386 | 25.5 | 0.182 | 0.13 | 1.4 |
| SM(18:1/24:2)_(18:2/24:1) | 23.1 | 0.282 | 23.5 | 0.334 | 0.19 | 1.3 |
| SM(40:3) | 17.9 | 0.252 | 17.9 | 0.259 | 0.81 | 0.96 |
| **Triglycerides (TG)** | | | | | | |
| TG(48:1) | 26.1 | 0.224 | 26.8 | 0.268 | 0.002 | 1.7 |
| TG(48:2) | 26.5 | 0.229 | 27.3 | 0.402 | 0.022 | 1.8 |
| TG(50:0) | 23.6 | 0.18 | 24.3 | 0.408 | 0.067 | 1.6 |
| TG(50:1) | 27.6 | 0.342 | 28.3 | 0.265 | 0.034 | 1.6 |
| TG(50:2) | 28.6 | 0.227 | 29.2 | 0.249 | 0.017 | 1.5 |
| TG(50:3) | 28 | 0.25 | 28.7 | 0.393 | 0.077 | 1.6 |
| TG(50:4) | 26.5 | 0.115 | 27.3 | 0.403 | 0.022 | 1.7 |
| TG(52:1) | 26.4 | 0.358 | 26.9 | 0.486 | 0.21 | 1.5 |
| TG(52:2) | 29 | 0.33 | 29.5 | 0.258 | 0.067 | 1.5 |
| TG(52:3) | 29.6 | 0.195 | 30.2 | 0.281 | 0.06 | 1.4 |
| TG(52:4) | 29.2 | 0.281 | 29.7 | 0.335 | 0.13 | 1.4 |
| TG(52:5) | 27.1 | 0.151 | 27.9 | 0.394 | 0.034 | 1.7 |
| TG(54:2) | 27.2 | 0.332 | 27.7 | 0.432 | 0.24 | 1.4 |

| | | | | | | |
|---|---|---|---|---|---|---|
| TG(54:3) | 29.1 | 0.324 | 29.5 | 0.272 | 0.14 | 1.4 |
| TG(54:4) | 29.5 | 0.176 | 30 | 0.28 | 0.1 | 1.4 |
| TG(54:5) | 29.2 | 0.323 | 29.6 | 0.324 | 0.19 | 1.3 |
| TG(54:6) | 27.8 | 0.188 | 28.4 | 0.345 | 0.077 | 1.5 |
| TG(56:6) | 26 | 0.209 | 26.5 | 0.428 | 0.17 | 1.4 |
| TG(58:10) | 21.6 | 0.0808 | 22.1 | 0.518 | 0.3 | 1.4 |
| TG(58:6) | 23.7 | 0.434 | 24 | 0.416 | 0.48 | 1.2 |
| TG(58:8) | 24.1 | 0.227 | 24.5 | 0.497 | 0.32 | 1.3 |
| TG(60:10) | 20.7 | 0.0927 | 20.8 | 0.605 | 0.89 | 1 |
| TG(60:11) | 19.8 | 0.297 | 20.2 | 0.383 | 0.28 | 1.3 |

**Table S5. Lipid species detected in megakaryocytes from the bone marrow of *Lepr-cre; Cxcl14*$^{fl/fl}$ and littermate control mice fed a high fat diet (HFD2).** Fold change (FC) reflects values from megakaryocytes from *Lepr-cre; Cxcl14* $^{fl/fl}$ mice divided by megakaryocytes from control mice. Lipid species highlighted in red have an FDR < 0.05 and a log2 FC > 0.5 in either direction.

| Lipid species | Control | | *Lepr-cre;Cxcl14*$^{fl/fl}$ | | | |
|---|---|---|---|---|---|---|
| | Log2-mean | Log2-SD | Log2-mean | Log2-SD | FDR | FC |
| **Free Fatty acids (FFA)** | | | | | | |
| (16:0) palmitic acid | 20.9 | 0.569 | 20.9 | 0.265 | 0.98 | 1 |
| (18:0) stearic acid | 20.8 | 0.985 | 21 | 0.374 | 0.93 | 1.2 |
| (18:1) oleic acid | 19.3 | 0.194 | 19.3 | 0.364 | 0.95 | 0.96 |
| (18:2) linoleic acid | 16.8 | 0.214 | 16.3 | 0.362 | 0.21 | 0.73 |
| **Diacylglycerols (DG)** | | | | | | |
| 1,2-DG(18:1/18:3)_(18:2/18:2) | 22.3 | 0.343 | 22 | 0.41 | 0.72 | 0.84 |
| 1,2-DG(20:4/16:0) | 22.1 | 0.108 | 22.3 | 0.278 | 0.71 | 1.1 |
| 1,2-DG(32:0) | 21.1 | 0.599 | 21.1 | 0.361 | 0.98 | 0.97 |
| 1,2-DG(32:1) | 20.7 | 0.151 | 20.4 | 0.272 | 0.36 | 0.84 |
| 1,2-DG(32:2) | 19.3 | 0.773 | 19.1 | 0.769 | 0.95 | 0.9 |
| 1,2-DG(34:0) | 23.3 | 0.492 | 23.1 | 0.475 | 0.85 | 0.86 |
| 1,2-DG(34:1) | 23.5 | 0.225 | 23.4 | 0.162 | 0.65 | 0.9 |
| 1,2-DG(34:2) | 23.8 | 0.256 | 23.6 | 0.316 | 0.83 | 0.9 |
| 1,2-DG(34:3) | 21.1 | 0.476 | 21 | 0.552 | 0.95 | 0.91 |
| 1,2-DG(36:0) | 25 | 0.596 | 24.6 | 0.617 | 0.71 | 0.75 |
| 1,2-DG(36:1) | 22.7 | 0.391 | 22.5 | 0.147 | 0.68 | 0.87 |
| 1,2-DG(36:2) | 24.7 | 0.229 | 24.4 | 0.237 | 0.51 | 0.86 |
| 1,2-DG(36:3) | 24.5 | 0.297 | 24.3 | 0.265 | 0.79 | 0.9 |
| 1,2-DG(38:0) | 12.3 | 1.54 | 13.4 | 2.57 | 0.83 | 2.1 |
| 1,2-DG(38:1) | 19.1 | 1.53 | 19.1 | 1.2 | 0.99 | 1 |
| 1,2-DG(38:4) | 24.6 | 0.0789 | 24.6 | 0.236 | 0.95 | 0.98 |
| 1,2-DG(38:5) | 22.3 | 0.198 | 22.4 | 0.294 | 0.95 | 1 |
| 1,2-DG(38:6) | 20.1 | 0.165 | 20 | 0.321 | 0.95 | 0.96 |
| 1,2-DG(40:6) | 20.6 | 0.276 | 20.5 | 0.108 | 0.93 | 0.96 |
| 1,2-DG(40:7) | 19.7 | 0.279 | 19.6 | 0.35 | 0.91 | 0.92 |
| **Cholesteryl esters (CE)** | | | | | | |
| CE(20:4) | 18.5 | 0.32 | 18.2 | 0.28 | 0.41 | 0.81 |
| CE(22:6) | 17.6 | 1.14 | 17.5 | 0.907 | 0.98 | 0.93 |
| **Ceramides (Cer)** | | | | | | |
| Cer(40:2) | 25.3 | 0.46 | 25.3 | 0.521 | 0.99 | 1 |
| Cer(d18:0/16:0) | 21.1 | 0.137 | 21 | 0.201 | 0.83 | 0.94 |
| Cer(d18:0/22:0) | 24 | 0.462 | 24 | 0.373 | 0.99 | 0.99 |
| Cer(d18:0/22:1) | 23.8 | 0.26 | 23.7 | 0.53 | 0.95 | 0.92 |

| | | | | | | |
|---|---|---|---|---|---|---|
| Cer(d18:0/24:0) | 21.3 | 0.332 | 21.2 | 0.281 | 0.93 | 0.93 |
| Cer(d18:0/24:1) | 23.2 | 0.458 | 23.4 | 0.541 | 0.95 | 1.1 |
| Cer(d18:0/26:0) | 20.1 | 0.262 | 20.3 | 0.319 | 0.83 | 1.1 |
| Cer(d18:0/26:2)/(18:2/26:0) | 20.1 | 0.269 | 20.3 | 0.378 | 0.83 | 1.1 |
| Cer(d18:1/16:0) | 23.4 | 0.104 | 23.4 | 0.305 | 0.99 | 1 |
| Cer(d18:1/18:0) | 22.8 | 0.416 | 22.9 | 0.206 | 0.85 | 1.1 |
| Cer(d18:1/20:0) | 21.4 | 0.507 | 21.5 | 0.266 | 0.95 | 1 |
| Cer(d18:1/22:0) | 23.4 | 0.493 | 23.6 | 0.135 | 0.84 | 1.1 |
| Cer(d18:1/23:0) | 20.6 | 0.425 | 20.6 | 0.395 | 0.96 | 1 |
| Cer(d18:1/24:0) | 21.1 | 0.336 | 21.1 | 0.0644 | 0.98 | 1 |
| Cer(d18:1/24:1)/(d18:2/24:0) | 24.9 | 0.207 | 25 | 0.0436 | 0.33 | 1.1 |
| Cer(d18:1/25:0) | 18.1 | 0.198 | 18.2 | 0.258 | 0.82 | 1.1 |
| Cer(d18:1/26:0) | 18.8 | 0.371 | 18.8 | 0.349 | 0.95 | 0.96 |
| Cer(d18:1/26:1) | 18 | 0.329 | 18.1 | 0.315 | 0.95 | 1 |
| Cer(d18:2/23:0)/(18:1/23:1) | 21.9 | 0.389 | 22 | 0.483 | 0.98 | 1 |
| Cer(d18:2/24:1)(18:1/24:2) | 21 | 0.143 | 21 | 0.218 | 0.99 | 1 |
| Cer(d36:2) | 19.2 | 0.264 | 19.3 | 0.439 | 0.95 | 1 |
| Cholesterol | 18.7 | 0.344 | 18.2 | 0.606 | 0.51 | 0.72 |
| **lyso-phosphatidylcholines (LPC)** | | | | | | |
| LPC(16:0) | 21.2 | 0.964 | 21.6 | 0.61 | 0.83 | 1.3 |
| LPC(16:1) | 20.2 | 0.828 | 20 | 0.872 | 0.95 | 0.89 |
| LPC(18:1) | 19.4 | 0.953 | 19.9 | 0.642 | 0.8 | 1.3 |
| LPC(18:2) | 17.4 | 1.43 | 18.1 | 0.645 | 0.73 | 1.6 |
| LPC(20:4) | 16.8 | 1.42 | 16 | 0 | 0.65 | 0.61 |
| **lyso-phosphatidylethanolamines (LPE)** | | | | | | |
| LPE(16:0) | 17 | 1.21 | 17.4 | 1.47 | 0.93 | 1.4 |
| LPE(18:0) | 17.9 | 0.668 | 18.9 | 0.661 | 0.21 | 2 |
| LPE(18:1) | 19.6 | 1.1 | 20.9 | 0.948 | 0.29 | 2.4 |
| LPE(20:1) | 17.2 | 1.14 | 19.9 | 1.15 | 0.045 | 6.5 |
| LPE(20:4) | 17.8 | 1.88 | 19.4 | 1.85 | 0.55 | 3.1 |
| LPE(22:4) | 16.7 | 1.69 | 17.4 | 2.2 | 0.91 | 1.7 |
| LPE(22:6) | 17 | 1.32 | 18.1 | 1.66 | 0.68 | 2.1 |
| **lyso-phosphatidylinositol (LPI)** | | | | | | |
| LPI(18:0) | 15.1 | 2.09 | 14.1 | 0 | 0.65 | 0.48 |
| **Phosphatidylcholines (PC)** | | | | | | |
| PC(16:0_16:0) | 25.6 | 0.157 | 25.8 | 0.272 | 0.75 | 1.1 |
| PC(16:0_18:0) | 24.3 | 0.25 | 24.3 | 0.266 | 0.95 | 0.95 |
| PC(16:0_18:1) | 27.1 | 0.306 | 27.2 | 0.258 | 0.86 | 1.1 |
| PC(16:0_18:2)/(16:1_18:1) | 25.2 | 0.1 | 25.4 | 0.217 | 0.29 | 1.2 |
| PC(16:0_20:2)/(18:1_18:1)/(18:0_18:2) | 25.7 | 0.285 | 25.7 | 0.244 | 0.99 | 1 |
| PC(16:0_20:3)/(18:2_18:1) | 23.5 | 1.04 | 23.7 | 0.829 | 0.95 | 1.1 |
| PC(16:0_20:4) | 25.9 | 0.124 | 26.1 | 0.274 | 0.51 | 1.2 |
| PC(16:0_20:5) | 21.4 | 0.388 | 21.5 | 0.424 | 0.95 | 1.1 |

| | | | | | | |
|---|---|---|---|---|---|---|
| PC(16:0_22:4)/(18:2_20:2)/(18:1_20:3) | 22.7 | 0.352 | 22.6 | 0.205 | 0.93 | 0.94 |
| PC(16:0_22:5)/(18:0_20:5)/(18:1_20:4) | 24.7 | 0.12 | 24.8 | 0.28 | 0.95 | 1 |
| PC(16:0_22:6) | 24.4 | 0.183 | 24.7 | 0.164 | 0.15 | 1.2 |
| PC(16:1_18:2)/(18:3_16:0) | 20.4 | 0.997 | 20.9 | 0.876 | 0.83 | 1.4 |
| PC(16:1_20:4) | 20.9 | 0.571 | 20.4 | 0.865 | 0.72 | 0.71 |
| PC(18:0_20:4) | 25.4 | 0.279 | 25.3 | 0.246 | 0.95 | 0.97 |
| PC(18:0_22:4)/(40:4) | 22 | 0.0511 | 21.7 | 0.3 | 0.33 | 0.83 |
| PC(18:0_22:5)(20:4_20:1)/(40:5) | 22.3 | 0.272 | 22.3 | 0.24 | 0.98 | 0.99 |
| PC(18:0_22:6) | 23 | 0.389 | 23 | 0.203 | 0.97 | 1 |
| PC(18:1_22:5)/(20:4_20:2)/(18:2_22:4) | 20.9 | 0.235 | 20.9 | 0.204 | 0.95 | 1 |
| PC(18:1_22:6)/(20:3_20:4)/(40:7) | 21.3 | 0.204 | 21.6 | 0.281 | 0.25 | 1.3 |
| PC(18:2_18:2) | 21 | 0.355 | 21.4 | 0.289 | 0.27 | 1.3 |
| PC(18:2_20:4) | 21.8 | 0.0998 | 22 | 0.248 | 0.51 | 1.1 |
| PC(32:1) | 23.4 | 0.155 | 23.8 | 0.227 | 0.069 | 1.4 |
| PC(32:2) | 19.5 | 0.467 | 20.1 | 0.375 | 0.28 | 1.4 |
| PC(36:0) | 24.3 | 0.131 | 23.8 | 0.546 | 0.33 | 0.72 |
| PC(36:1) | 26 | 0.0517 | 26 | 0.249 | 0.98 | 0.99 |
| PC(38:2) | 23 | 0.048 | 22.9 | 0.354 | 0.95 | 0.96 |
| PC(40:2) | 21.5 | 0.206 | 21.2 | 0.366 | 0.65 | 0.85 |
| PC(40:8) | 20.7 | 0.135 | 20.9 | 0.329 | 0.51 | 1.2 |
| **Phosphatidylcholine-ether lipids (PC(O))** | | | | | | |
| PC(O-16:0) | 17.9 | 0.79 | 18.1 | 0.702 | 0.95 | 1.1 |
| PC(O-16:0_16:0) | 23.8 | 0.57 | 23.9 | 0.237 | 0.95 | 1.1 |
| PC(O-16:0_20:4) | 23.6 | 0.14 | 23.5 | 0.326 | 0.95 | 0.96 |
| PC(O-16:1_20:4) | 23.2 | 0.222 | 23 | 0.533 | 0.85 | 0.88 |
| PC(O-18:0_18:3) | 21.7 | 0.246 | 21.8 | 0.169 | 0.67 | 1.1 |
| PC(O-30:0) | 18.8 | 0.359 | 19.3 | 0.116 | 0.071 | 1.4 |
| PC(O-32:1) | 21.9 | 0.061 | 22 | 0.331 | 0.65 | 1.1 |
| PC(O-32:2) | 21.4 | 0.28 | 21.4 | 0.515 | 0.98 | 1 |
| PC(O-34:0) | 21.8 | 0.501 | 21.7 | 0.297 | 0.93 | 0.91 |
| PC(O-34:1) | 24.9 | 0.181 | 24.9 | 0.248 | 0.98 | 1 |
| PC(O-34:2)/(O-16:1_18:1) | 22.7 | 0.287 | 22.8 | 0.176 | 0.8 | 1.1 |
| PC(O-36:1) | 21.5 | 0.23 | 21.4 | 0.268 | 0.86 | 0.93 |
| PC-(O-36:2) | 23.6 | 0.264 | 23.5 | 0.434 | 0.83 | 0.88 |
| PC(O-38:5) | 23.3 | 0.189 | 23.3 | 0.413 | 0.99 | 1 |
| **Plasmenylcholines (PC(P))** | | | | | | |
| PC(P-16:0_16:0) | 19.4 | 0.125 | 19.4 | 0.371 | 0.99 | 0.99 |
| PC(P-18:0_22:6) | 20.8 | 0.212 | 21 | 0.299 | 0.75 | 1.1 |
| **Phosphatidylethanolamines (PE)** | | | | | | |
| PE(16:0_18:2)/(16:1_18:1) | 17.9 | 0.226 | 19.6 | 0.917 | 0.045 | 3.2 |

| | | | | | | |
|---|---|---|---|---|---|---|
| PE(16:0_20:2)/(18:1_18:1)/(18:0_18:2) | 21.2 | 0.612 | 22.4 | 0.996 | 0.25 | 2.3 |
| PE(16:0_20:3)/(18:1_18:2) | 18.2 | 0.384 | 19.9 | 1.01 | 0.069 | 3.2 |
| PE(16:0_20:4) | 19.9 | 0.429 | 21.5 | 0.928 | 0.069 | 2.9 |
| PE(16:0_22:4)/(18:0_20:4) | 22.4 | 0.621 | 23.6 | 0.883 | 0.21 | 2.2 |
| PE(16:0_22:6) | 19.4 | 0.324 | 21 | 1.04 | 0.069 | 3.2 |
| PE(16:1_20:4) | 15.6 | 0 | 16 | 0.907 | 0.76 | 1.3 |
| PE(18:0_20:3) | 14.6 | 1.1 | 16.4 | 1.66 | 0.29 | 3.6 |
| PE(18:0_22:4) | 19.5 | 0.289 | 20.6 | 0.968 | 0.21 | 2.2 |
| PE(18:0_22:5) | 17.5 | 0.73 | 18.9 | 1.01 | 0.21 | 2.5 |
| PE(18:0_22:6) | 21 | 0.702 | 22.3 | 1.12 | 0.25 | 2.4 |
| PE(18:1_20:4)/(16:0_22:5) | 20.4 | 0.413 | 21.9 | 1.04 | 0.099 | 3 |
| PE(18:1_22:4) | 16.7 | 0.838 | 17.9 | 1.15 | 0.33 | 2.3 |
| PE(18:1_22:5)/(20:4_20:2) | 14.9 | 1.49 | 16.8 | 1.04 | 0.21 | 3.7 |
| PE(18:1_22:6) | 18.4 | 0.364 | 20 | 1.11 | 0.099 | 3.2 |
| PE(18:2_18:2) | 13.4 | 0.5 | 15.1 | 1.88 | 0.33 | 3.2 |
| PE(18:2_20:3) | 16.8 | 0.846 | 18.2 | 0.961 | 0.21 | 2.7 |
| PE(18:2_20:4) | 15.7 | 0.655 | 17.4 | 1.11 | 0.11 | 3.3 |
| PE(20:4_20:1) | 18.8 | 0.368 | 19.7 | 0.751 | 0.25 | 1.8 |
| PE(34:1) | 20.1 | 0.59 | 21.2 | 0.973 | 0.25 | 2.2 |
| PE(36:1) | 21.4 | 0.361 | 22.4 | 0.963 | 0.27 | 2 |
| PE(38:2) | 20.1 | 0.286 | 21.1 | 1.03 | 0.29 | 2 |
| PE(38:3) | 16.8 | 1.08 | 18.3 | 0.955 | 0.21 | 3 |
| **Phosphatidylethanolamines-ether lipids (PE (O))** | | | | | | |
| PE(O-16:1_22:4) | 20.1 | 0.634 | 21.3 | 0.836 | 0.21 | 2.2 |
| PE(O-18:0_18:2) | 20.2 | 0.31 | 20.3 | 0.174 | 0.68 | 1.1 |
| PE(O-36:2) | 22.9 | 0.346 | 24.1 | 1.01 | 0.21 | 2.2 |
| PE(O-38:5) | 21.8 | 0.405 | 23 | 0.856 | 0.14 | 2.4 |
| **Plasmenylethalomines (PE (P))** | | | | | | |
| PE(P-18:0_22:6) | 19.9 | 0.693 | 21.3 | 0.968 | 0.21 | 2.6 |
| **Phosphatidylglycerols (PG)** | | | | | | |
| PG(34:1) | 22 | 0.202 | 22.3 | 0.115 | 0.21 | 1.2 |
| PG(34:2) | 18.8 | 0.334 | 18.8 | 0.123 | 0.95 | 1 |
| PG(36:1) | 19.6 | 0.37 | 19.6 | 0.108 | 0.98 | 0.99 |
| PG(36:2) | 19.6 | 0.272 | 19.9 | 0.151 | 0.25 | 1.2 |
| PG(36:3) | 21.3 | 0.498 | 21.3 | 0.552 | 0.98 | 0.96 |
| PG(36:4) | 19.5 | 0.939 | 19.7 | 0.643 | 0.95 | 1.2 |
| PG(38:5) | 22 | 0.532 | 21.8 | 0.655 | 0.95 | 0.9 |
| PG(38:6) | 21.3 | 0.546 | 21.3 | 0.688 | 0.99 | 1 |
| PG(40:5) | 19.8 | 0.537 | 20 | 0.554 | 0.85 | 1.2 |
| PG(40:6) | 19.1 | 0.649 | 19.1 | 0.677 | 0.98 | 1 |
| PG(40:7) | 20.2 | 0.535 | 20.4 | 0.693 | 0.95 | 1.1 |
| PG(40:8) | 21.5 | 0.504 | 21.4 | 0.712 | 0.98 | 0.96 |
| **Phosphatidylinositols (PI)** | | | | | | |

| | | | | | | |
|---|---|---|---|---|---|---|
| PI(16:0_20:4) | 20.1 | 0.601 | 20.2 | 0.743 | 0.98 | 1 |
| PI(16:0_22:6) | 17.7 | 0.825 | 17.9 | 0.775 | 0.95 | 1.2 |
| PI(18:0_18:2)/(18:1_18:1) | 20.1 | 0.186 | 20.5 | 0.423 | 0.43 | 1.3 |
| PI(18:0_20:3) | 18.3 | 0.408 | 18.3 | 0.879 | 0.99 | 0.99 |
| PI(18:0_20:4) | 24 | 0.142 | 24.2 | 0.772 | 0.93 | 1.2 |
| PI(18:0_22:4) | 18.2 | 0.175 | 18.1 | 0.788 | 0.95 | 0.92 |
| PI(18:0_22:5) | 14.2 | 1.16 | 14.3 | 1.44 | 0.98 | 1.1 |
| PI(18:0_22:6) | 18.9 | 0.229 | 19.3 | 0.749 | 0.73 | 1.3 |
| PI(18:1_20:4) | 20.6 | 0.74 | 20.6 | 0.804 | 0.98 | 1 |
| PI(20:1_20:4) | 17.4 | 0.23 | 17.8 | 0.769 | 0.75 | 1.3 |
| PI(36:3) | 17.4 | 0.416 | 17.5 | 0.869 | 0.95 | 1.1 |
| **Sphingomyelins (SM)** | | | | | | |
| SM(18:0/16:0) | 21.8 | 0.452 | 22.1 | 0.41 | 0.65 | 1.3 |
| SM(18:0/18:0) | 20.3 | 0.268 | 20.2 | 0.405 | 0.98 | 0.97 |
| SM(18:0/20:0) | 20.3 | 0.299 | 20 | 0.368 | 0.51 | 0.8 |
| SM(18:0/22:0) | 21.7 | 0.325 | 21.4 | 0.482 | 0.58 | 0.78 |
| SM(18:0/22:2)/(18:1/22:1)/(18:2/22:0) | 22 | 0.11 | 22 | 0.367 | 0.95 | 0.95 |
| SM(18:0/24:1)/(18:1/24:0) | 24.6 | 0.191 | 24.4 | 0.316 | 0.68 | 0.87 |
| SM(18:1/16:0) | 24.3 | 0.444 | 24.7 | 0.226 | 0.23 | 1.4 |
| SM(18:1/18:0) | 23.2 | 0.212 | 23.4 | 0.296 | 0.79 | 1.1 |
| SM(18:1/20:0) | 22.7 | 0.14 | 22.7 | 0.313 | 0.93 | 0.94 |
| SM(18:1/21:0) | 17.1 | 1.17 | 17.5 | 0.575 | 0.91 | 1.3 |
| SM(18:1/22:0) | 24.6 | 0.13 | 24.4 | 0.297 | 0.75 | 0.9 |
| SM(18:1/23:0) | 20.8 | 0.192 | 20.7 | 0.347 | 0.95 | 0.94 |
| SM(18:1/23:1) | 18.8 | 0.144 | 18.7 | 0.321 | 0.73 | 0.89 |
| SM(18:1/24:1)_(18:2/24:0) | 25.7 | 0.139 | 25.6 | 0.278 | 0.84 | 0.93 |
| SM(18:1/24:2)_(18:2/24:1) | 23 | 0.133 | 23 | 0.285 | 0.95 | 0.97 |
| SM(40:3) | 16.5 | 0.587 | 16.7 | 0.618 | 0.95 | 1.1 |
| **Triglycerides (TG)** | | | | | | |
| TG(48:1) | 27.8 | 0.316 | 27.1 | 0.38 | 0.099 | 0.63 |
| TG(48:2) | 27.7 | 0.433 | 27.3 | 0.439 | 0.45 | 0.74 |
| TG(50:0) | 25.3 | 0.57 | 24.5 | 0.432 | 0.21 | 0.57 |
| TG(50:1) | 29.1 | 0.402 | 28.5 | 0.375 | 0.21 | 0.66 |
| TG(50:2) | 29.6 | 0.298 | 29.1 | 0.429 | 0.21 | 0.68 |
| TG(50:3) | 28.8 | 0.435 | 28.4 | 0.511 | 0.57 | 0.75 |
| TG(50:4) | 26.8 | 0.555 | 26.4 | 0.662 | 0.79 | 0.79 |
| TG(52:1) | 27.9 | 0.667 | 27.1 | 0.566 | 0.25 | 0.57 |
| TG(52:2) | 30.2 | 0.384 | 29.7 | 0.421 | 0.29 | 0.71 |
| TG(52:3) | 30.3 | 0.305 | 29.8 | 0.456 | 0.29 | 0.71 |
| TG(52:4) | 29.3 | 0.425 | 28.8 | 0.536 | 0.47 | 0.72 |
| TG(52:5) | 27 | 0.524 | 26.6 | 0.638 | 0.65 | 0.73 |
| TG(54:2) | 28.4 | 0.575 | 27.7 | 0.621 | 0.29 | 0.59 |
| TG(54:3) | 30 | 0.387 | 29.5 | 0.476 | 0.33 | 0.71 |

| | | | | | | |
|---|---|---|---|---|---|---|
| TG(54:4) | 29.8 | 0.34 | 29.3 | 0.485 | 0.33 | 0.71 |
| TG(54:5) | 28.8 | 0.453 | 28.2 | 0.565 | 0.36 | 0.67 |
| TG(54:6) | 27.1 | 0.489 | 26.6 | 0.532 | 0.51 | 0.71 |
| TG(56:6) | 26.4 | 0.478 | 25.8 | 0.642 | 0.45 | 0.67 |
| TG(58:10) | 21.3 | 0.519 | 20.9 | 0.531 | 0.75 | 0.8 |
| TG(58:6) | 23.8 | 0.4 | 23.4 | 0.606 | 0.65 | 0.76 |
| TG(58:8) | 24.2 | 0.509 | 23.7 | 0.656 | 0.59 | 0.71 |
| TG(60:10) | 20.5 | 0.485 | 20.2 | 0.546 | 0.68 | 0.77 |
| TG(60:11) | 19.1 | 2.17 | 19.3 | 1.93 | 0.98 | 1.2 |

**Table S6. Primers, sgRNA and ssDNA used in this study.**

| Primer | Application | Sequence |
|---|---|---|
| Cxcl14-F | qRT-PCR | 5'-TACCCACACTGCGAGGAGAA-3' |
| Cxcl14-R | qRT-PCR | 5'-CGTTCCAGGCATTGTACCACT-3' |
| LepR-F | qRT-PCR | 5'-TGATGTGTCAGAAATTCTATGTG-3' |
| LepR-R | qRT-PCR | 5'-TGCCAGGTTAAGTGCAGCTAT-3' |
| CD36-F | qRT-PCR | 5'-GGACATTGAGATTCTTTTCCTCTG-3' |
| CD36-R | qRT-PCR | 5'-GCAAAGGCATTGGCTGGAAGAAC-3' |
| Fabp4-F | qRT-PCR | 5'-TGAAATCACCGCAGACGACAGG-3' |
| Fabp4-R | qRT-PCR | 5'-GCTTGTCACCATCTCGTTTTCTC-3' |
| Actinb-F | qRT-PCR | 5'-GCTCTTTTCCAGCCTTCCTT-3' |
| Actinb-R | qRT-PCR | 5'-CTTCTGCATCCTGTCAGCAA-3' |
| Cre-F | Genotyping | 5'-GCGGTCTGGCAGTAAAAACTATC-3' |
| Cre-R | Genotyping | 5'- GTGAAACAGCATTGCTGTCACTT-3' |
| Scf-GFP-F | Genotyping | 5'- CCCGCAGCTCTGGTATATTTGC-3' |
| Scf-GFP-R1 | Genotyping | 5'- CGGACACGCTGAACTTGTGG-3' |
| Scf-GFP-R2 | Genotyping | 5'- AAGCACTTCAGATTCTAGGG-3' |
| Cxcl14 full length F | Genotyping | 5'-GAGAGAGGCGTGCTTGAAA-3' |
| Cxcl14 full length R | Genotyping | 5'-GCAGAAGAGACAGGAGCATTC-3' |
| Cxcl14 flox-F | Genotyping | 5'-GAGAGAGGCGTGCTTGAAA-3' |
| Cxcl14 flox-R | Genotyping | 5'-TTGGACCCTGCGGAAGA-3' |
| Cxcl14 dsRed full length-F | Genotyping | 5'-CCTGCTGGCATTCTCACTT-3' |
| Cxcl14 dsRed full length-R | Genotyping | 5'-GTGGATCTGGTGAAGATGGATG-3' |
| Cxcl14 dsRed-KI-F | Genotyping | 5'-GCTTGAAACCGAGAACC-3' |
| Cxcl14 dsRed-KI-R | Genotyping | 5'-ACACTTACACTTGGACCCT-3' |
| Cxcl14 5'gRNA | sgRNA | CAAGCACCACCTAGGCATCCTGG (Sense strand) |
| Cxcl14 3'gRNA | sgRNA | TGCCACTTTGAATCCTAACATCCAGG (AntiSense strand) |
| Cxcl14 dsRed-KI gRNA | sgRNA | GCGCCCCTCCGGCCAGCATG AGG (sense strand) |

| | | |
|---|---|---|
| Cxcl14 exon2 ssDNA | Donor oligo | GGCGTGCTTGAAACCGAGAACCAAGCCGGGCGGCA TCCCCCGGCCGCCGCACGCACAGGCCGGCGCCCT CCTTGCCTCCCTGCTCCCCACCGCGCCCCTCCGGC CAGCATGAGGCTCCTGGCGGCCGCGCTGCTCCTGC TGCTCCTGGCGCTGTGCGCCTCGCGCGTGGACGGT GAGTGCCGCGAGGGCCCTCTGTCGCGGTCCTGCCC CGTCCTAGGGACCCCAAGCACCACCTAGGCAGACG TCATAACTTCGTATAATGTATGCTATACGAAGTTATTC CTGGAACCCGGGGCGGGGGGTGGGGGGGTGGATC CCGGGCGGACACCTTGGGCGGTCGCGGAGACCTGT ACGGCGGCGACTCAGCAGCCCGGCCGCGCTCTTCC GCAGGGTCCAAGTGTAAGTGTTCCCGGAAGGGGCC CAAGATCCGCTACAGCGACGTGAAGAAGCTGGAAAT GAAGCCAAAGTACCCACACTGCGAGGAGAAGATGG TTATGTGAGTTCTGGGGTAGCATATCCTGTGGAGGA GCCCAACCTAGGGTTAGATCGGACTGATAACAGCCT TTCTCTCTGCCTTTGACAAGACCGACTGGCAAGTCC TGGGGACATTGCCTGAGAGTCTGCGGAAAGGGGAT GAGGGTGTCTTTGGCCGGTCCCTCTGAGCCTGTCCT TGTGATATCATAACTTCGTATAATGTATGCTATACGA AGTTATTAGGATTCAAAGTGGCAGGCGAGGTTTATT GTAGGCTAGTTAGTGGTTCTGGTGCGCCCGCTTAGA ACATCCATCTTCACCAGATCCACTCGGGACACGCAC CCGGCCTGTTGGGCCGGCTGGGCTGCCTTCCTCTT ACTGAAGAGAGAAGTTAACTTTTCC |
| Cxcl14-DsRed-PolyA ssDNA | Donor oligo | CCAAGGCACTGAGGTATGGGTGTGCAAGCGTAAAG GCGGGCGTGCTGTGGAGCGAGAGGGTAGCGGATGT GAGTGTGTCCGTGTGTGCGCGCGTGGCTCCGAGTG TGCGCCGCTGGGATCGCGATGGGTCCGGGGCGGG CGGCAGGGCGGTCTGGGAGATCTCCTCCCCCCACC ACATTGAGAAATCTCAGTGAGTCACCGAGTGGTTCT GCATATTAATGAGCTCGCTCTCCGAGAGGGCAGGAG CGAATTTAAAAGAGGCCAGGGTGGGCGGAGGGAAG CTGTGGGGAGATCGCAGCACCCAGCGCCAAGCGCA GCGCGGCACCGCGACAGACGGCAGGAGCACCCATC GACGGGCGTAGCTGGAGCGAGCCGAGCAGAGCAG AGAGAGGCGTGCTTGAAACCGAGAACCAAGCCGGG CGGCATCCCCCGGCCGCCGCACGCACAGGCCGGC GCCCTCCTTGCCTCCCTGCTCCCCACCGCGCCCCT CCGGCCAGCATGATGGATAGCACTGAGAACGTCATC AAGCCCTTCATGCGCTTCAAGGTGCACATGGAGGGC TCCGTGAACGGCCACGAGTTCGAGATCGAGGGCGA GGGCGAGGGCAAGCCCTACGAGGGCACCCAGACC GCCAAGCTGCAGGTGACCAAGGGCGGCCCCCTGCC CTTCGCCTGGGACATCCTGTCCCCCCAGTTCCAGTA CGGCTCCAAGGTGTACGTGAAGCACCCCGCCGACA TCCCCGACTACAAGAAGCTGTCCTTCCCCGAGGGCT TCAAGTGGGAGCGCGTGATGAACTTCGAGGACGGC GGCGTGGTGACCGTGACCCAGGACTCCTCCCTGCA GGACGGCACCTTCATCTACCACGTGAAGTTCATCGG CGTGAACTTCCCCTCCGACGGCCCCGTAATGCAGAA |

| | | |
|---|---|---|
| | | GAAGACTCTGGGCTGGGAGCCCTCCACCGAGCGCC TGTACCCCCGCGACGGCGTGCTGAAGGGCGAGATC CACAAGGCGCTGAAGCTGAAGGGCGGCGGCCACTA CCTGGTGGAGTTCAAGTCAATCTACATGGCCAAGAA GCCCGTGAAGCTGCCCGGCTACTACTACGTGGACT CCAAGCTGGACATCACCTCCCACAACGAGGACTACA CCGTGGTGGAGCAGTACGAGCGCGCCGAGGCCCG CCACCACCTGTTCCAGTAGGCGGCCGCGACTCTAG ATCATAATCAGCCATACCACATTTGTAGAGGTTTTAC TTGCTTTAAAAAACCTCCCACACCTCCCCCTGAACCT GAAACATAAAATGAATGCAATTGTTGTTGTTAACTTG TTTATTGCAGCTTATAATGGTTACAAATAAAGCAATA GCATCACAAATTTCACAAATAAAGCATTTTTTTCACTG CATTCTAGTTGTGGTTTGTCCAAACTCATCAATGTAT CTTAAGGCTCCTGGCGGCCGCGCTGCTCCTGCTGC TCCTGGCGCTGTGCGCCTCGCGCGTGGACGGTGAG TGCCGCGAGGGCCCTCTGTCGCGGTCCTGCCCCGT CCTAGGGACCCCAAGCACCACCTAGGCATCCTGGA ACCCGGGGCGGGGGGTGGGGGGGTGGATCCCGGG CGGACACCTTGGGCGGTCGCGGAGACCTGTACGGC GGCGACTCAGCAGCCCGGCCGCGCTCTTCCGCAGG GTCCAAGTGTAAGTGTTCCCGGAAGGGGCCCAAGAT CCGCTACAGCGACGTGAAGAAGCTGGAAATGAAGC CAAAGTACCCACACTGCGAGGAGAAGATGGTTATGT GAGTTCTGGGGTAGCATATCCTGTGGAGGAGCCCAA CCTAGGGTTAGATCGGACTGATAACAGCCTTTCTCT CTGCCTTTGACAAGACCGACTGGCAAGTCCTGGGGA CATTGCCTGAGAGTCTGCGGAAAGGGGATGAGGGT GTCTTTGG |

**Movie S1: *Cxcl14*-dsRed⁺ stromal cells extend fine processes that wrap around megakaryocytes.** Representative video of a megakaryocyte (anti-CD41 antibody staining in magenta) in femur bone marrow that was closely associated with *Cxcl14*-dsRed⁺ stromal cells (yellow) that extend find processes that wrap around the megakaryocyte.
